# Validation of a high-performance liquid chromatography-tandem mass spectrometry immunopeptidomics assay for the identification of HLA class I ligands suitable for pharmaceutical therapies

**DOI:** 10.1101/821249

**Authors:** Michael Ghosh, Marion Gauger, Ana Marcu, Annika Nelde, Monika Denk, Heiko Schuster, Hans-Georg Rammensee, Stefan Stevanović

## Abstract

For more than two decades naturally presented, human leukocyte antigen (HLA)-restricted peptides (immunopeptidome) have been eluted and sequenced using liquid chromatography-tandem mass spectrometry (LC-MS/MS). Since, identified disease-associated HLA ligands have been characterized and evaluated as potential active substances. Treatments based on HLA-presented peptides have shown promising results in clinical application as personalized T cell-based immunotherapy. Peptide vaccination cocktails are produced as investigational medicinal products under GMP conditions. In order to support clinical trials based on HLA-presented tumor-associated antigens, in this study the sensitive LC-MS/MS HLA class I antigen identification pipeline was fully validated for our technical equipment according to the current US Food and Drug Administration (FDA) and European Medicines Agency (EMA) guidelines.

The immunopeptidomes of JY cells with or without spiked-in, isotope labeled peptides, of peripheral blood mononuclear cells of healthy volunteers as well as a chronic lymphocytic leukemia and a bladder cancer sample were reliably identified using a data-dependent acquisition method. As the LC-MS/MS pipeline is used for identification purposes, the validation parameters include accuracy, precision, specificity, limit of detection and robustness.

## Introduction

The immunopeptidome is a vast and diverse compilation of HLA-presented peptides (HLA ligands), which serve as a showcase of inter- and intracellular processes. T cells recognize presented peptides in the immunopeptidome, which is constantly modulated by gene expression, transcription, translation, posttranslational modification, and antigen processing and presentation^1–4^. Especially in tumor immunology, HLA ligands are used in many ways. They are suited as biomarkers, presenting intracellular abnormalities like malignant transformation and as active pharmaceuticals, activating cancer specific T cells^5^.

Natural HLA ligands have been isolated and sequenced using LC-MS/MS for almost three decades^6–12^. So far, the LC-MS/MS analysis is the only method to investigate the entirety of HLA presented peptides. However, based on these peptide data *in silico* prediction tools have been developed, which allow the prediction of possibly presented peptides from exome, RNA or whole genome sequencing data and have extended the toolbox even further^13–18^.

Developing from such identifications, peptide vaccination cocktails have been produced as active pharmaceuticals under GMP conditions^19^. The acceptance, safety and efficacy of peptide vaccinations has been investigated^19^ and several clinical studies testing peptide vaccinations have been performed with our contribution (GAPVAC^5^, NCT02149225 and NOA-16, NCT02454634) or are ongoing (iVAC-CLL01, NCT02802943). The procedures of active substance production, analysis, and batch release have been validated and reliably lead to reproducible products of desired quality. However, the initial antigen identification procedure using mass spectrometry-based immunopeptidomics has not been validated yet. In this study, the LC-MS/MS antigen identification procedure was fully validated for our technical equipment according to current FDA and EMA guidelines to support further clinical trials based on HLA-presented tumor-associated antigens. This validation should serve as a guidance that can be adapted to other LC-MS/MS platforms and samples.

Protocols for large-scale immunopeptidomics using LC-MS/MS and the identification of HLA ligands are established and have been published^20–22^. To our knowledge, no validation of an omics method using LC-MS/MS-has been published so far^23,24^. This article presents an immunopeptidomics assay using LC-MS/MS, which is fully validated according to the latest US Food and Drug Administration (FDA) and European Medicines Agency (EMA) guidelines^23–27^. We have to emphasize that such validations are specific only for dedicated equipment of one distinct laboratory. We provide a first protocol and template to enhance the validation of other laboratories with similar equipment and other omics fields using LC-MS/MS such as proteomics, metabolomics, and lipidomics.

## Experimental procedures

### Peptide synthesis

The automated peptide synthesizer Liberty Blue (CEM) was used to synthesize peptides following the 9-fluorenylmethyl-oxycarbonyl/tert-butyl (Fmoc/tBu) strategy. The identity and purity of the peptides were confirmed using a reversed-phase liquid chromatography (Alliance e2965, Waters) and an uHPLC system (nanoUHPLC, UltiMate 3000 RSLCnano, Dionex) on-line coupled LTQ Orbitrap XL hybrid mass spectrometer (ThermoFisher) system. Synthesized peptides were employed in the validation of LC-MS/MS identifications. Peptide sequences used for the validation are listed in supplemental Table S1.

### Tissue samples

The EBV-transformed human B-cell line JY (ECACC 94022533) was cultured in RPMI1640 with 10% heat-inactivated fetal bovine serum (FBS) and 1% penicillin/streptomycin to a total number of 1*10^11^ cells, centrifuged at 1,500 rpm for 15 min at 4°C, washed two times with cold PBS and aliquots containing 75*10^6^ cells were frozen and stored at −80°C until use. The cells were tested negative for mycoplasma contamination via PCR.

The peripheral blood mononuclear cells (PBMC), chronic lymphocytic leukemia (CLL) and bladder cancer (BC) tissue samples were collected at the University Hospital of Tübingen with the informed consent of patients according to the principles of the Declaration of Helsinki. The local institutional review board (Ethics Committee at the Medical Faculty and the University Hospital of Tübingen) has approved the use of the patient samples.

### Immunoaffinity purification of HLA ligands

HLA class I molecules were isolated using standard immunoaffinity purification as described^9,20,28,29^ using the HLA class I-specific monoclonal antibody W6/32^30^. First, the cell pellets were lysed in 10 mM CHAPS (Applichem)/PBS (Lonza) containing protease inhibitors (Complete, Roche) and subsequently HLA molecules were purified using the pan-HLA class I– specific monoclonal W6/32 Ab covalently linked to CNBr-activated Sepharose (GE Healthcare). Repeated addition of 0.2% trifluoroacetic acid (Merck) eluted HLA molecules and peptides. The peptides were isolated employing ultrafiltration with centrifugal filter units (Amicon, Merck Millipore), extracted and desalted using ZipTip C18 pipette tips (Merck Milli-pore), eluted in 35 µl acetonitrile (Merck)/0.1% trifluoroacetic acid, vacuum centrifuged to 5 µl, and resuspended in 25 µl of 1% acetonitrile/0.05% trifluoroacetic acid. Finally, the peptide solutions were stored at −20°C until analysis by LC-MS/MS.

### Analysis of HLA ligands by LC-MS/MS

Peptides were separated by nanoflow high-performance liquid chromatography (nanoUHPLC, UltiMate 3000 RSLCnano, Dionex) and subsequently analyzed in an on-line coupled Orbitrap Fusion Lumos or LTQ Orbitrap XL mass spectrometer (Thermo Fisher Scientific). Volumes of 5 μl peptide solution were injected onto a 75 μm × 2 cm trapping column (Acclaim PepMap RSLC, Dionex) at 4 μl/min for 5.75 min in five technical replicates. Subsequently, peptide separation was performed at 50°C at a flow rate of 300 nl/min on a 50 μm × 25 cm separation column (Acclaim PepMap RSLC, Dionex) applying a gradient ranging from 2.4 to 32.0% of AcN over the course of 90 min. Eluted peptides were ionized by nanospray ionization and analyzed in the Orbitrap Fusion Lumos implementing top speed collision-induced dissociation (CID) fragmentation. Survey scans were performed at 120,000 resolution and fragment detection at 60,000 resolution in the Orbitrap.

To demonstrate a method transfer, the immunopeptidomics pipeline was transferred from the Orbitrap Fusion Lumos to a LTQ Orbitrap XL. In the LTQ Orbitrap XL peptides were analyzed using a top five CID method with survey scans at 60,000 resolution and fragment ion detection in the ion trap operated at normal scan speed. On both instruments, the mass range was limited to 400–650 m/z with precursors of charge states 2+ and 3+ eligible for fragmentation.

Maintenance and OQ of the LC-MS/MS system are performed annually (Thermo Fisher Scientific). A positive ion calibration using a Pierce™ LTQ velos or LTQ ESI positive ion calibration solution (Thermo Fisher Scientific) and a system suitability test using natural HLA class I-presented peptides of JY cells is performed weekly.

### Database search and spectral annotation

Data was processed against the human proteome included in the Swiss-Prot database (http://www.uniprot.org, release September 27, 2013; containing 20,279 reviewed protein sequences) applying the Sequest algorithm^31^ in the Proteome Discoverer (version 1.3, Thermo Fisher) software.

Precursor mass tolerance was set to 5 ppm, product ions mass tolerance was set to 0.02 Da for Orbitrap Fusion Lumos data and 0.5 Da for LTQ Orbitrap XL Data and oxidized methionine was allowed as the only dynamic modification with no restriction by enzymatic specificity. Percolator^32^-assisted false discovery rate (FDR) calculation was set at a target value of q ≤ 0.05 (5% FDR). Peptide-spectrum matches with q ≤ 0.05 were filtered according to additional orthogonal parameters to ensure spectral quality and validity. Peptide lengths were limited to 8– 12 amino acids.

### Validation procedures

The validation of the immunopeptidomics procedure was done according to the OECD principles of Good Laboratory Practice (GLP)^33^ and accuracy, precision, specificity, limit of detection and robustness were validated according to the FDA and EMA guidelines^25,26^. Clear definitions can be found in^25,26,33^.

As the immunopeptidome LC-MS/MS system separates the peptides using the LC and subsequently identifies the peptide ions in MS mode and product ions in the MS/MS mode, we tried to consider these three parts for every validation parameter summarized in Table 1 and Figure 1.

**Table 1:** Acceptance criteria for the different parameters selected for LC-MS/MS validation. The acceptance criteria for the selected parameters are indicated for the mass spectrometer and the LC. Abbreviations: PC, Pearson correlation; SD, standard deviation.

|  | MS and MS/MS | LC |
| --- | --- | --- |
| Characteristics | Specification/ acceptance criteria |  |
| **Accuracy**<br>(comparison to theoretical masses and synthetic peptides) | The median of the deviation of the theoretical masses ($\Delta M$ ppm) of<br>the entirety of peptides<br>the selected peptides between natural and synthetic peptides should be $\leq 2$ ppm | PC $\geq 95\%$ |
| **Precision**<br>(natural peptides)<br>repeatability<br>intermediate precision | SD of peptide number: $\leq 10\%$<br>Recovery rate: $80\% \pm 20\%$ | PC $\geq 95\%$ |
| **Specificity**<br>(natural and synthetic peptides) | The SD between the precursor ion and five top fragment masses of selected peptides: $\leq 0.001$ Da<br>Selectivity of all identified peptides based on precursor mass combined with top five fragments | PC $\geq 95\%$ |
| **Limit of detection**<br>(synthetic peptides) | 50% ($n = 31$) of the peptides have to be identified<br>Recovery rate: $80\% \pm 20\%$ | PC $\geq 95\%$ |
| **Robustness**<br>(natural peptides from three primary samples isolated by three different persons) | **Accuracy:** as mentioned above<br>**Precision:**<br>repeatability: as mentioned above<br>**Specificity:** as mentioned above | PC $\geq 95\%$ |

**Figure 1:**
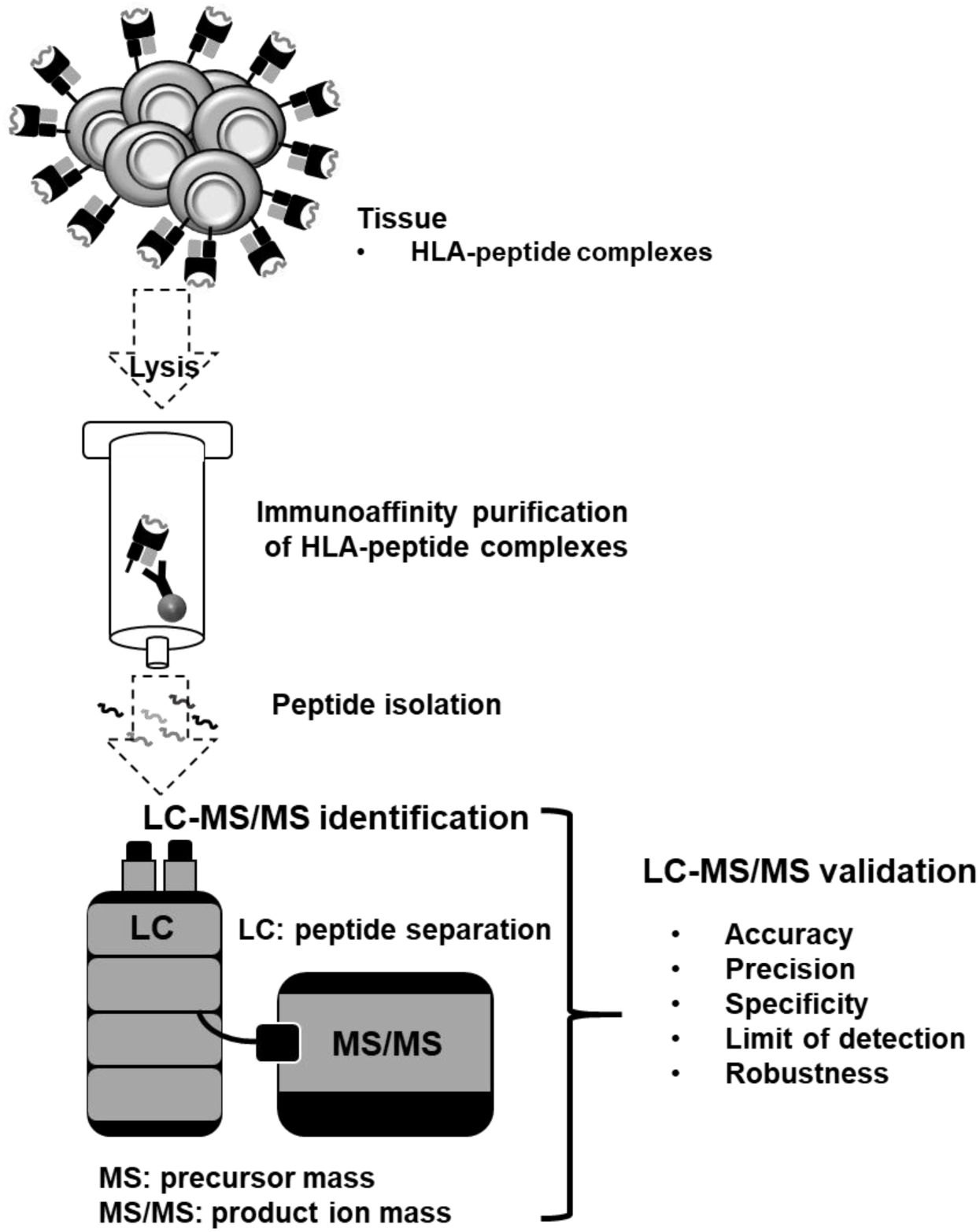
Schematic overview of the validation of the LC-MS/MS immunopeptidomics assay for the identification of HLA ligands suitable for pharmaceutical therapies. The LC-MS/MS pipeline is used for identification purposes, consequently the validation parameters accuracy, precision, specificity, limit of detection and robustness were validated according to current FDA and EMA guidelines.

### Experimental design and statistical rationale

A summary of the performed experiments, samples, technical replicates and MS RAW files is given in supplemental Table S2. Results were analyzed using GraphPad Prism (GraphPad software Inc). The recovery rate was obtained by taking the average of the percentual overlapping peptides between the technical replicates normalized to the total number of peptides. The LC peptide retention times (RTs) were compared calculating the average of the Pearson correlations of the technical replicates.

## Results

### Accuracy

To investigate the accuracy and specificity, the purified HLA-eluted peptides from one JY batch was spiked with 100 fmol isotope labeled synthetic peptides (Table S1) and analyzed in three separate analytical replicates (for identified peptides, see supplemental Table S3). The accuracy of the mass spectrometer did fulfill the acceptance criteria (Table 1) with a deviation below 1 ppm between the median mass deviation from the theoretical mass of all identified natural (median ΔM: 0.02 ppm) and synthetic peptides (median ΔM: 0.32 ppm) in Figure 2A. The peptide RTs between the replicates of all natural and all synthetic peptides do have a mean Pearson correlation above 95%, verifying the accuracy of the LC (Figure 2B).

**Figure 2:**
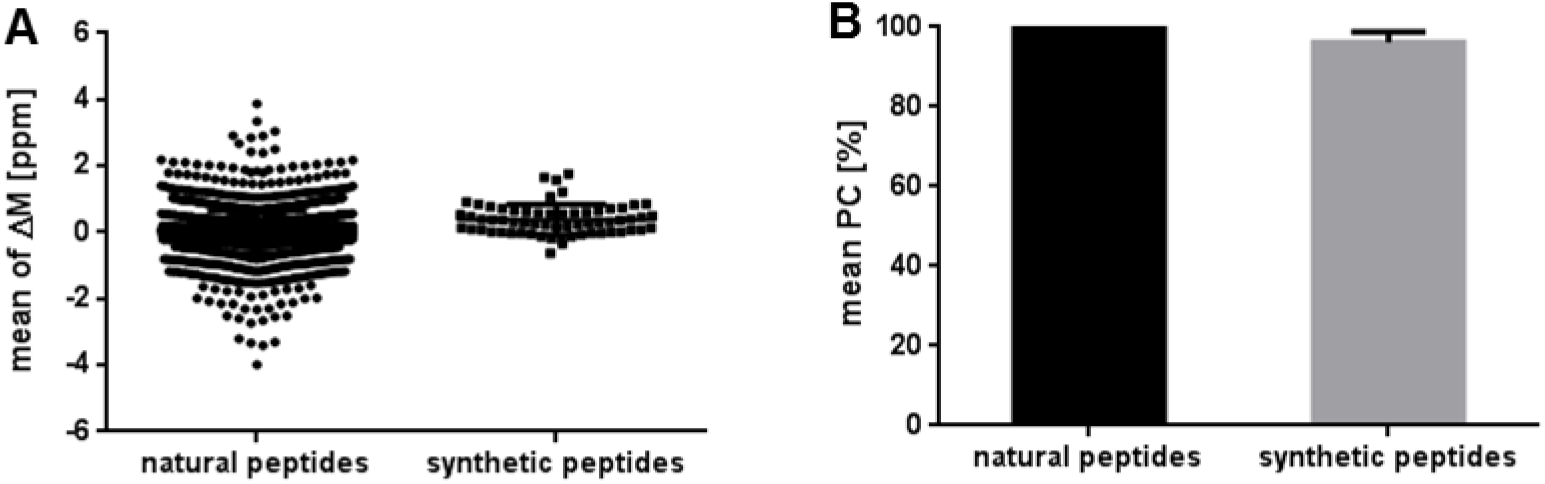
Validation of the accuracy using immunopeptidomes from JY cells and spiked isotope labeled synthetic peptides. Three replicates were analyzed. (A) Mass deviation of the detected precursor mass from the theoretical mass (ΔM ppm) of all identified natural (n = 2330) and synthetic peptides (n = 62). (B) Mean Pearson correlation of the peptide retention times. Abbreviations: PC, Pearson correlation; ppm, parts per million; ΔM, mass deviation.

### Specificity

Based on our experience from the first series of analyses, the six peptides AIVDKVPSV, SPQGRVMTI, RPSGPGPEL, YLLPAIVHI, KVLEYVIKV, and SPSSILSTL are expected as natural HLA class I-presented peptides of JY cells. In order to prove the specificity, the mass spectrometer must fulfill the MS mode acceptance criteria for precursor ions and the MS/MS mode acceptance criteria for five selected top product ions of the expected six peptides (Table 1). Here, we use two ways to select the five top product ions, we simply choose the top five most intensive fragments (last paragraph of specificity) or the most intensive fragments such as b and y fragments with the highest intensity and relevance (penultimate paragraph of specificity). Furthermore, in the LC separation, the correlation of the retention times of the natural and synthetic counterparts of the six peptides have to fulfill the acceptance criteria.

The difference of the median of the mass deviation from the theoretical mass (ΔM ppm) of the six selected peptides AIVDKVPSV, SPQGRVMTI, RPSGPGPEL, YLLPAIVHI, KVLEYVIKV, and SPSSILSTL, which were identified as natural (median ΔM: 0.17 ppm) and synthetic peptides (median ΔM: 0.21 ppm), is below 1 ppm (Figure 3A) (for identified peptides and product ions, see supplemental Table S4).

**Figure 3:**
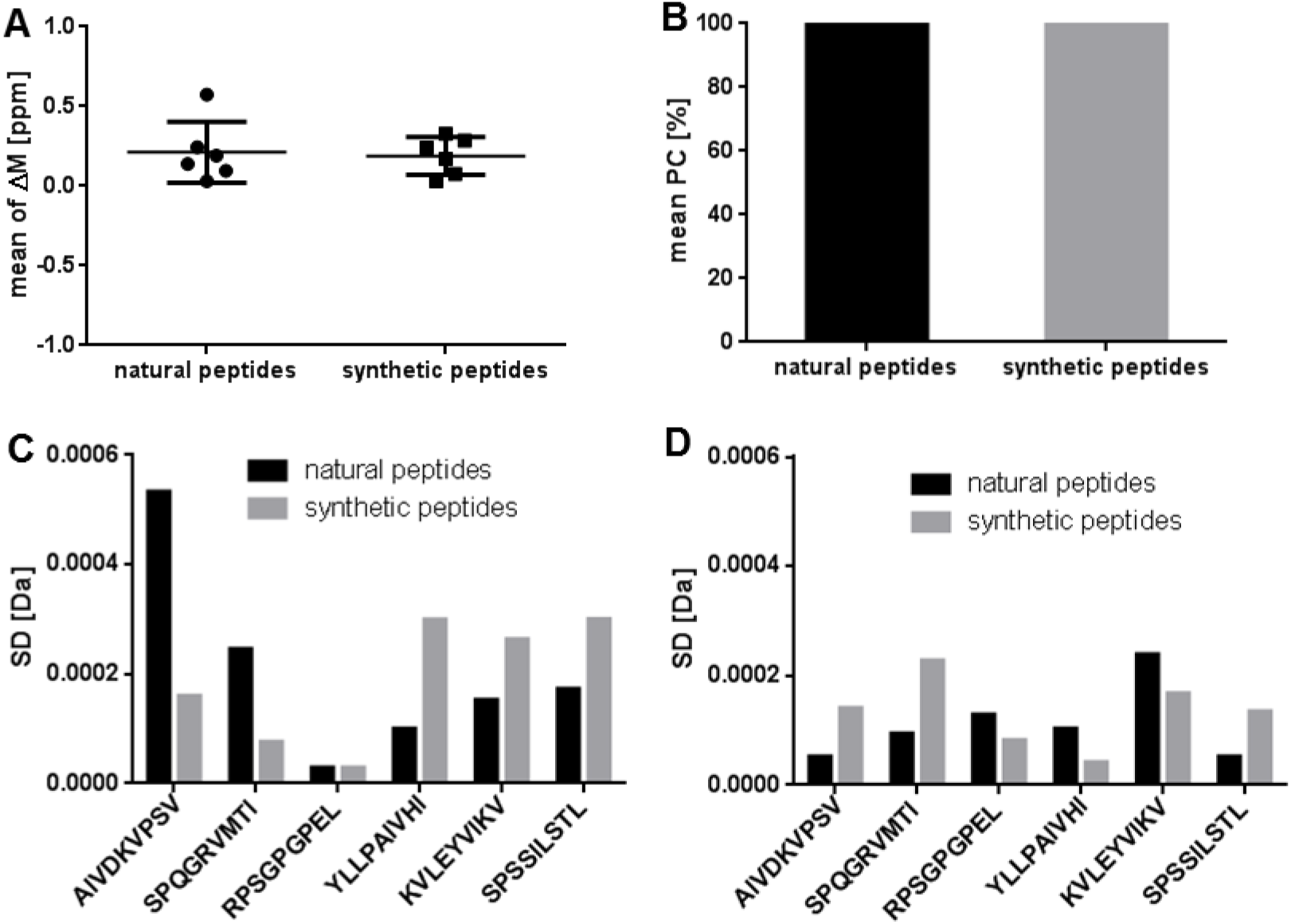
Validation of the specificity using immunopeptidomes from JY cells and spiked isotope labeled synthetic peptides. Three replicates were analyzed. (A) Mean of the mass deviation from the theoretical precursor mass (ΔM ppm) of the six identified natural and synthetic peptides AIVDKVPSV, SPQGRVMTI, RPSGPGPEL, YLLPAIVHI, KVLEYVIKV, and SPSSILSTL in three technical replicates. (B) Mean Pearson correlation of the RTs of the six identified natural and synthetic peptides. Mass deviation as SD of the (C) precursor ion masses in MS mode and the (D) resulting five selected top fragments in MS/MS modes of the six identified natural and synthetic peptides. Abbreviations: PC, Pearson correlation; ppm, parts per million; ΔM, mass deviation; SD, standard deviation.

The peptide RTs between the replicates of the six selected peptides AIVDKVPSV, SPQGRVMTI, RPSGPGPEL, YLLPAIVHI, KVLEYVIKV, and SPSSILSTL, which were identified as natural and synthetic peptides, do have a Pearson correlation above 95% (Figure 3B).

The standard deviation (SD) of the mass accuracy of the precursor masses in MS mode (Figure 3C) and of the five selected top product ions, selected based on intensity and relevance, in MS/MS mode (Figure 3D, for MS/MS spectra, see supplemental Figure S1) of the six selected peptides AIVDKVPSV, SPQGRVMTI, RPSGPGPEL, YLLPAIVHI, KVLEYVIKV and SPSSILSTL is below 0.001 Da for both the natural and synthetic peptides.

As an additional step to prove the specificity of the LC-MS/MS system, we validated our manual quality control method, to distinguish all peptides based on the mass of a precursor ion combined with the masses of the top five most intensive product ions. As the SD of the mass accuracy deviates at four decimals, our peptide identity criteria of the quality control should enable specificity at a four-decimal level (Table 1). Based on the precursor masses in MS mode at four-digit level there was an overlap of 21 to 23 natural peptide masses in the tree replicates (Table 2A). These peptide masses could be separated using the top five masses with the highest intensity in MS/MS mode. In MS/MS mode the highest number of overlapping product ion masses was three (Table 2B). All 62 synthetic peptides could be separated based on the precursor masses at four decimals (Table 2C).

**Table 2:**
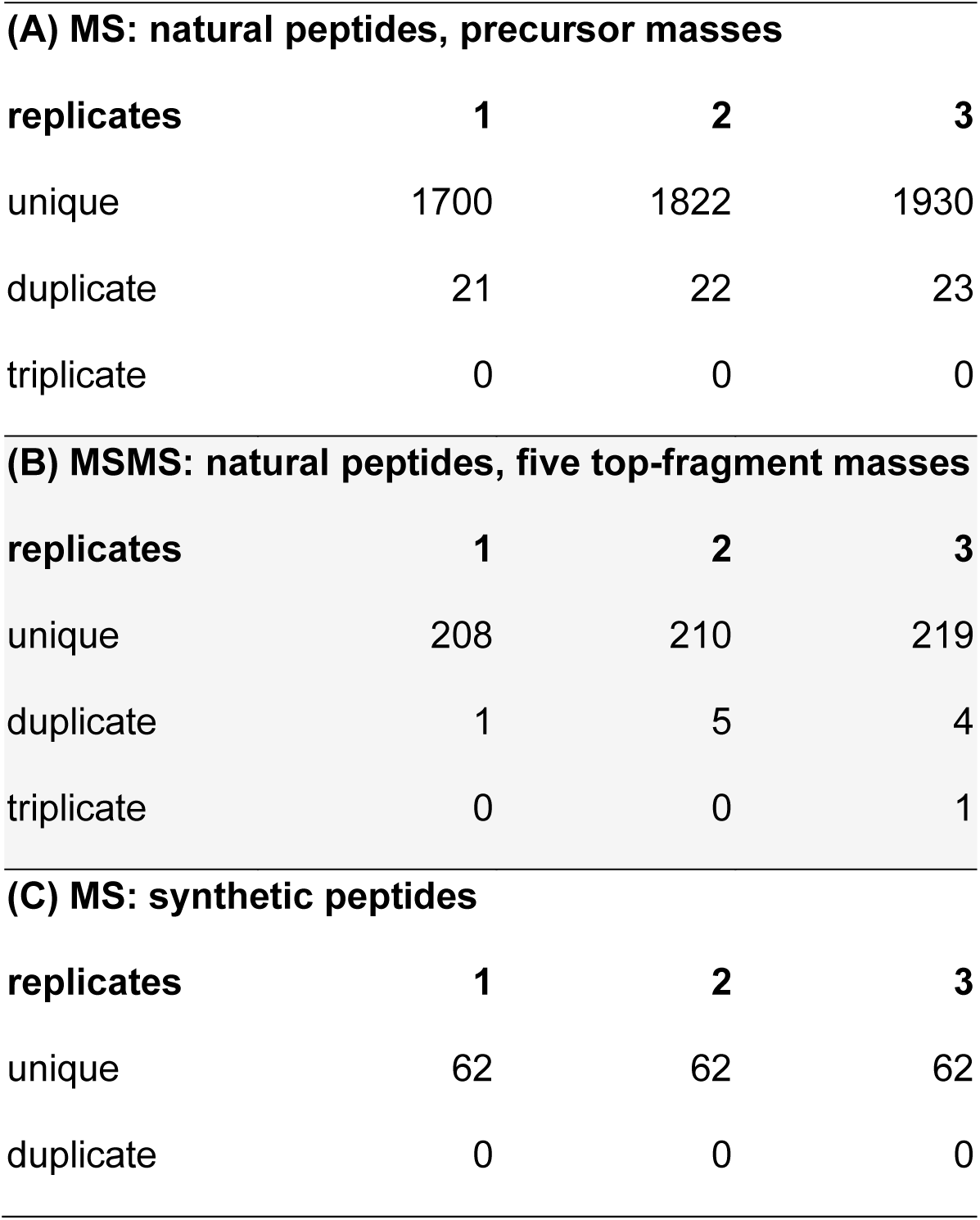
Validation of the specificity and suitability of the top five product ion peptide quality control using immunopeptidomes from JY cells and spiked isotope labeled synthetic peptides. (A) Overlap of the detected peptide precursor masses in MS mode of the natural peptides at four decimals. (B) Overlap of the measured top five product ion masses of the manifold peptide precursor masses from (A) in MS/MS mode of the natural peptides at four decimals. (C) Overlap of the measured 62 synthetic peptide precursor masses in MS mode at four decimals.

| **(A) MS: natural peptides, precursor masses** |  |  |  |
| --- | --- | --- | --- |
| **replicates** | **1** | **2** | **3** |
| unique | 1700 | 1822 | 1930 |
| duplicate | 21 | 22 | 23 |
| triplicate | 0 | 0 | 0 |
| **(B) MSMS: natural peptides, five top-fragment masses** |  |  |  |
| **replicates** | **1** | **2** | **3** |
| unique | 208 | 210 | 219 |
| duplicate | 1 | 5 | 4 |
| triplicate | 0 | 0 | 1 |
| **(C) MS: synthetic peptides** |  |  |  |
| **replicates** | **1** | **2** | **3** |
| unique | 62 | 62 | 62 |
| duplicate | 0 | 0 | 0 |

### Limit of Detection

To determine the limit of detection (LOD), four aliquots of purified HLA-eluted peptides from JY cells were spiked with 0.1 fmol, 1 fmol, 10 fmol, or 100 fmol isotope labeled synthetic peptides (Table S1) and analyzed in three replicates leading to 12 separate analytical replicates (for identified peptides, see supplemental Table S3). Based on our experience with JY cells, in HLA ligandomic experiments an optimal setting enables a peptide recovery rate of 80 ± 20% between two replicates. Thus, there is no LOD where 100% of the peptides will be discovered, especially in data-dependent acquisition. Here, we set the LOD to the peptide concentration that enables an identification of at least 50% (n = 31) of the peptides per replicate and with a recovery rate of 80% ± 20% and a Pearson correlation of the peptide retention times above 95% between three replicates.

The JY sample spiked with 1 fmol synthetic peptides had the lowest peptide content enabling a reproducible identification of 50% of the 62 added isotope labeled peptides (Figure 4A). At the LOD the recovery rate of peptides in a replicate mass spectrometric measurement is in the range of 80% ± 20% (Figure 4B) and the mean Pearson correlation of the retention times of the synthetic peptides in the three technical replicates analyzed in the LC is above 95% (Figure 4C).

**Figure 4:**
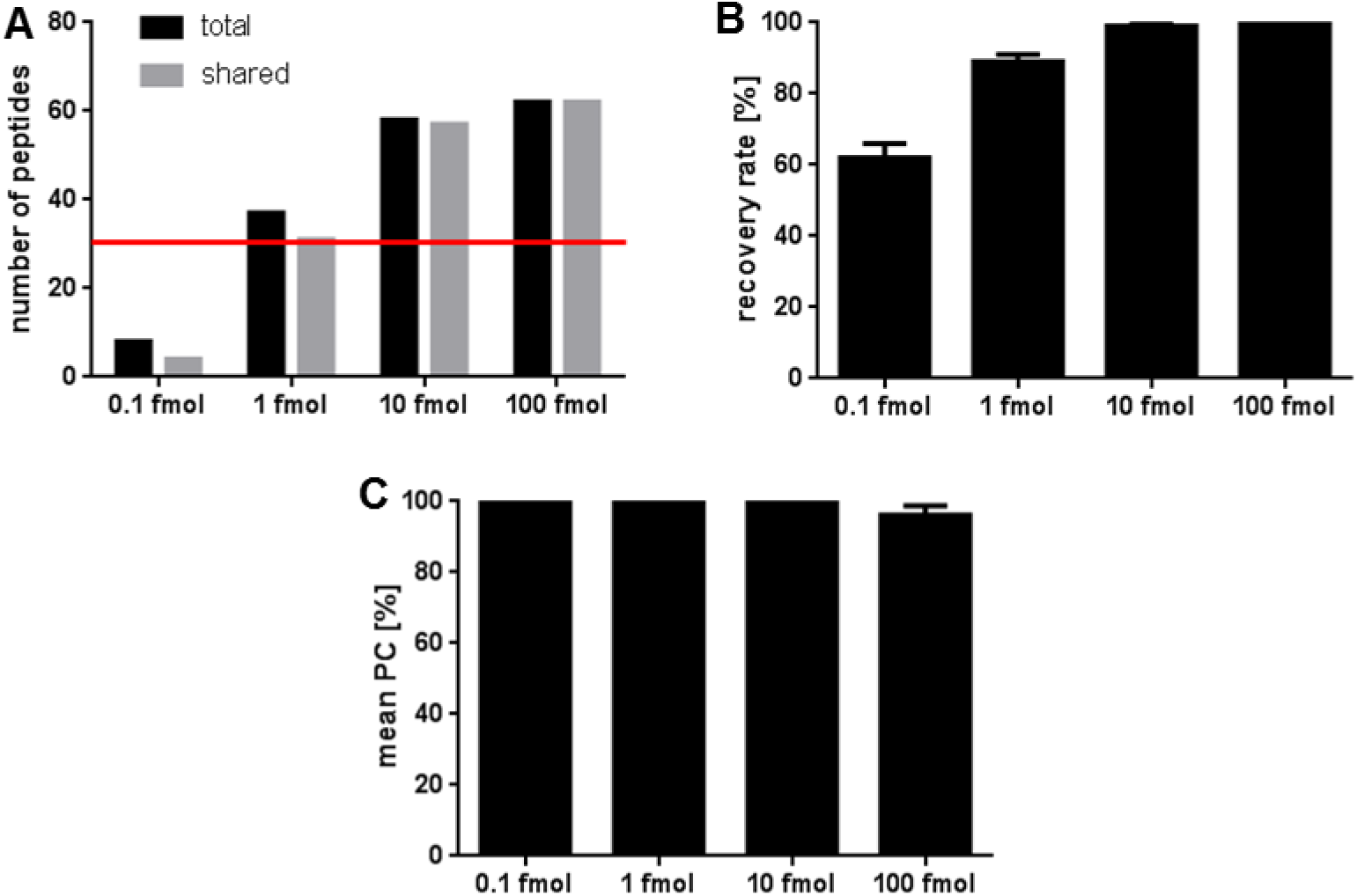
Validation of the limit of detection using spiked isotope labeled synthetic peptides. Three replicates of JY samples spiked with 0.1 fmol, 1 fmol, 10 fmol, and 100 fmol isotope labeled synthetic peptides were analyzed. (A) Number of total identified isotope labeled peptides and shared peptides identified in the three replicates. The LOD of 50% (n = 31) of the spiked synthetic peptides is indicated with a red line. (B) Recovery rate of synthetic peptides recovered between the three replicates of each condition. (C) Mean Pearson correlation of the peptide retention times between the replicates. Abbreviation: PC, Pearson correlation.

### Precision

The precision (also referred to as imprecision) was determined by assaying three aliquots of HLA-eluted peptides from JY samples in three technical replicates leading to nine separate analytical replicates. To prove the intermediate precision, the measurement series was repeated after seven days (for identified peptides, see supplemental Table S5). The number of identified peptides fulfilled the acceptance criteria of ± 10% SD of the repeatability on the initial day and after one week (Figure 5A, B). Furthermore, the acceptance criteria of the recovery rate with a recovery of 80 ± 20% of identified peptides in a repeated replicate were fulfilled on both measuring days (Figure 5C). A closer look at the LC demonstrates that the mean Pearson correlation of the peptide retention times between all nine replicates was above 95% and fulfilled the criteria of the repeatability and intermediate precision (Figure 5D).

**Figure 5:**
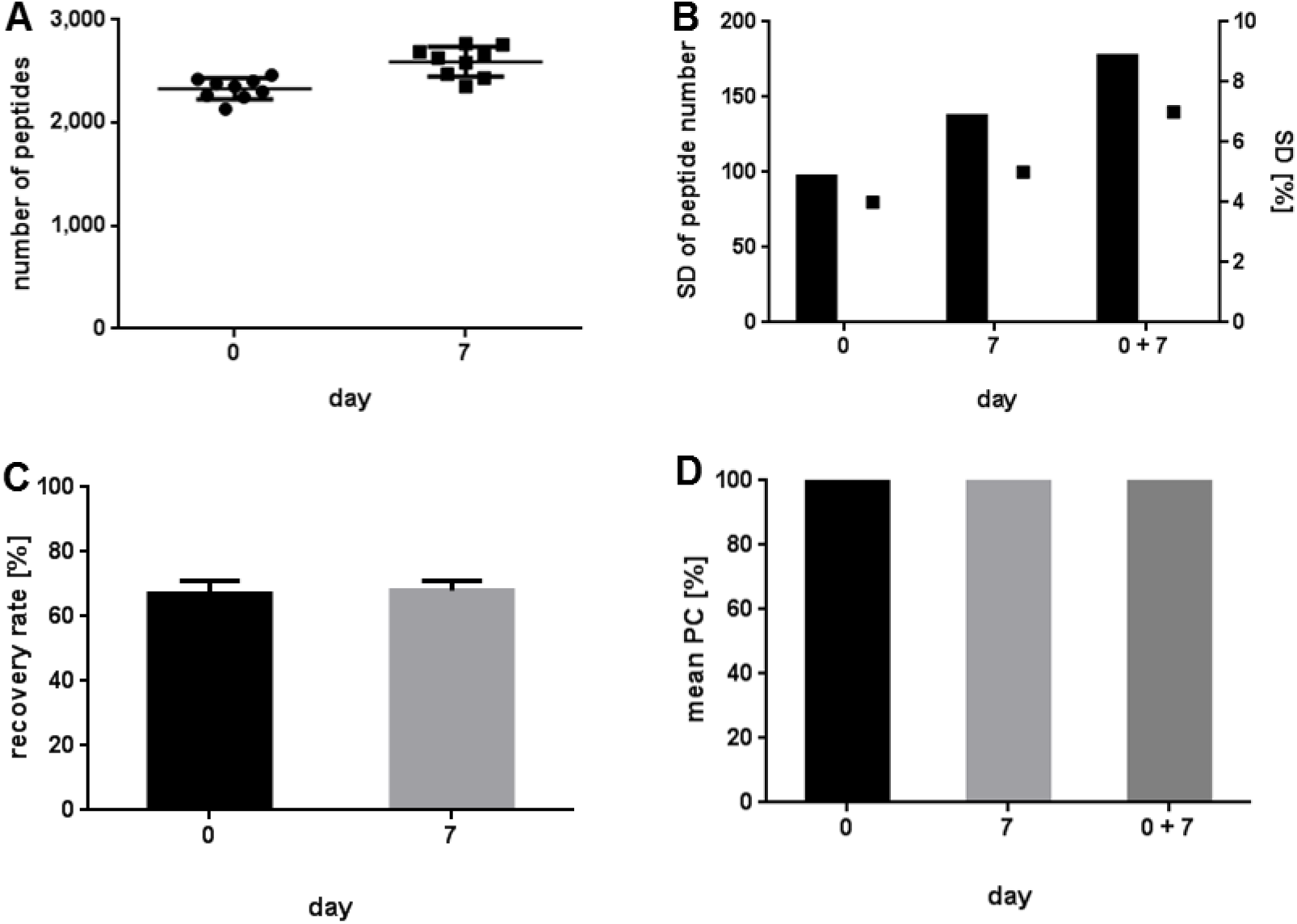
Validation of the precision using immunopeptidomes from JY cells. (A) Number of synthetic peptides identified at day 0 and day 7 in nine replicates, respectively. (B) Standard deviation (SD) given in total peptide numbers and in percent. (C) Recovery rate of peptides between the replicates. (D) Mean Pearson correlation of the peptide retention times between the replicates. Abbreviation: SD, standard deviation; PC, Pearson correlation.

### Robustness of the precision, accuracy and specificity

The robustness was investigated performing a retrospective analysis of the purified HLA-eluted peptides of three primary samples: peripheral blood mononuclear cells from a healthy donor, a chronic lymphocytic leukemia sample, as well as a bladder cancer sample. The immunopeptidomes were isolated and analyzed by three different persons. To validate the robustness the specifications indicated in Table 1 should be fulfilled with regard to accuracy, precision, and specificity.

The identified peptides of the three primary samples fulfill the acceptance criteria of the accuracy for the mass spectrometer and for the LC. The difference of the median of the mass deviation from the theoretical mass of all identified natural peptides (median ΔM: PBMC −0.09 ppm, CLL 0.04 ppm, BC 0.12 ppm) (Figure 6A) and synthetic peptides (median ΔM: 0.32 ppm) is below 1 ppm (Figure 6A). The peptide RTs between the replicates of all natural and all synthetic peptides have a Pearson correlation above 95% (Figure 6B).

**Figure 6:**
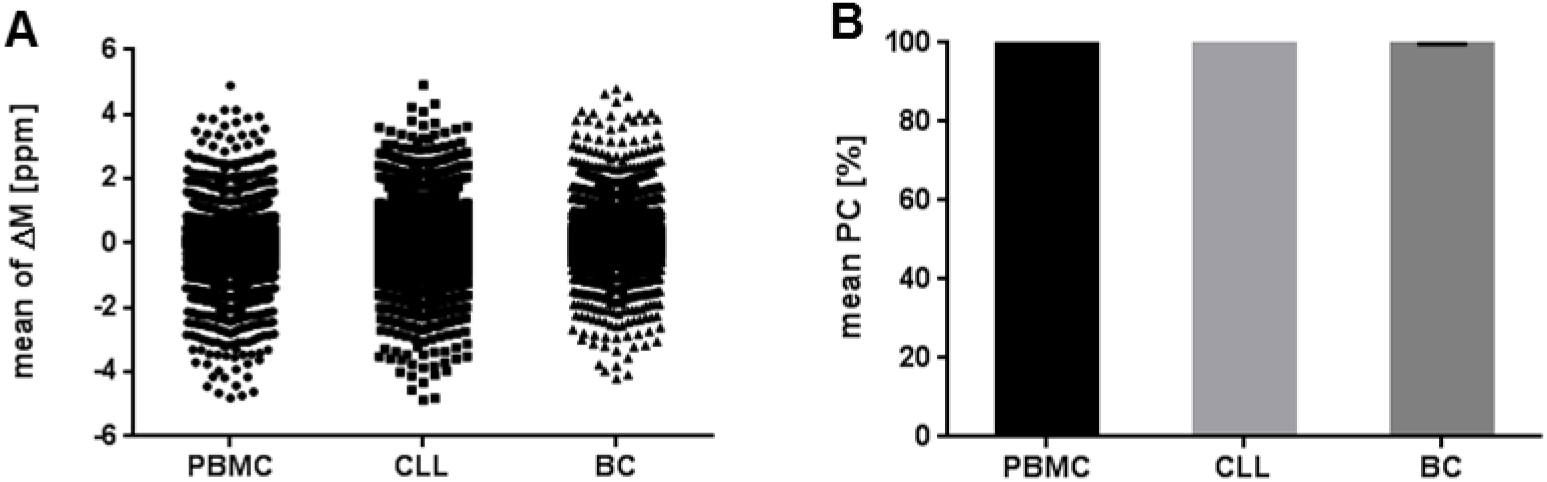
Validation of the accuracy using immunopeptidomes from primary PBMC, CLL, and BC samples. Three replicates were analyzed. (A) Mass deviation from the theoretical precursor mass (ΔM ppm) to the theoretical mass of all identified natural peptides. (B) Mean Pearson correlation of the peptide retention times between the replicates. Abbreviations: PC, Pearson correlation; ppm, parts per million; ΔM, mass deviation.

Regarding the precision, the three technical replicates of the three primary samples fulfill the acceptance criteria of the repeatability for both the mass spectrometer and the LC. The replicates have a percentage SD of identified peptide numbers below 10% (Figure 7A, B) and the recovery rate is in the range of 80% ± 20% (Figure 7C). The peptide RTs between the replicates have a Pearson correlation above 95% (Figure 6B).

**Figure 7:**
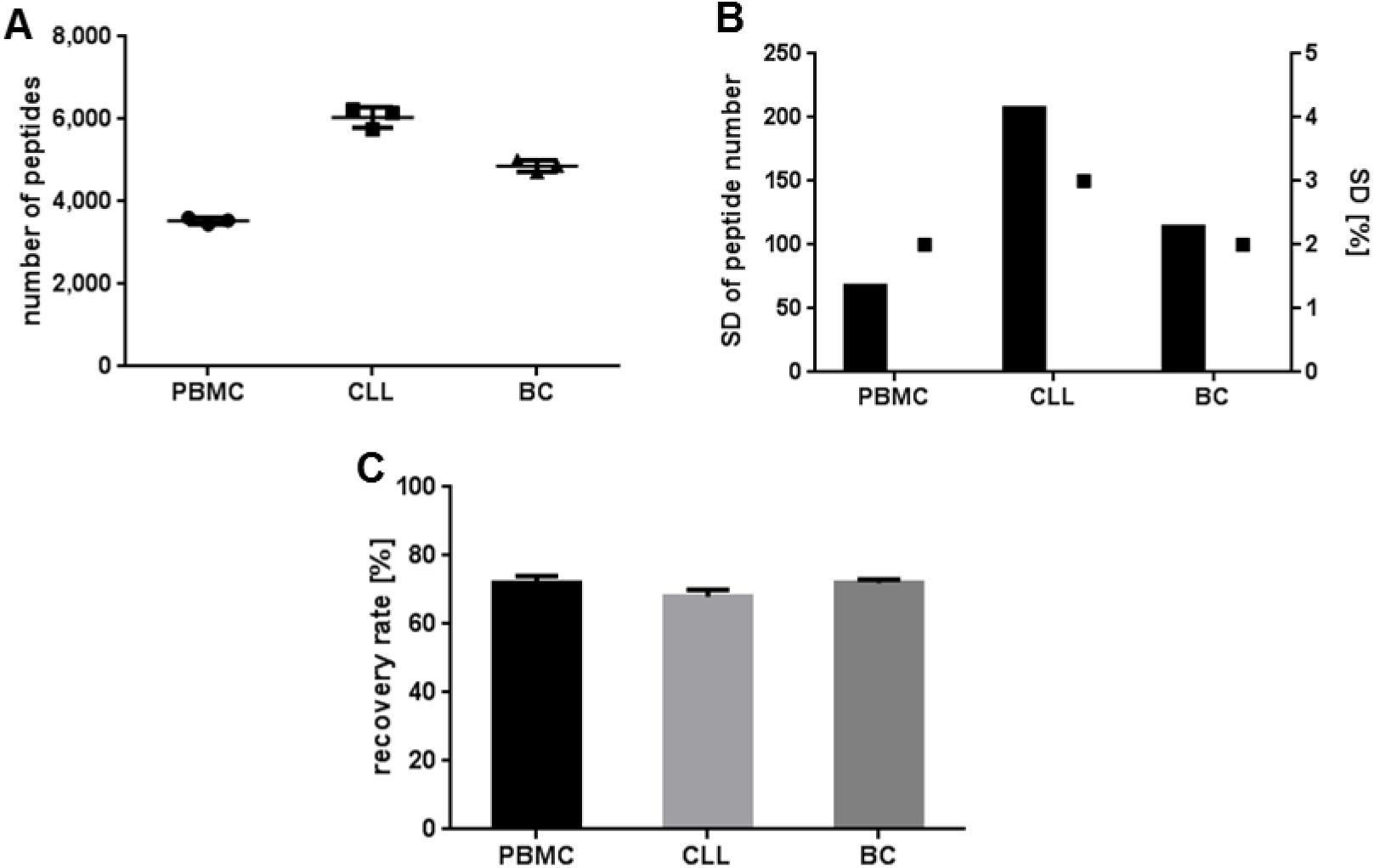
Validation of the repeatability using immunopeptidomes from primary PBMC, CLL, and BC samples. (A) Total number of natural peptides identified in three technical replicates, respectively. (B) Standard deviation (SD) given in total peptide numbers and in percent. (C) Recovery rate of peptide identifications between the replicates. Abbreviation: SD, standard deviation.

The specificity can be investigated, as all analyzed primary samples are expected to contain at least one of the previously used 62 synthetic peptides as natural HLA class I-presented peptide. The natural peptides should fulfill the acceptance criteria of the accuracy and specificity indicated in Table 1 for the mass spectrometer and the LC, respectively (for identified peptides and product ions, see supplemental Table S6).

The three replicates fulfill the acceptance criteria of the specificity for the mass spectrometer and for the LC. The difference of the median of the mass deviation from the theoretical mass (ΔM ppm) of the selected peptides AIVDKVPSV and YLLPAIVHI in PBMC and CLL and SLLQHLIGL in BC, which were identified as natural and 100 fmol spiked synthetic peptides, is below 1 ppm (Figure 8A).

**Figure 8:**
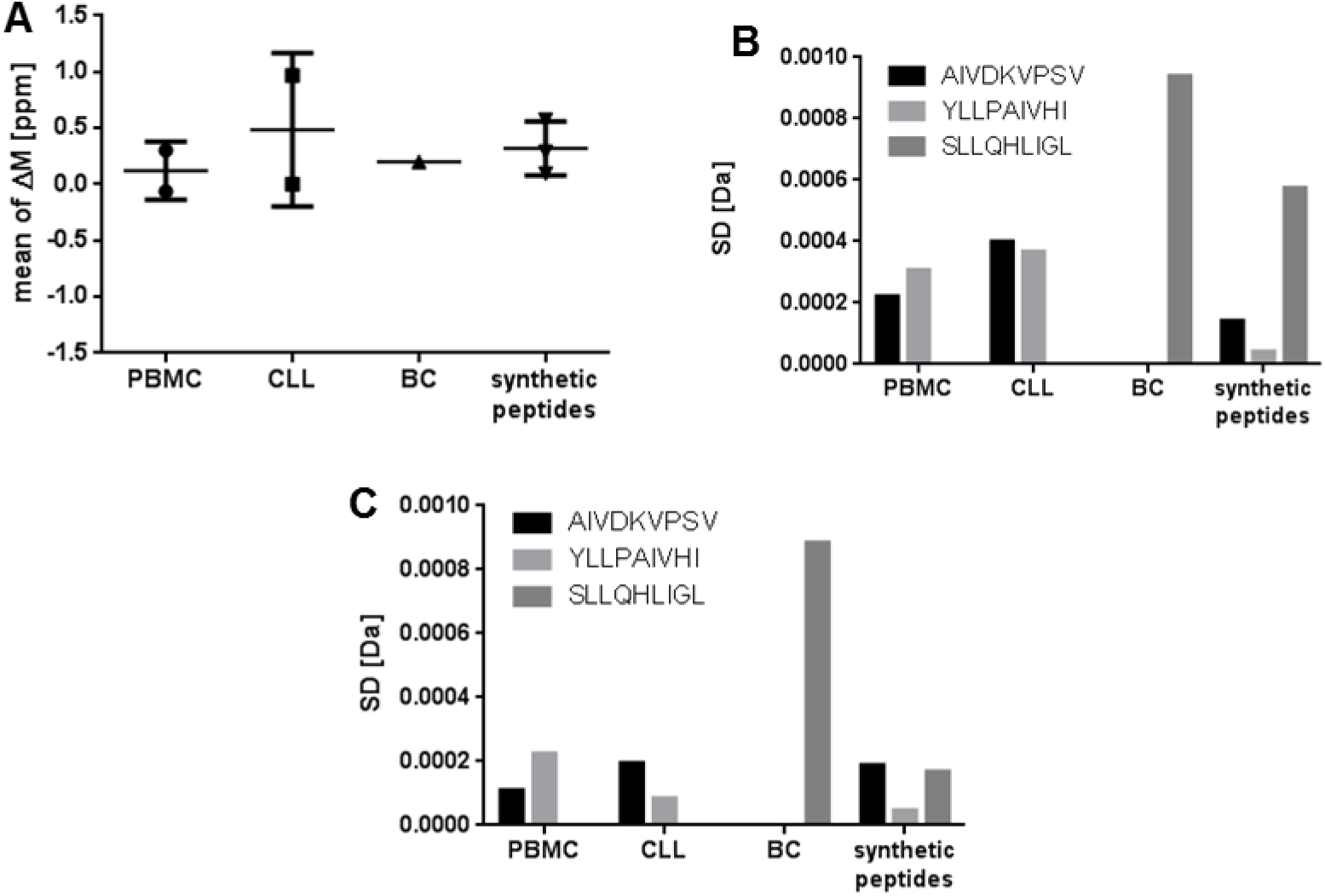
Validation of the specificity using immunopeptidomes from primary PBMC, CLL, and BC samples and isotope labeled synthetic peptides spiked into JY. The samples were analyzed in three replicates. (A) Mean mass deviation of the detected precursor masses from the theoretical masses (ΔM ppm) of the identified natural peptides AIVDKVPSV and YLLPAIVHI (in PBMC and CLL) and SLLQHLIGL (in BC) and 100 fmol synthetic peptides spiked into JY. (B) Mass deviation as SD of the precursor masses detected in MS mode and the resulting five selected top fragments in MS/MS modes of the identified natural and synthetic peptides. Abbreviations: ppm, parts per million; ΔM, mass deviation; SD, standard deviation.

The SD of the mass accuracy of the precursor masses in MS mode and of the selected five top product ions in MS/MS mode of the selected peptides AIVDKVPSV and YLLPAIVHI in PBMC and CLL and SLLQHLIGL in BC is below 0.001 Da for both the natural and synthetic peptides (Figure 8B and C, for MS/MS spectra, see supplemental Figure S1).

### Transfer of the method to other LC-MS/MS systems

In addition to the previous robustness analyses, the method was transferred to an LC-MS/MS system with a less sensitive LTQ Orbitrap XL and the HLA-eluted peptides from JY cells were analyzed. To demonstrate the method transfer to another LC-MS/MS system, the robustness measurements of the method were investigated with regard to accuracy, precision, and specificity on the LTQ Orbitrap XL containing LC-MS/MS system regardless of the specifications indicated in Table 1 set for an Orbitrap Fusion Lumos containing LC-MS/MS system. In order to increase the number of identified peptides, the MS/MS analysis is performed in the ion trap to enable a faster scanning throughput. Consequently, different peptide spectra are expected using adapted settings (described in experimental procedures) and therefore besides the JY samples also 500 fmol synthetic peptides of the six selected sequences AIVDKVPSV, SPQGRVMTI, RPSGPGPEL, YLLPAIVHI, KVLEYVIKV, and SPSSILSTL were spiked in JY matrix and measured on the LTQ Orbitrap XL for a spectral comparison. New five top product ions were selected based on intensity and relevance in MS/MS mode for the LTQ Orbitrap XL system. To investigate the accuracy, the purified HLA-eluted peptides from one JY batch were analyzed in three separate analytical replicates. Regarding the accuracy of the mass spectrometer in MS mode the median mass deviation from the theoretical mass of all identified natural peptides (median ΔM: −0.38 ppm) is below 1 ppm in Figure 9A similar to the Orbitrap Fusion Lumos system. The peptide RTs between the replicates of all natural peptides do have a mean Pearson correlation above 95%, demonstrating the accuracy of the LC (Figure 9B).

**Figure 9:**
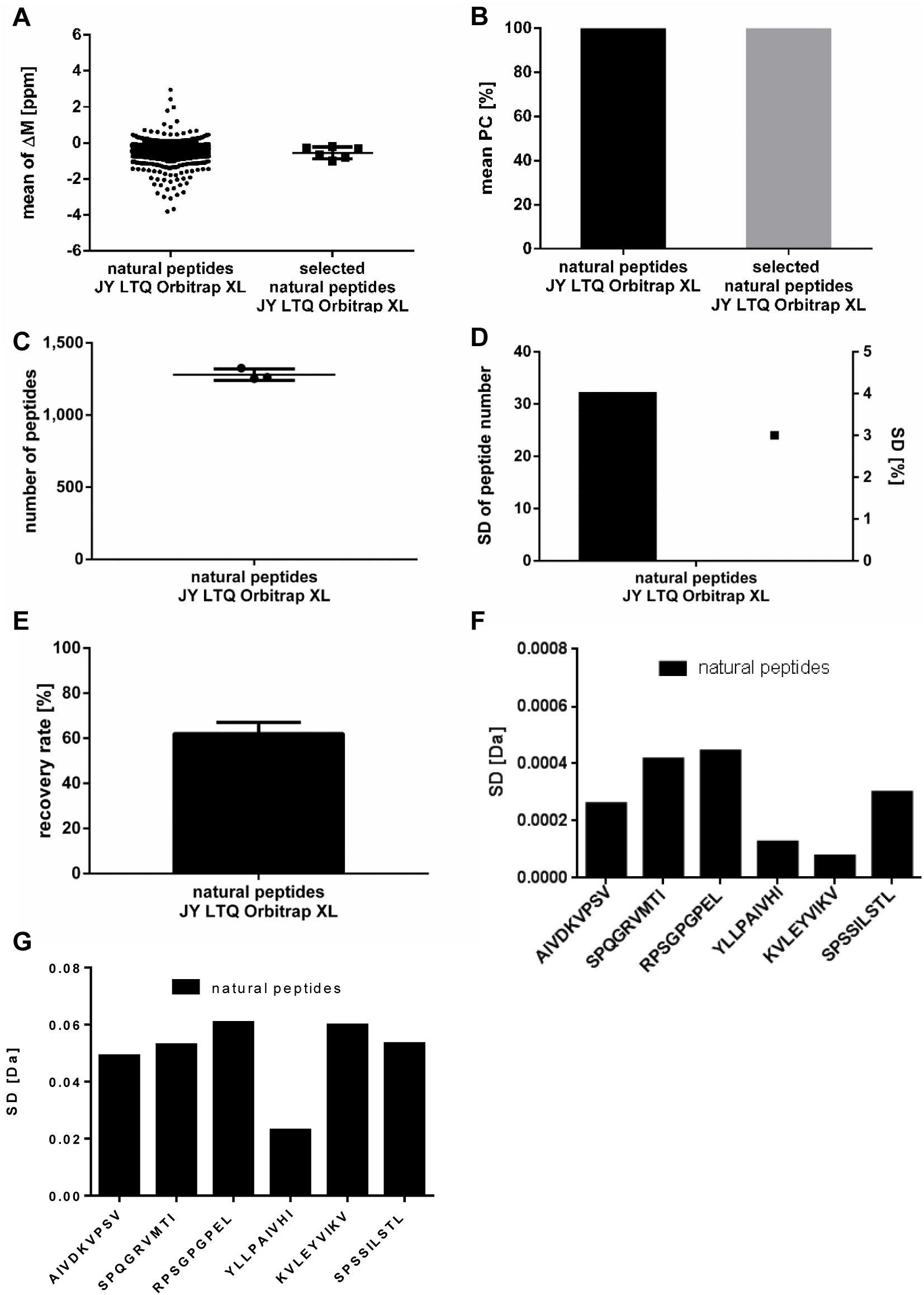
Investigation of the accuracy, repeatability and specificity using immunopeptidomes from JY cells analyzed after method transfer to a less sensitive LC-MS/MS system. Three replicates were analyzed. (A) Mass deviation from the theoretical precursor mass (ΔM ppm) to the theoretical mass of all identified natural peptides and of the six identified natural peptides AIVDKVPSV, SPQGRVMTI, RPSGPGPEL, YLLPAIVHI, KVLEYVIKV, and SPSSILSTL. (B) Mean Pearson correlation of the peptide retention times between the replicates for all identified natural peptides and the six selected peptides. (C) Total number of natural peptides identified in three technical replicates, respectively. (D) Standard deviation (SD) given in total peptide numbers and in percent. (E) Recovery rate of peptide identifications between the replicates. Abbreviation: SD, standard deviation. Mass deviation as SD of the (F) precursor ion masses in MS mode and the (G) resulting five selected top fragments in MS/MS modes of the six selected natural and synthetic peptides. Abbreviations: PC, Pearson correlation; ppm, parts per million; ΔM, mass deviation; SD, standard deviation.

The precision was determined by assaying the previously mentioned three technical replicates. The number of identified peptides was similar to the validated LC-MS/MS system with 3% SD of the repeatability on the initial day (Figure 9C, D). However, the recovery rate with a recovery of 62 ± 5% of identified peptides in a repeated replicate is different (Figure 9E).

The specificity was again investigated using the mass deviation from the theoretical mass of the six selected peptides AIVDKVPSV, SPQGRVMTI, RPSGPGPEL, YLLPAIVHI, KVLEYVIKV, and SPSSILSTL. The median of the identified natural peptides is ΔM: −0.50 ppm (Figure 9A) (for identified peptides and product ions, see supplemental Table S7).

The peptide RTs between the replicates of the six selected peptides identified as natural and synthetic peptides do have a Pearson correlation above 95% (Figure 9B).

The standard deviation (SD) of the mass accuracy of the precursor masses in MS mode (Figure 9F) and of the five selected top product ions in MS/MS mode (Figure 9G, for MS/MS spectra, see supplemental Figure S3), selected based on intensity and relevance, of the six selected peptides AIVDKVPSV, SPQGRVMTI, RPSGPGPEL, YLLPAIVHI, KVLEYVIKV and SPSSILSTL is below 0.001 Da in MS Mode, analyzed in the Orbitrap, and below 0.1 in MS/MS mode, analyzed in the ion trap, for the natural peptides.

## Conclusion/Discussion

To provide reliable biomarker and patient-individual tumor-associated target antigen identification for clinical studies, the fast and sensitive LC-MS/MS assay for the identification of natural and synthetic HLA-restricted peptides was validated for the technical equipment of our laboratory, consisting of a nanoUHPLC, UltiMate 3000 RSLCnano on-line coupled to an Orbitrap Fusion Lumos mass spectrometer. The immunopeptidomics pipeline is used for identification and impurity detection, thus according to FDA and EMA guidelines a validation of the specificity and LOD is required. Additionally, we validated the accuracy, precision, and robustness to demonstrate the reliability of the pipeline.

The results of the JY samples spiked with isotope labeled synthetic peptides enabled verification of the accuracy of the LC-MS/MS system in terms of similarity of the analyzed natural peptides to the theoretical and synthetic peptide masses. With the same dataset we were able to identify six specific peptides, expected as natural and synthetic peptides, which fulfil the acceptance criteria of the accuracy and could prove the specificity. Furthermore, we could show that with five selected MS/MS product ions all identified peptides within one replicate can be distinguished. Instead of picking simply the five most intensive product ions for therapeutic peptide candidates, our quality control selects the top five product ions also according to meaningfulness (expert review: b- or y-ions are preferred), thus further increasing specificity. The validation of the precision showed a reliable identification of peptides with a uniform recovery rate proving the repeatability and intermediate precision after one week.

A major limitation of MS-based data-dependent acquisition (DDA) discovery approaches is the low recovery rate. In our immunopeptidomics experiments a recovery rate of 80% ± 20% was achieved for cell lines and tissue samples, owing to the tissue heterogeneity and high dynamic range. Due to the recovery rate, in our lab routinely triplicate measurements are performed. At the LOD a reasonable reliability should be provided with a recovery rate of 50%, when triplicate measurements are performed. A peptide content of 1 fmol synthetic peptides in JY matrix enabled a reliable identification of 50% of the peptides. An improvement of the recovery rate might be obtained with a replacement of the DDA analysis with data-independent acquisition, which has demonstrated a superior reproducibility^34–38^.

In order to prove the robustness of the immunopeptidomics assay, we synthesized a large variety of known HLA ligands, with different length, mass, grand average of hydropathicity (GRAVY), theoretical isoelectric point (pI), and HLA allotype restriction. In addition, we employed several primary, clinically relevant samples in addition to the JY cell line and further analyzed soluble peripheral blood mononuclear cells from a healthy donor, a soluble chronic lymphocytic leukemia and a solid bladder cancer sample. Lastly, for the three primary samples the HLA immunoaffinity chromatography and immunopeptidomics analysis was performed from three different persons. We could successfully verify the specifications of the accuracy, precision, and specificity for both the mass spectrometer and LC, respectively.

Besides the robustness of the method, we exemplarily further investigated for precision, accuracy and specificity after the method transfer to a LC-MS/MS system utilizing an LTQ Orbitrap XL. However, in order to obtain a high number of identified peptides, the MS/MS analysis is performed in the ion trap, instead of the Orbitrap, to enable a faster scanning throughput. Consequently, the mass accuracy of the product ions in the MS/MS analysis varies already at the second decimal and for the previously selected top five ions, two new ions had to be defined for SPSSILSTL and one new ion for the other peptides, except of AIVDKVPSV. Furthermore, the recovery-rate is much lower using the LTQ Orbitrap XL.

The specifications of our validated Orbitrap Fusion Lumos based LC-MS/MS system are not met by the LTQ Orbitrap XL. In order to fully validate the method on the LTQ Orbitrap XL for peptide identification according to the current FDA and EMA guidelines, at least the analyses regarding specificity and LOD have to be performed or, ideally, all mentioned parameters should be investigated. The specifications of the precision, specificity and LOD have to be adapted to the capabilities of the less sensitive mass spectrometer. However, the specifications should be tight enough to recognize immediately any functional problems with the LC-MS/MS system.

To enable standardization between different MS platforms and laboratories, the same immunopeptidome batch from one cell line should be used for immunopeptidomics. Here we present how the parameters accuracy, specificity, LOD, precision and robustness can be investigated for the validation of LC-MS/MS systems. However, every LC-MS/MS system in every laboratory needs to be validated independently with its own specifications^27^. For the validation the specifications should be adapted as closely as possible to the optimal performance of the respective LC-MS/MS system, but should also consider the harmless device-related performance variations. Furthermore, it must be considered that more sensitive devices can detect peptides in lower quantities that less sensitive devices cannot identify. Here, the Orbitrap Fusion Lumos discovers twice the number of peptides. The validation is performed in a new process to show that the previously specified requirements (acceptance criteria) are reproducibly met in practical use and that the analytical method is appropriate for its intended use. After the validation a system suitability test (SST) is used to continuously monitor the performance of the instrument in different fixed intervals, to verify that an analytical method is suitable for the intended purpose on the day of analysis. Control peptides known from experience to be reliably identified in the respective immunopeptidome of the cell line in higher quantity should be selected for this purpose. The identification and the retention time of these peptides in the immunopeptidomes could be routinely checked in the LC-MS/MS analysis. The selected peptides can be standardized between different suitable LC-MS/MS systems and laboratories. The scope of each LC-MS/MS validation and the SSTs can prove the suitability and comparability of different laboratories for a particular analysis. The immunopeptidomic pipeline is currently in use for the identification of tumor-associated target antigens of multiple patients in the peptide vaccination study iVAC-CLL01(NCT02802943) and for the preparation of further studies (*e.g.*, PepIVAC01). Due to the peptide recovery rate of 80±20% per replicate, patient samples are routinely analyzed in triplicates to ensure a high recovery. Finally, candidate peptide antigens are always verified using synthetic peptides to exclude false positives and artefacts. The identification procedure of tumor-specific HLA ligands using a comparison of ligand source proteins to established tumor antigens or ligand mapping on different malignant and benign tissues was carried out for cell lines and various tumor entities^8,28,39–43^. During the last decades and in ongoing studies the immunopeptidomics pipeline has demonstrated its reliability and applicability.

In addition, we have now validated the accuracy, precision, specificity, LOD, and robustness in line with the current FDA and EMA guidelines. This validated pipeline enables the reliable identification of tumor-associated HLA-presented target antigens to support current and future clinical studies. Furthermore, different validation approaches are presented, that can be translated to other laboratories with similar equipment, or to other MS-based discovery approaches, such as proteomics, metabolomics, and lipidomics.

## Abbreviations

AcN: Acetonitrile
BC: Bladder cancer
CLL: Chronic lymphocytic leukemia
EMA: European Medicines Agency
FDA: Food and Drug Administration
FDR: False discovery rate
GLP: Good laboratory practice
GMP: Good manufacturing practice
HLA: Human leukocyte antigen
LOD: Limit of detection
OECD: Organisation for Economic Co-operation and Development
PBMC: Peripheral blood mononuclear cells
PPM: Parts per million
SD: Standard deviation

## Acknowledgements

This work was supported by the German Cancer Consortium (DKTK) and the Natural and Medical Sciences Institute at the University of Tübingen NMI. We thank the Wirkstoffpeptidlabor, especially Patricia Hrstić, Ulrich Wulle, Nicole Bauer, Camille Supper, and Mirijam Bohn for expert peptide synthesis and quality control.

