## Supplemental Figure S1 for "Validation of a high-performance liquid chromatography-tandem mass spectrometry immunopeptidomics assay for the identification of HLA class I ligands suitable for pharmaceutical therapies"

### Slide 1
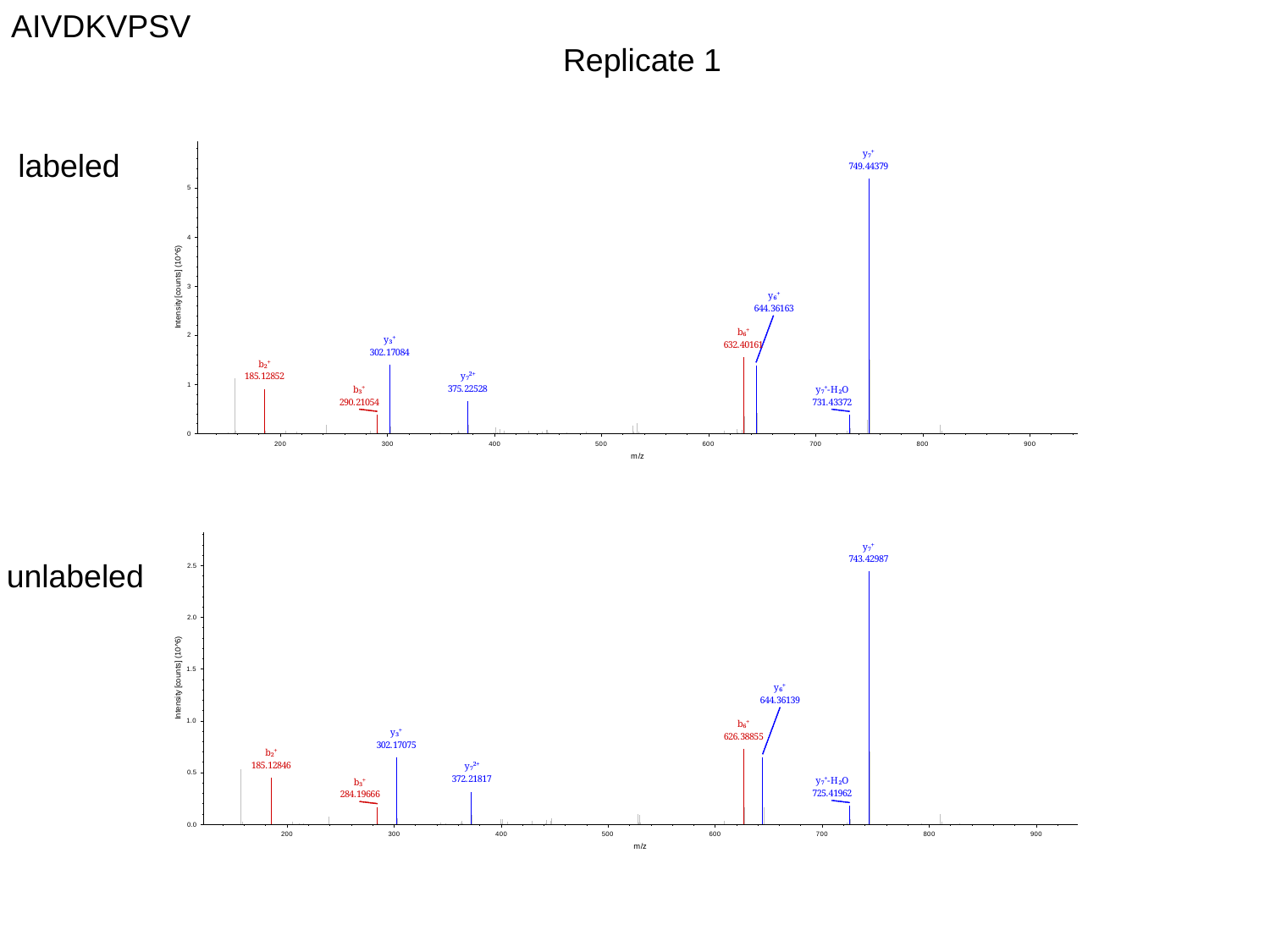

AIVDKVPSV
Replicate 1
labeled
unlabeled

### Slide 2
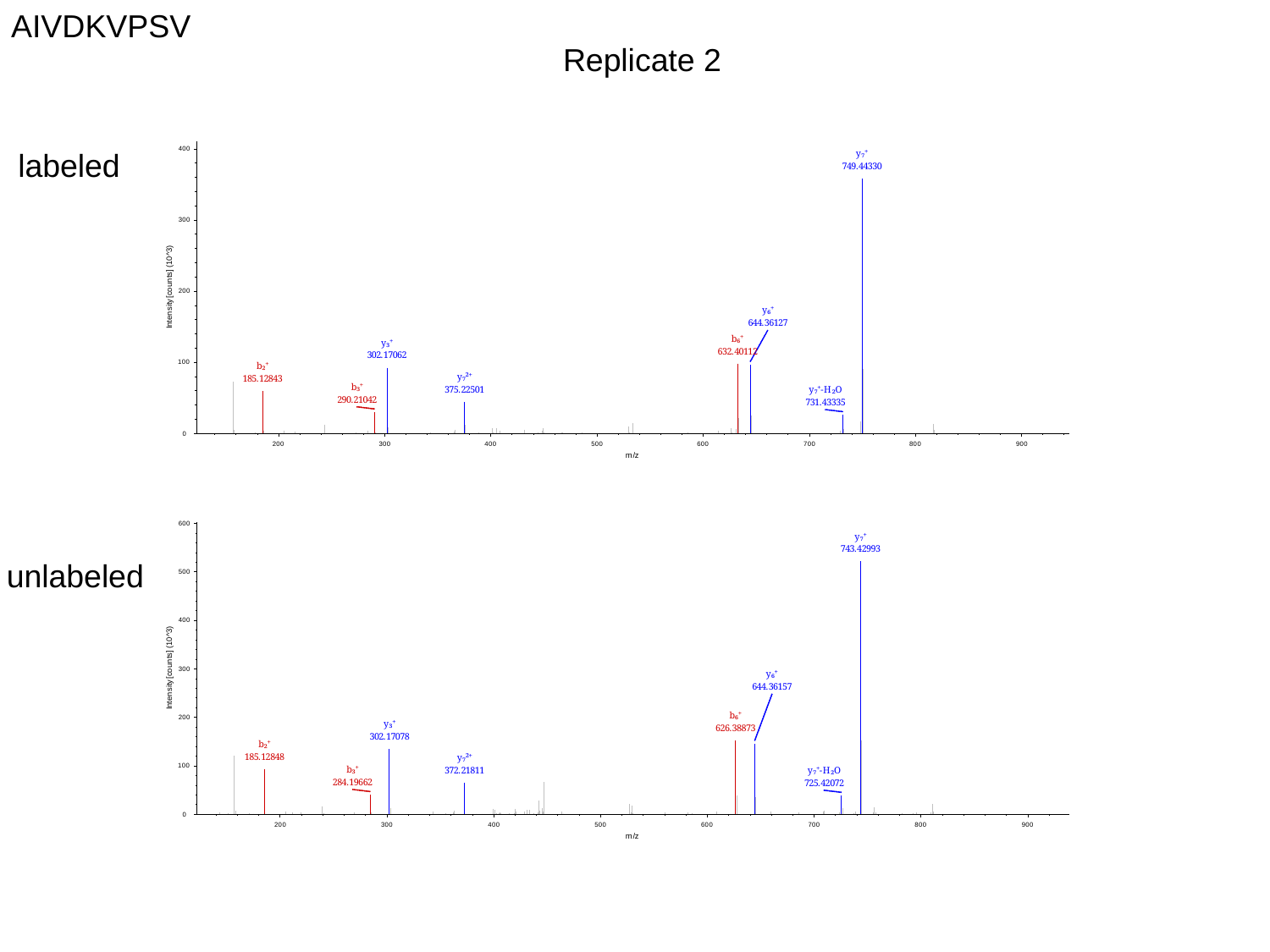

AIVDKVPSV
Replicate 2
labeled
unlabeled

### Slide 3
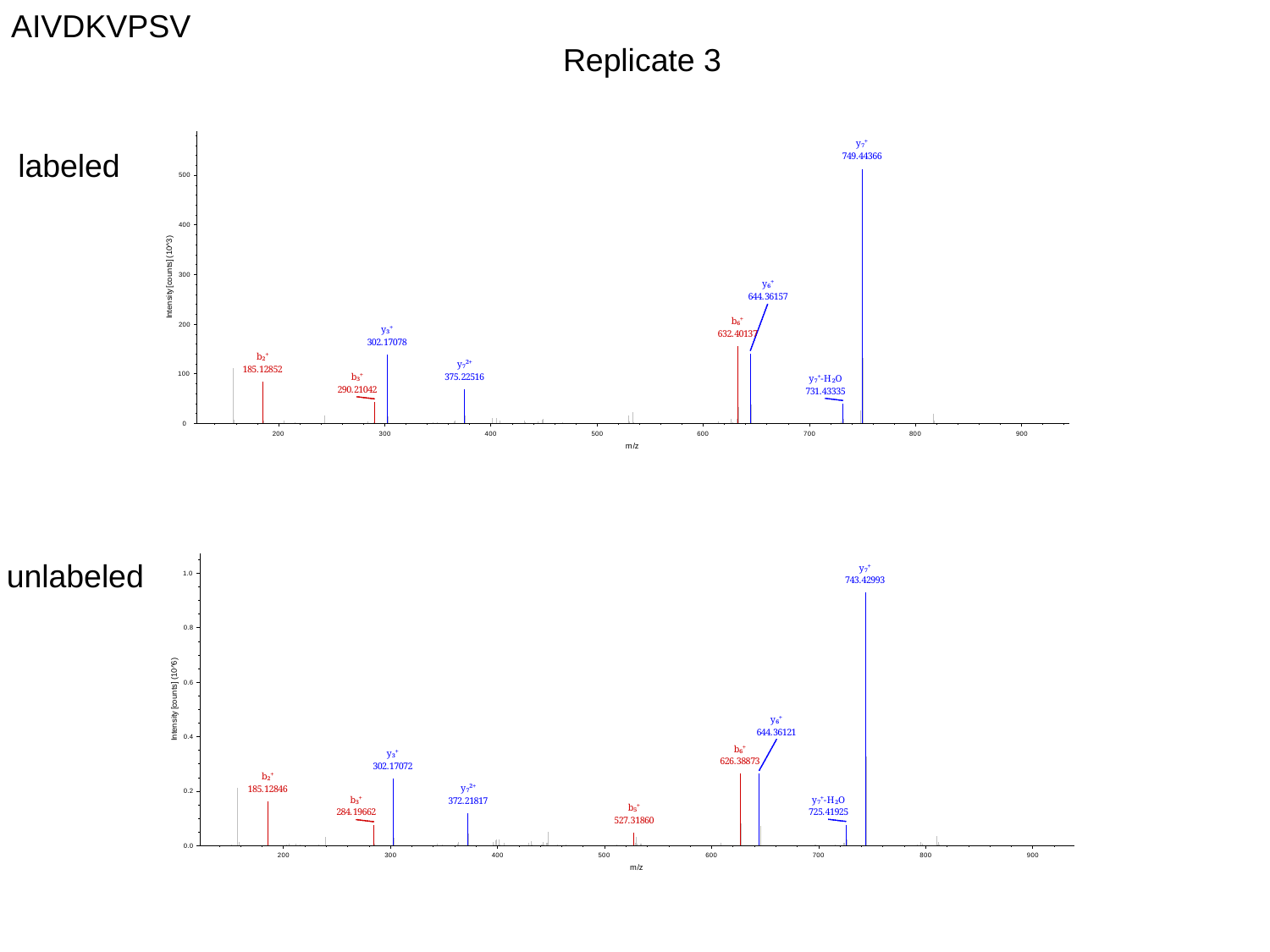

AIVDKVPSV
Replicate 3
labeled
unlabeled

### Slide 4
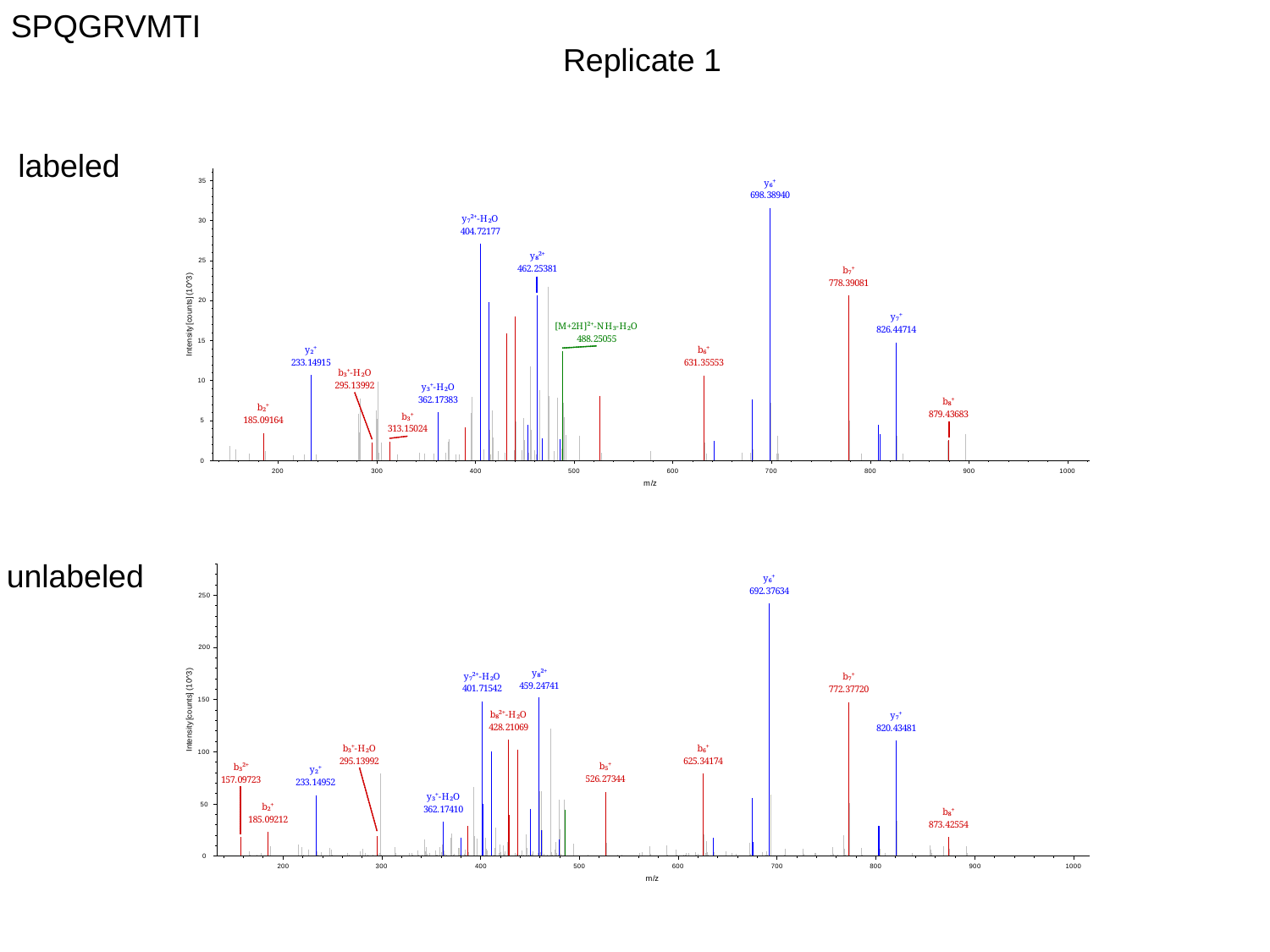

SPQGRVMTI
Replicate 1
labeled
unlabeled

### Slide 5
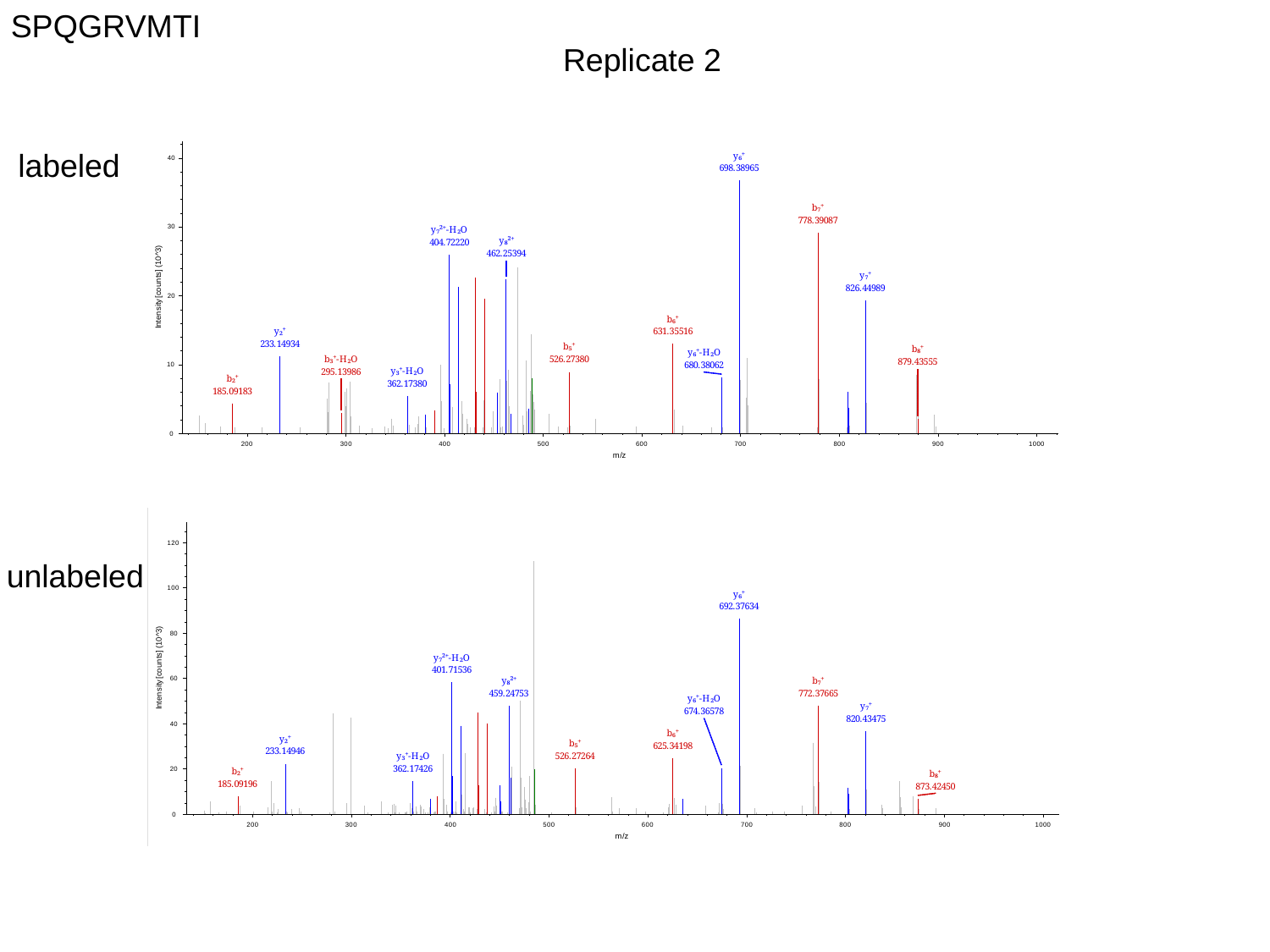

SPQGRVMTI
Replicate 2
labeled
unlabeled

### Slide 6
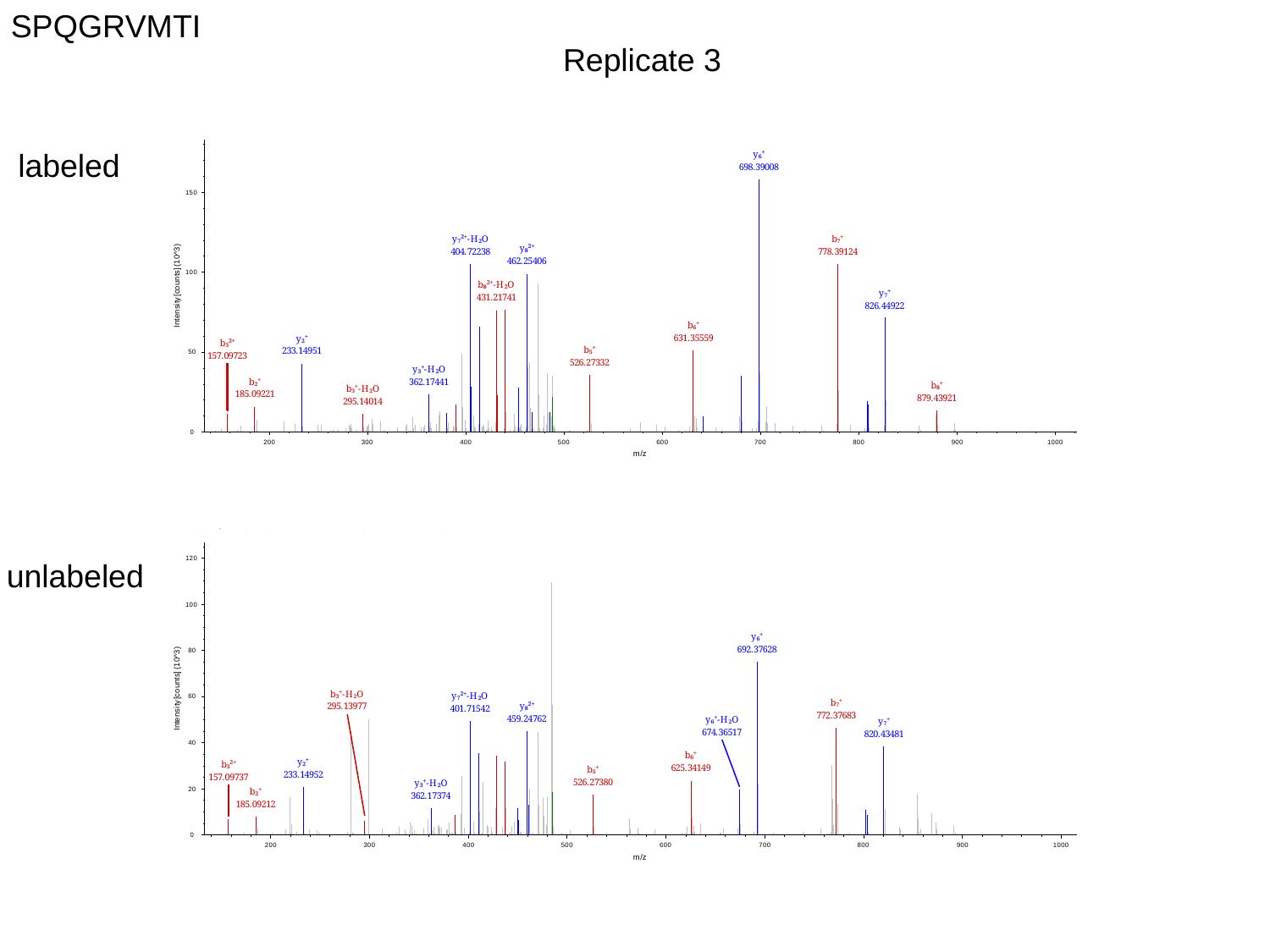

SPQGRVMTI
Replicate 3
labeled
unlabeled

### Slide 7
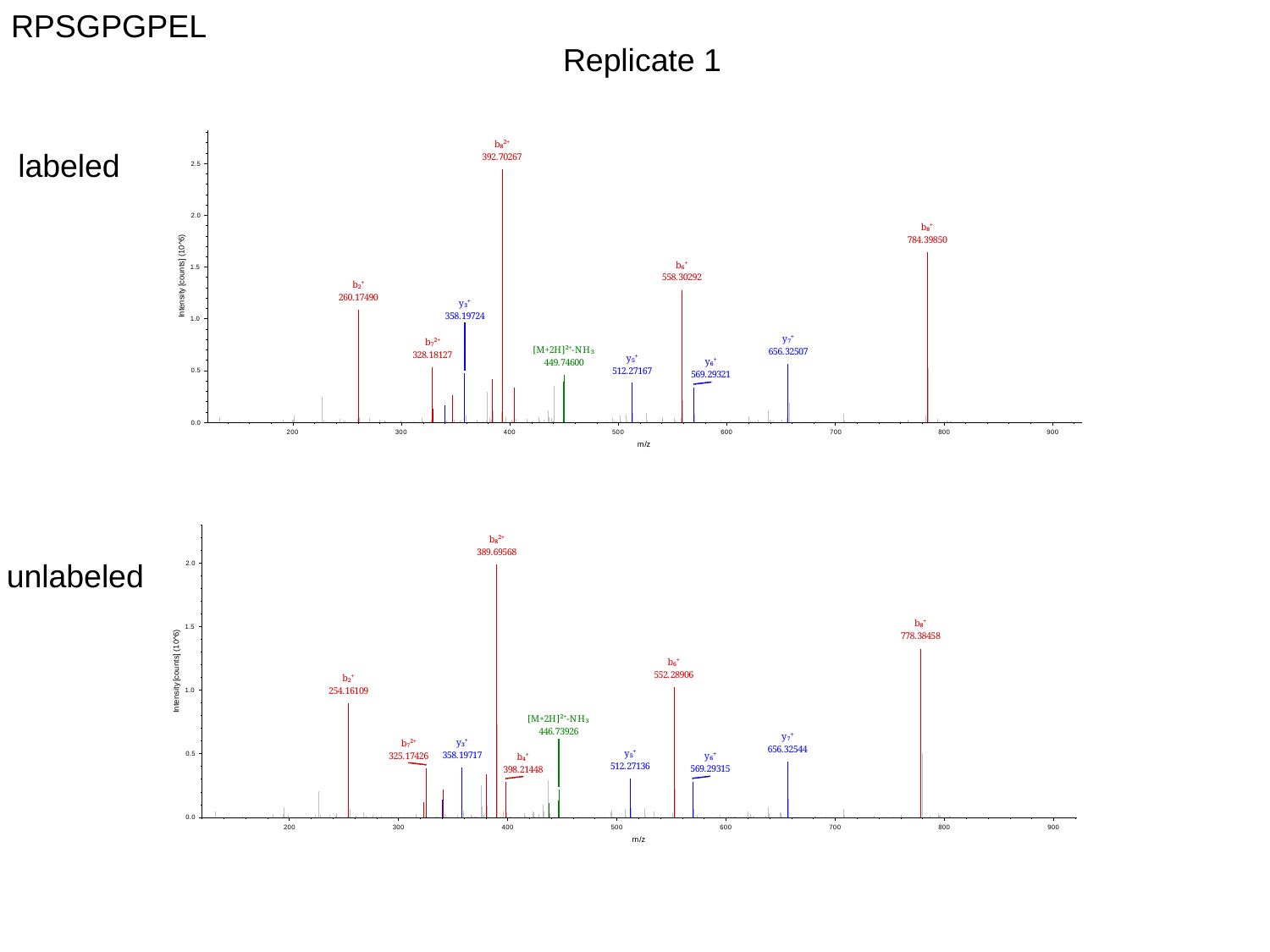

RPSGPGPEL
Replicate 1
labeled
unlabeled

### Slide 8
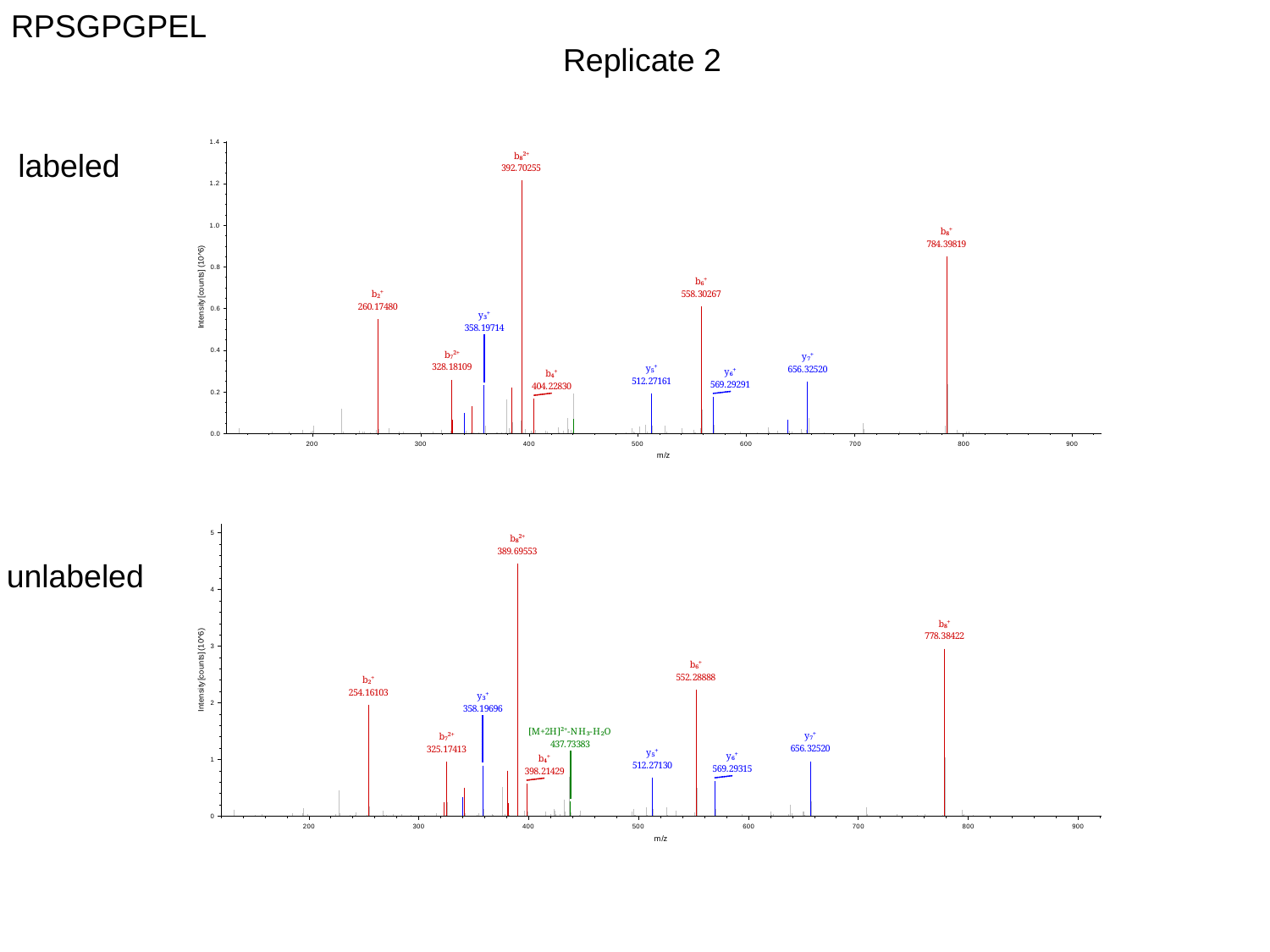

RPSGPGPEL
Replicate 2
labeled
unlabeled

### Slide 9
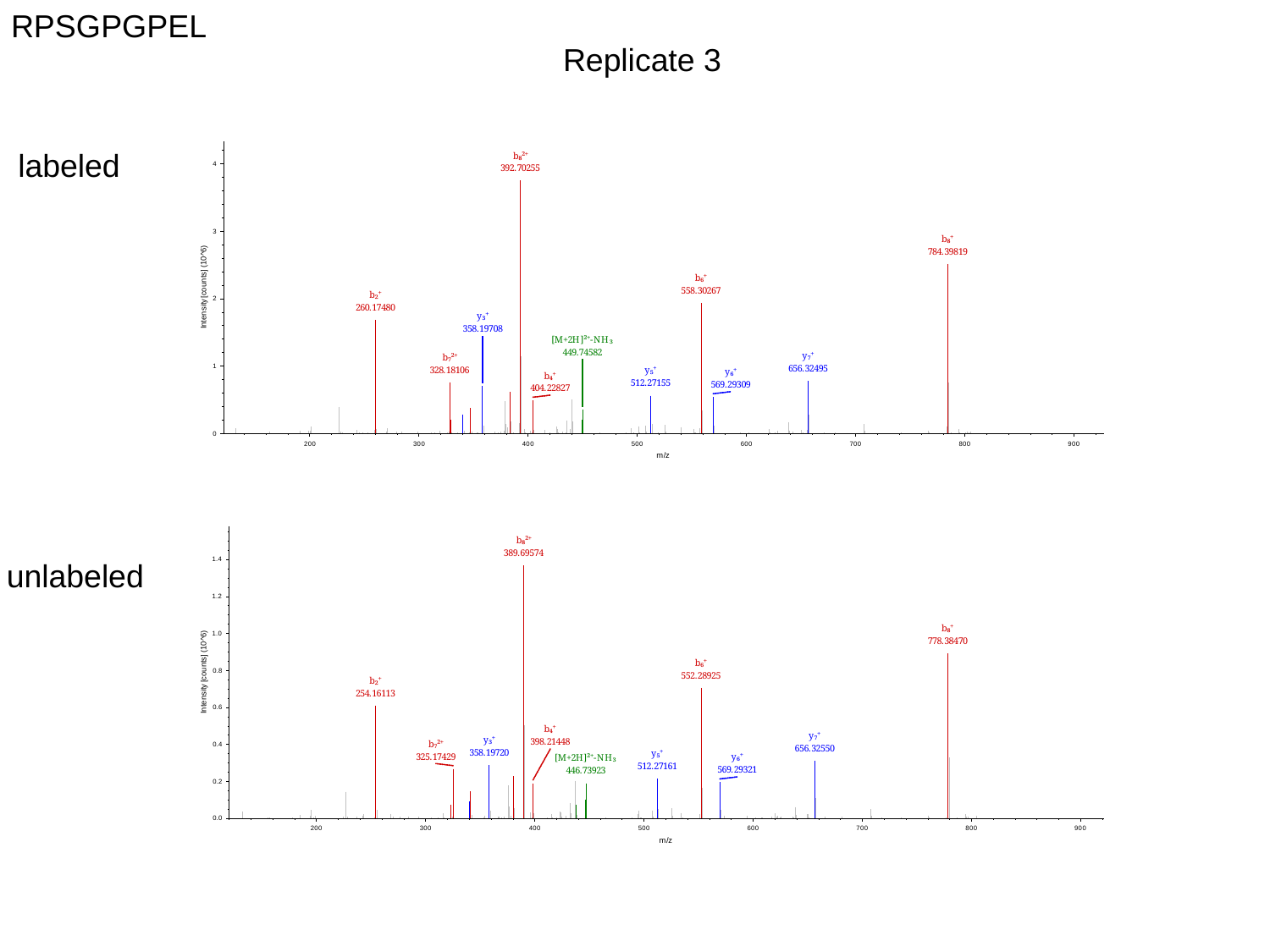

RPSGPGPEL
Replicate 3
labeled
unlabeled

### Slide 10
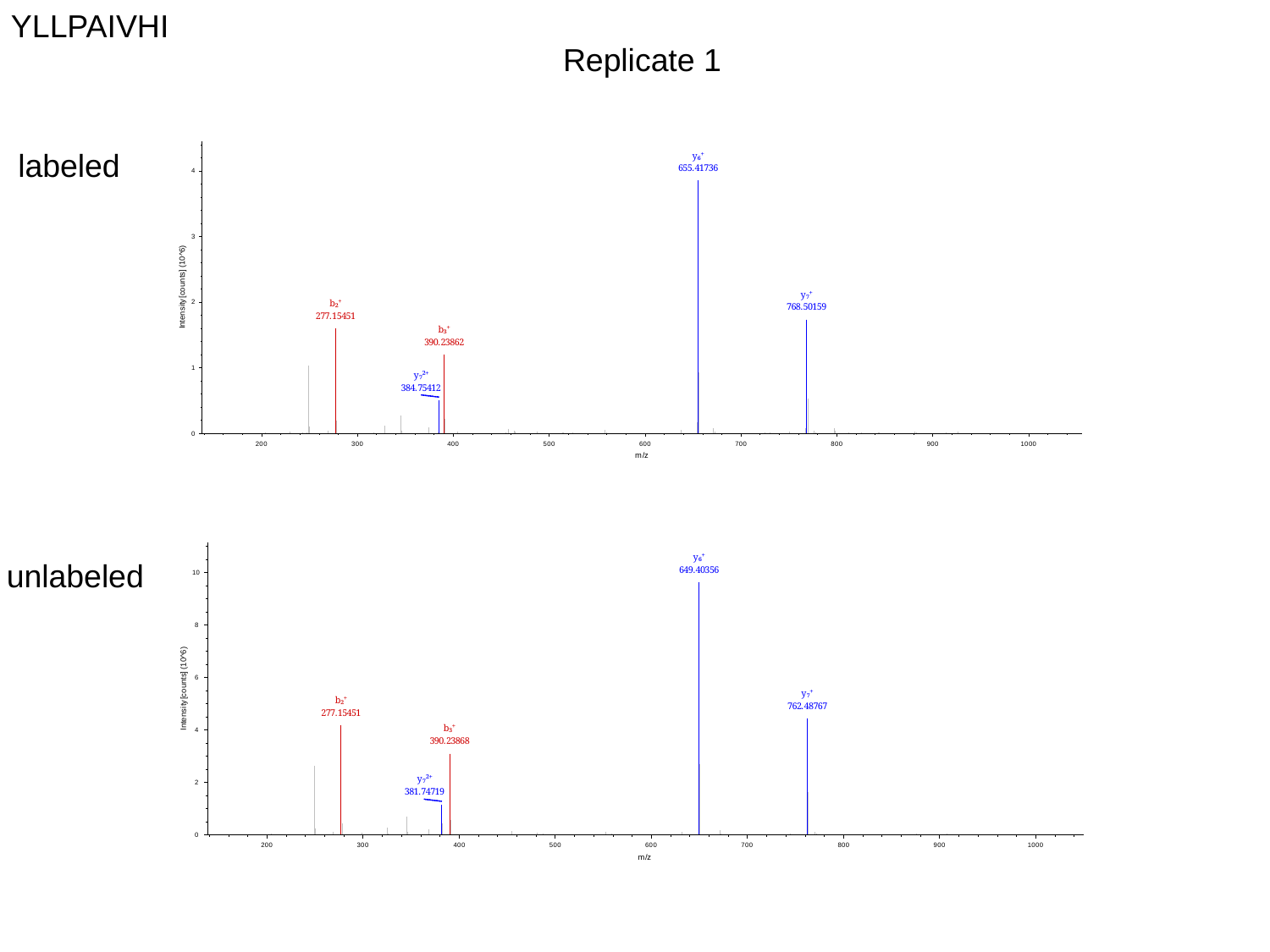

YLLPAIVHI
Replicate 1
labeled
unlabeled

### Slide 11
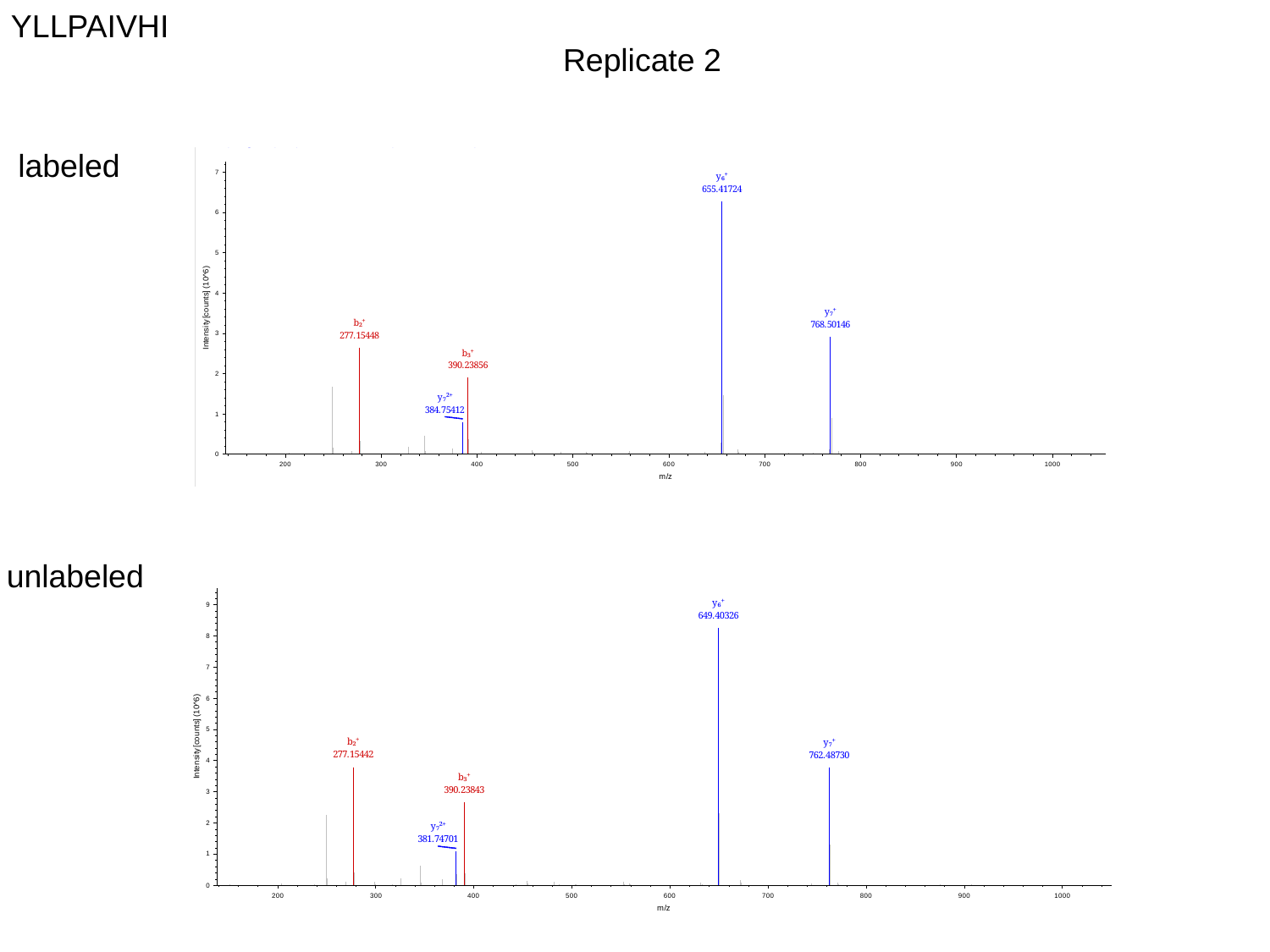

YLLPAIVHI
Replicate 2
labeled
unlabeled

### Slide 12
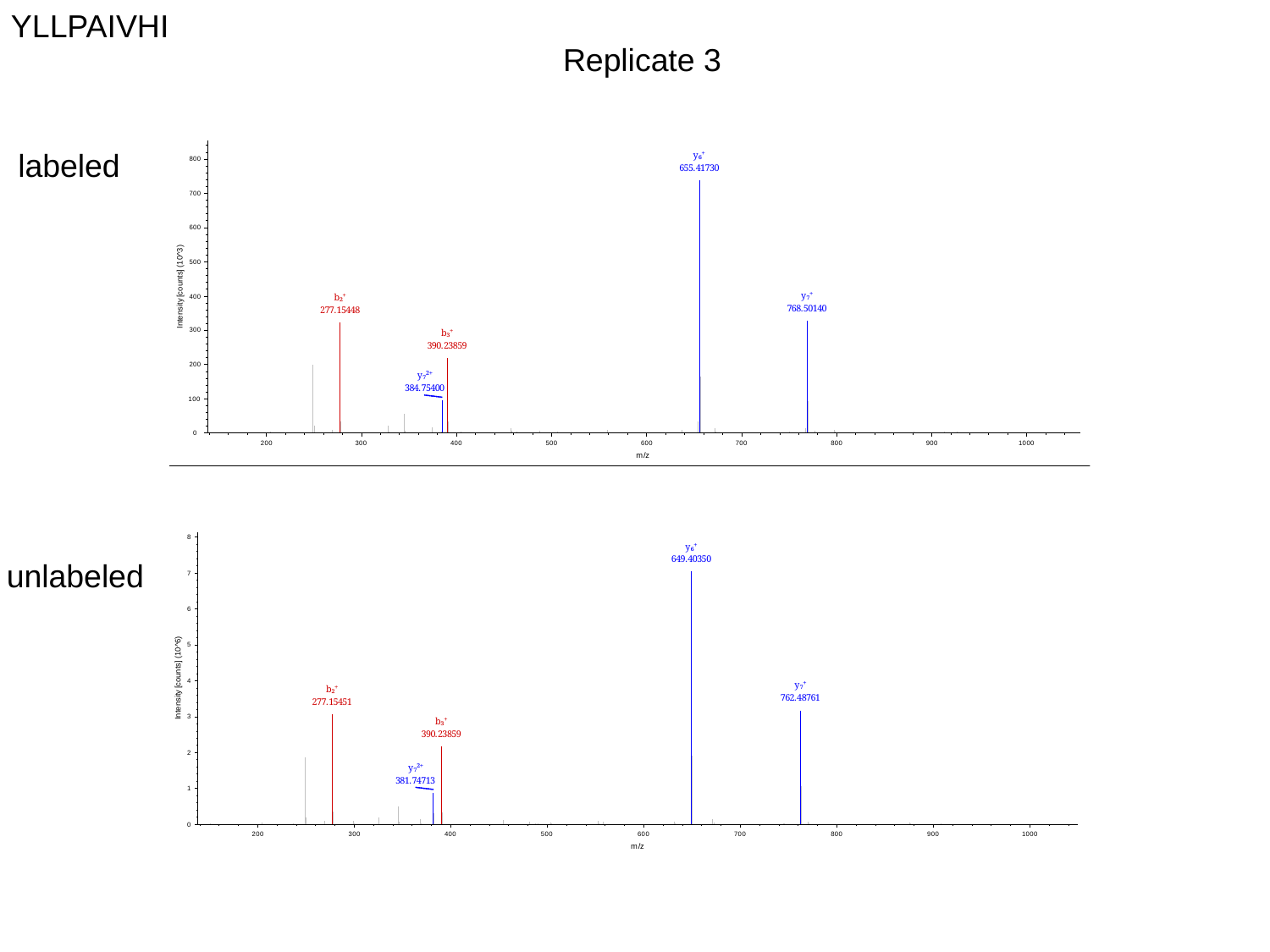

YLLPAIVHI
Replicate 3
labeled
unlabeled

### Slide 13
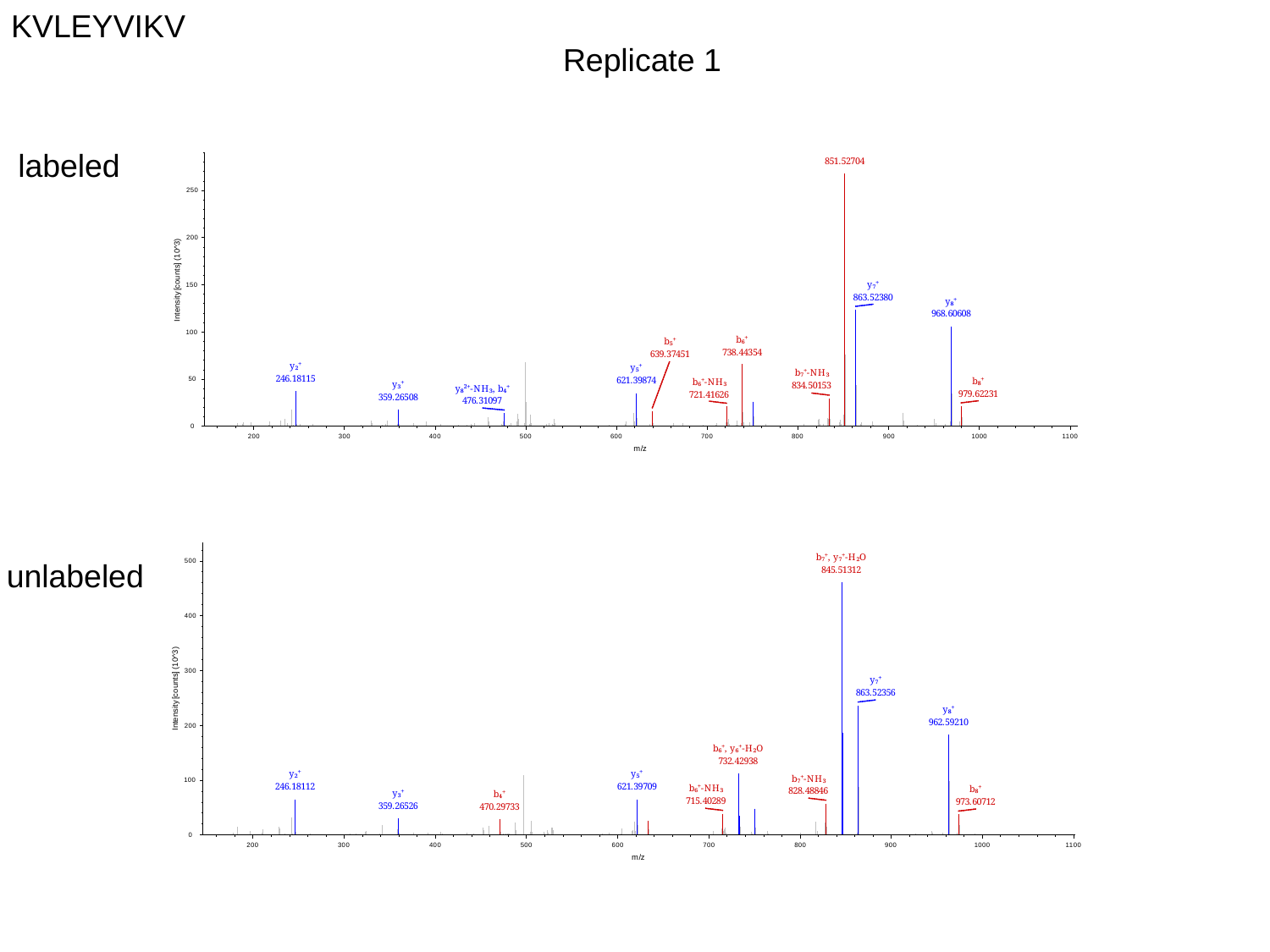

KVLEYVIKV
Replicate 1
labeled
unlabeled

### Slide 14
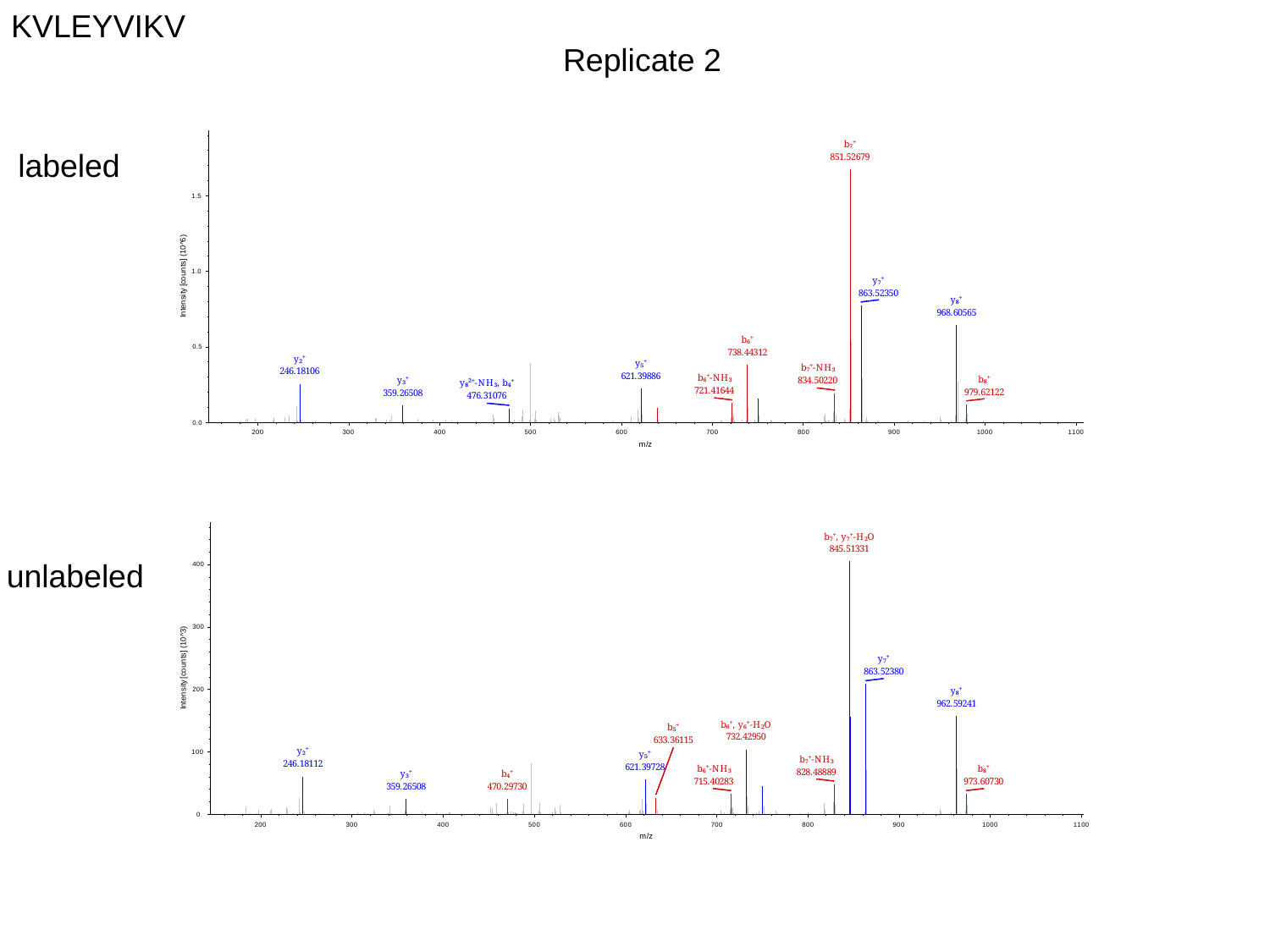

KVLEYVIKV
Replicate 2
labeled
unlabeled

### Slide 15
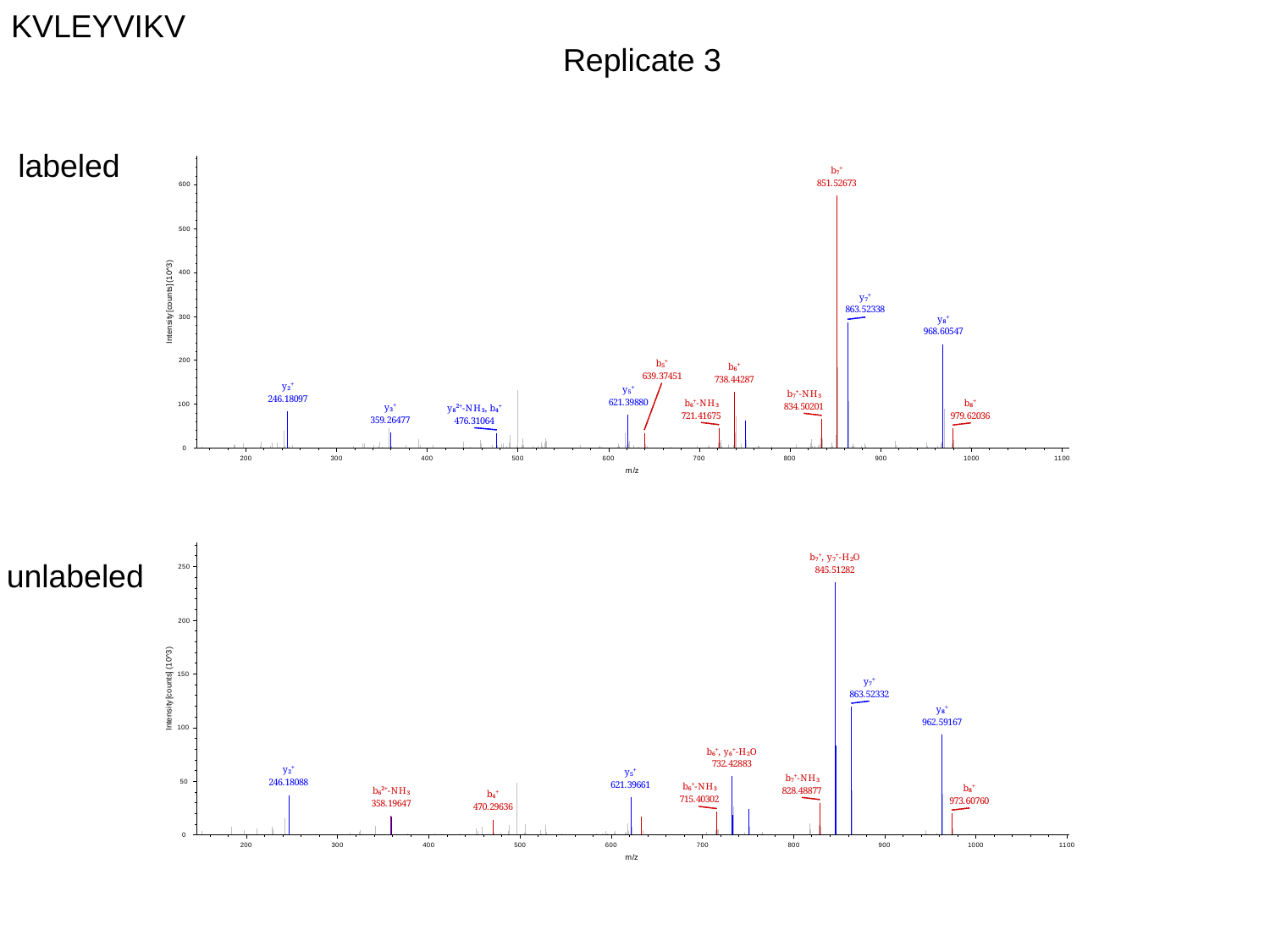

KVLEYVIKV
Replicate 3
labeled
unlabeled

### Slide 16
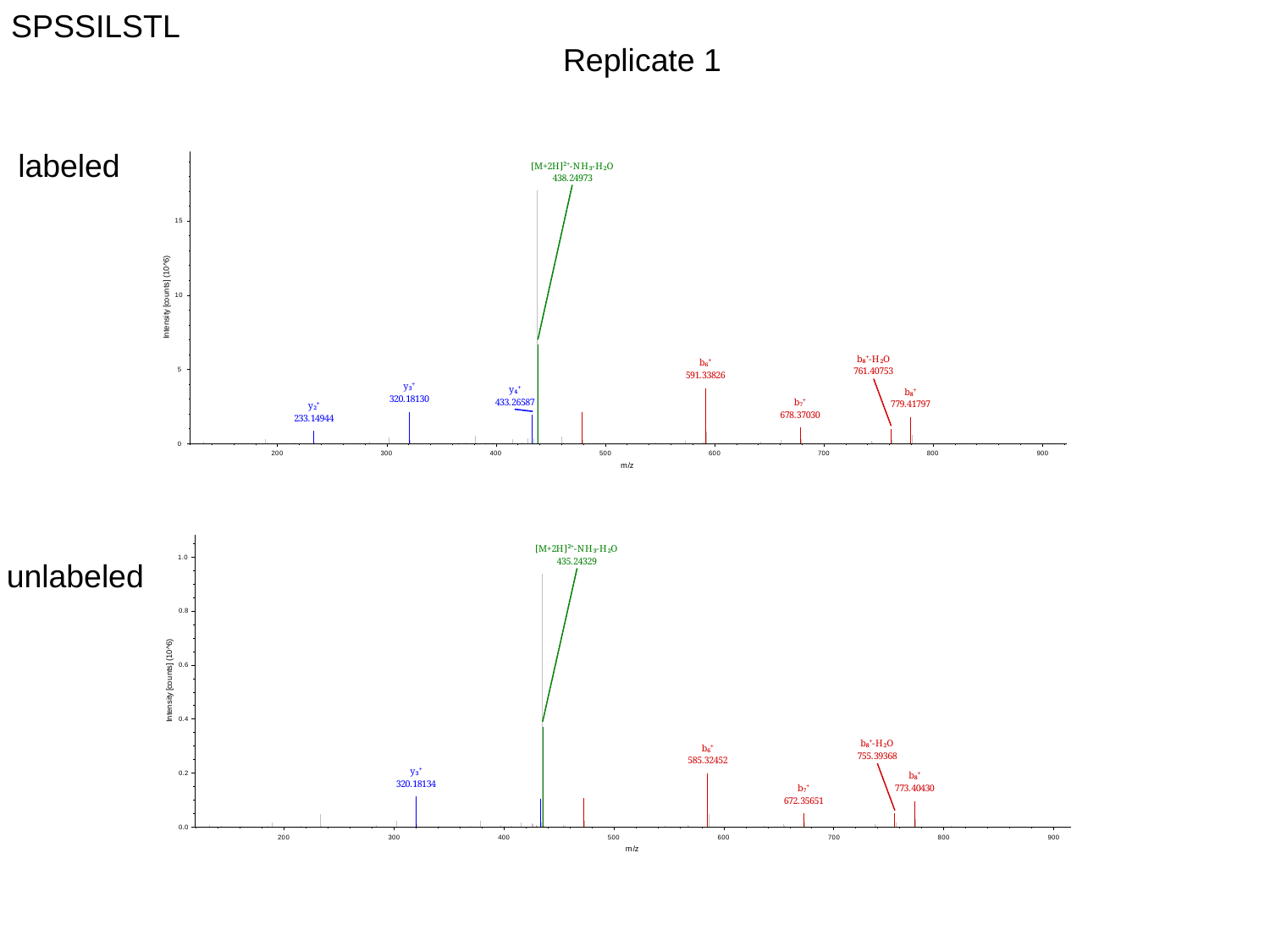

SPSSILSTL
Replicate 1
labeled
unlabeled

### Slide 17
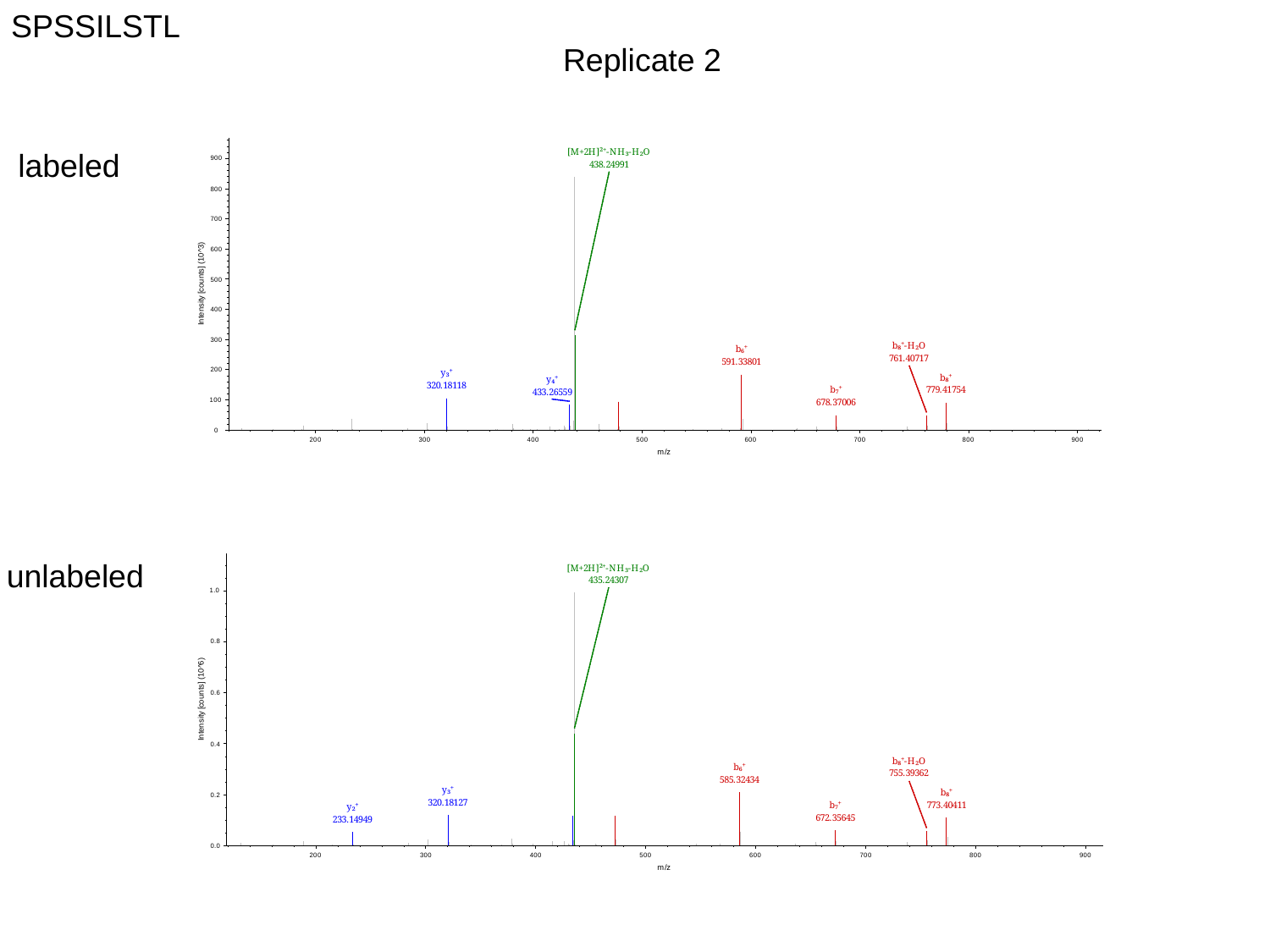

SPSSILSTL
Replicate 2
labeled
unlabeled

### Slide 18
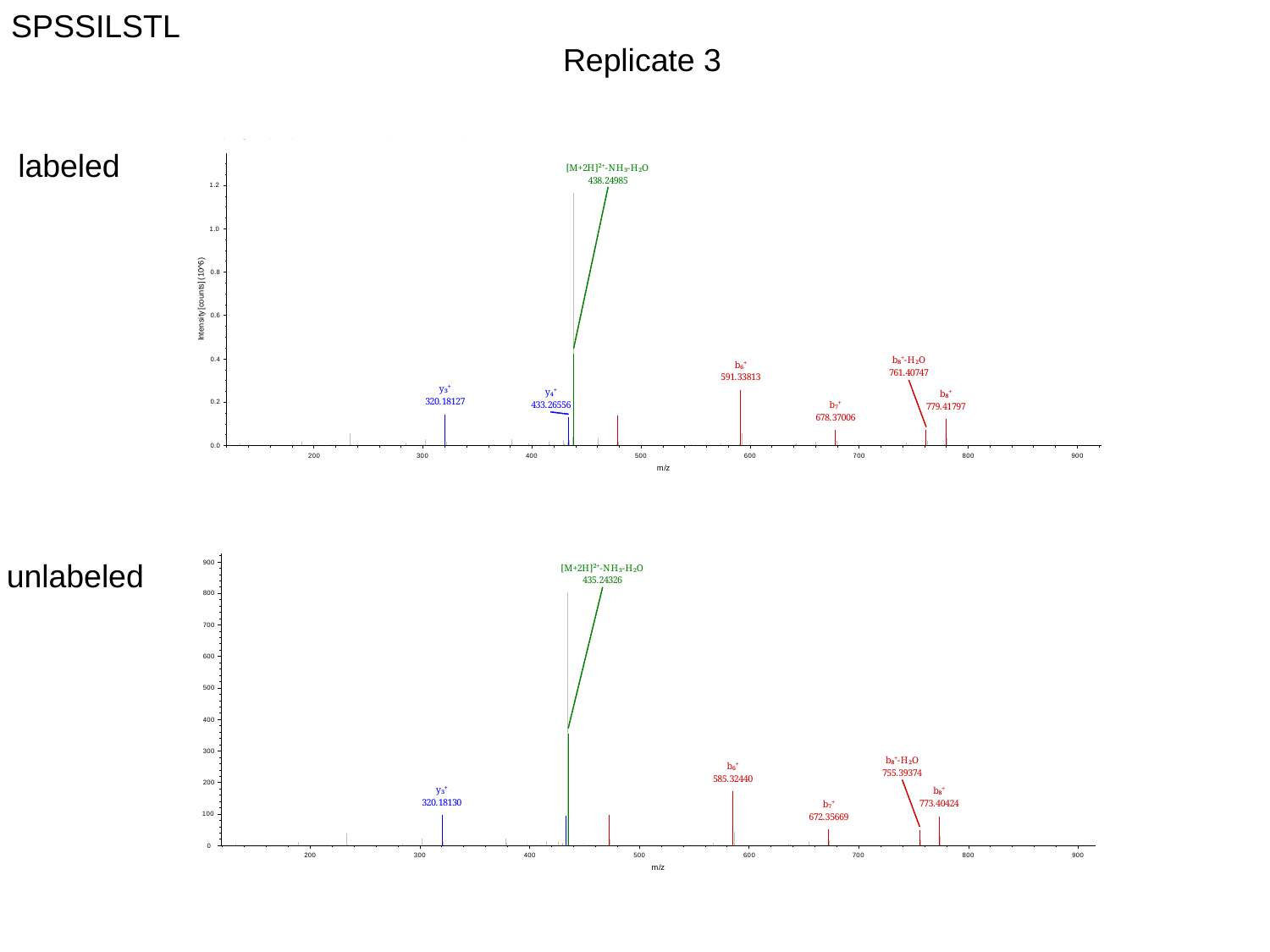

SPSSILSTL
Replicate 3
labeled
unlabeled
