## Supplemental Figure S2 for "Validation of a high-performance liquid chromatography-tandem mass spectrometry immunopeptidomics assay for the identification of HLA class I ligands suitable for pharmaceutical therapies"

### Slide 1
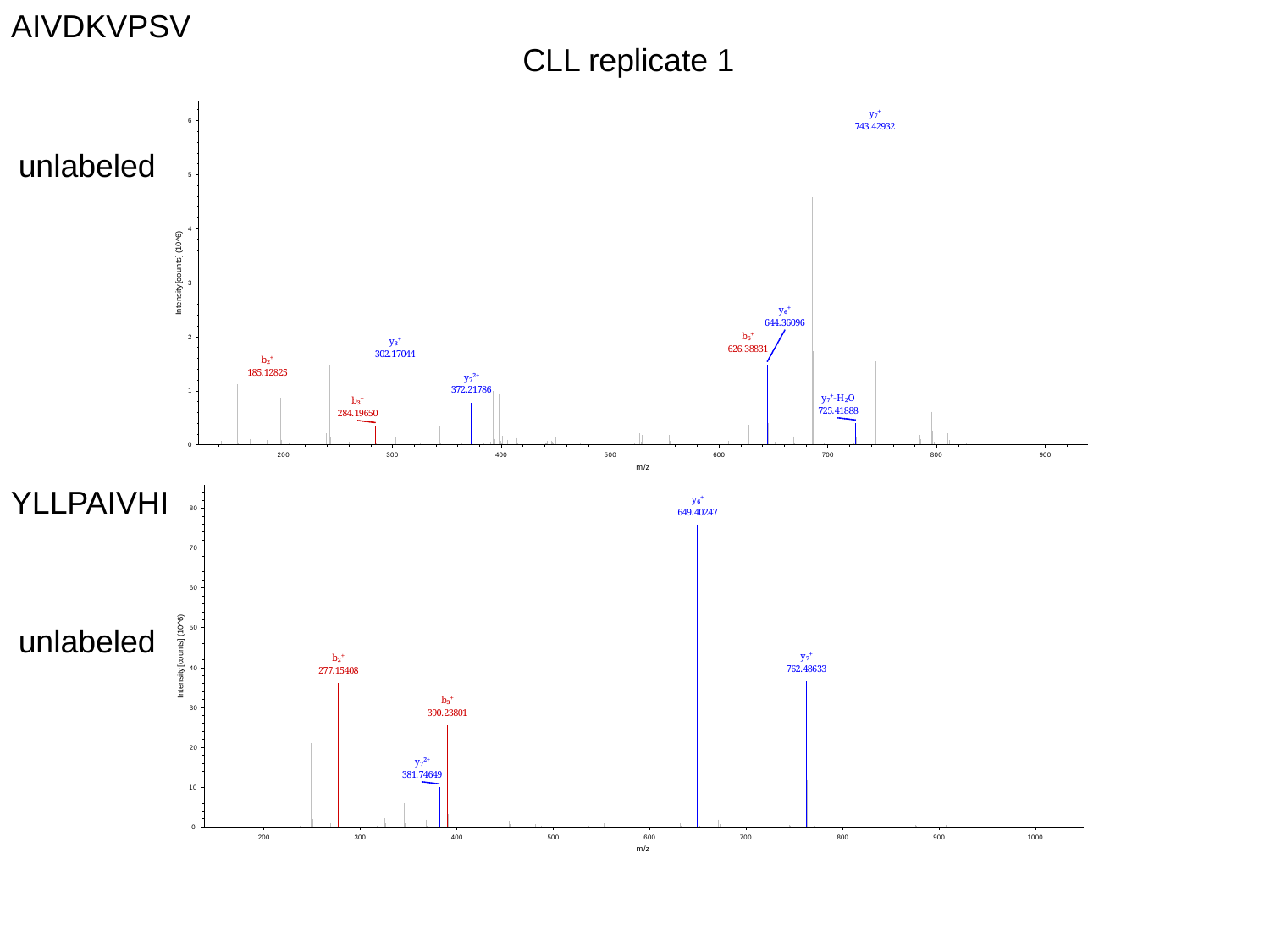

AIVDKVPSV
CLL replicate 1
unlabeled
YLLPAIVHI
unlabeled

### Slide 2
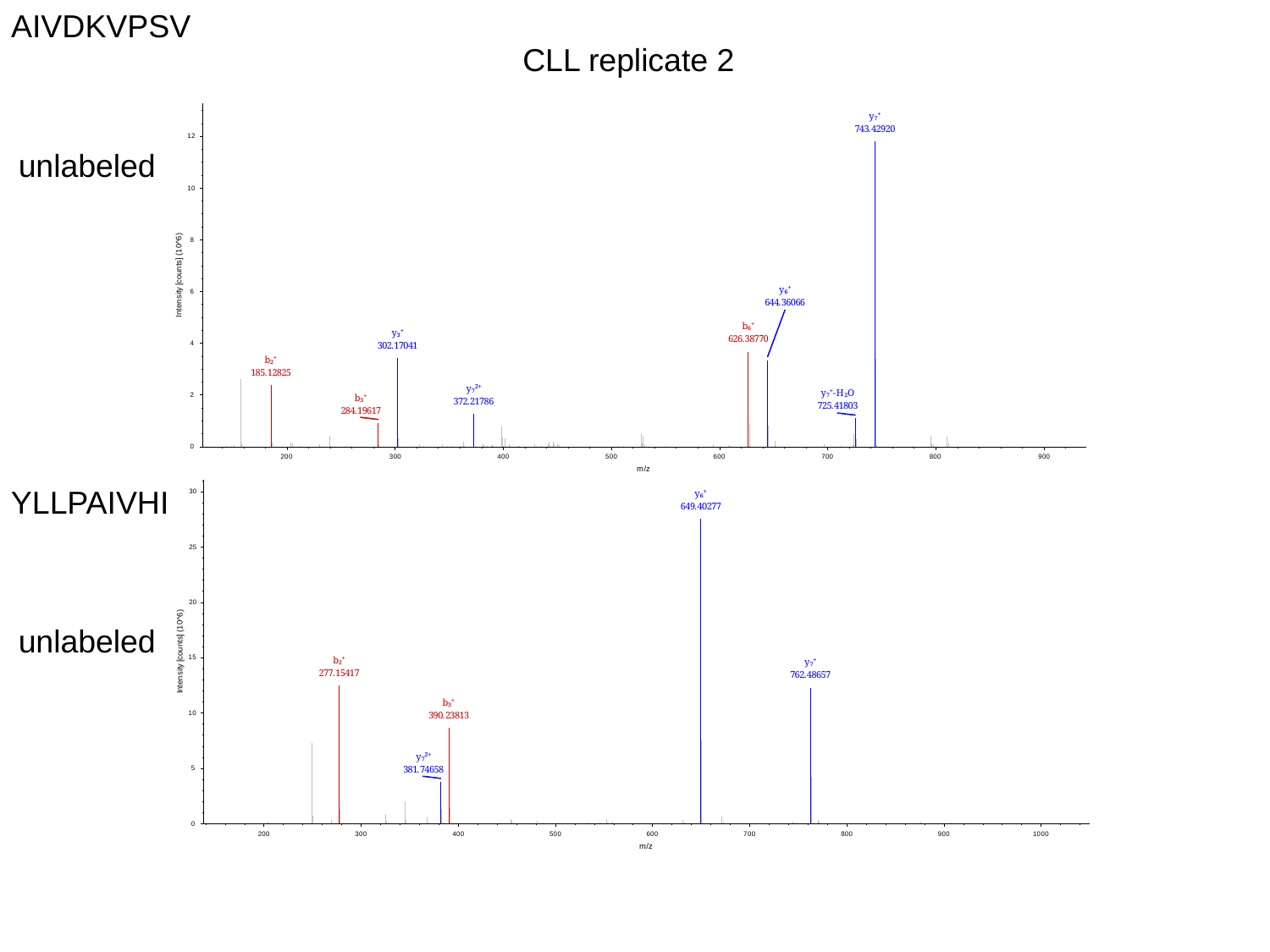

AIVDKVPSV
CLL replicate 2
unlabeled
YLLPAIVHI
unlabeled

### Slide 3
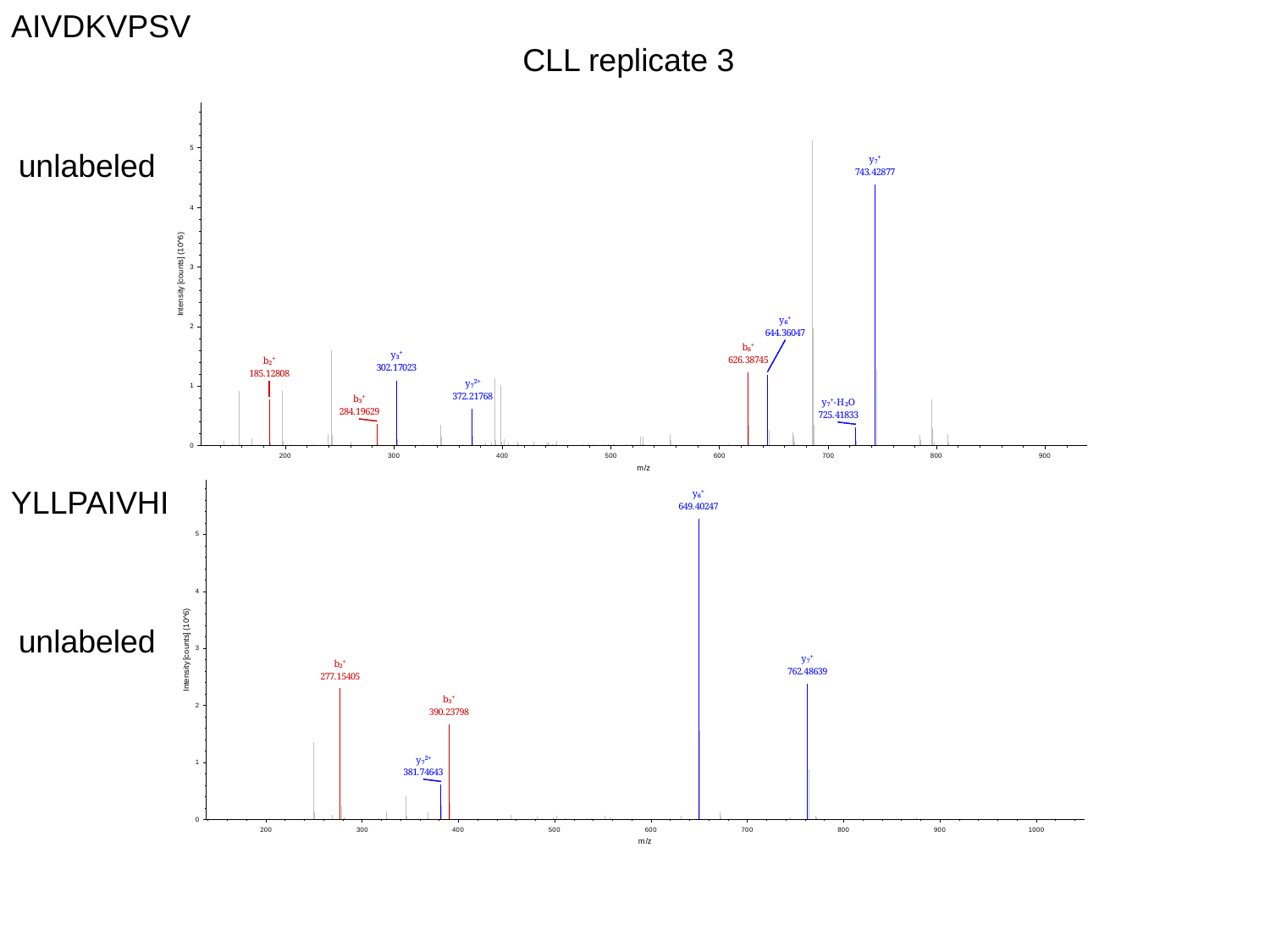

AIVDKVPSV
CLL replicate 3
unlabeled
YLLPAIVHI
unlabeled

### Slide 4
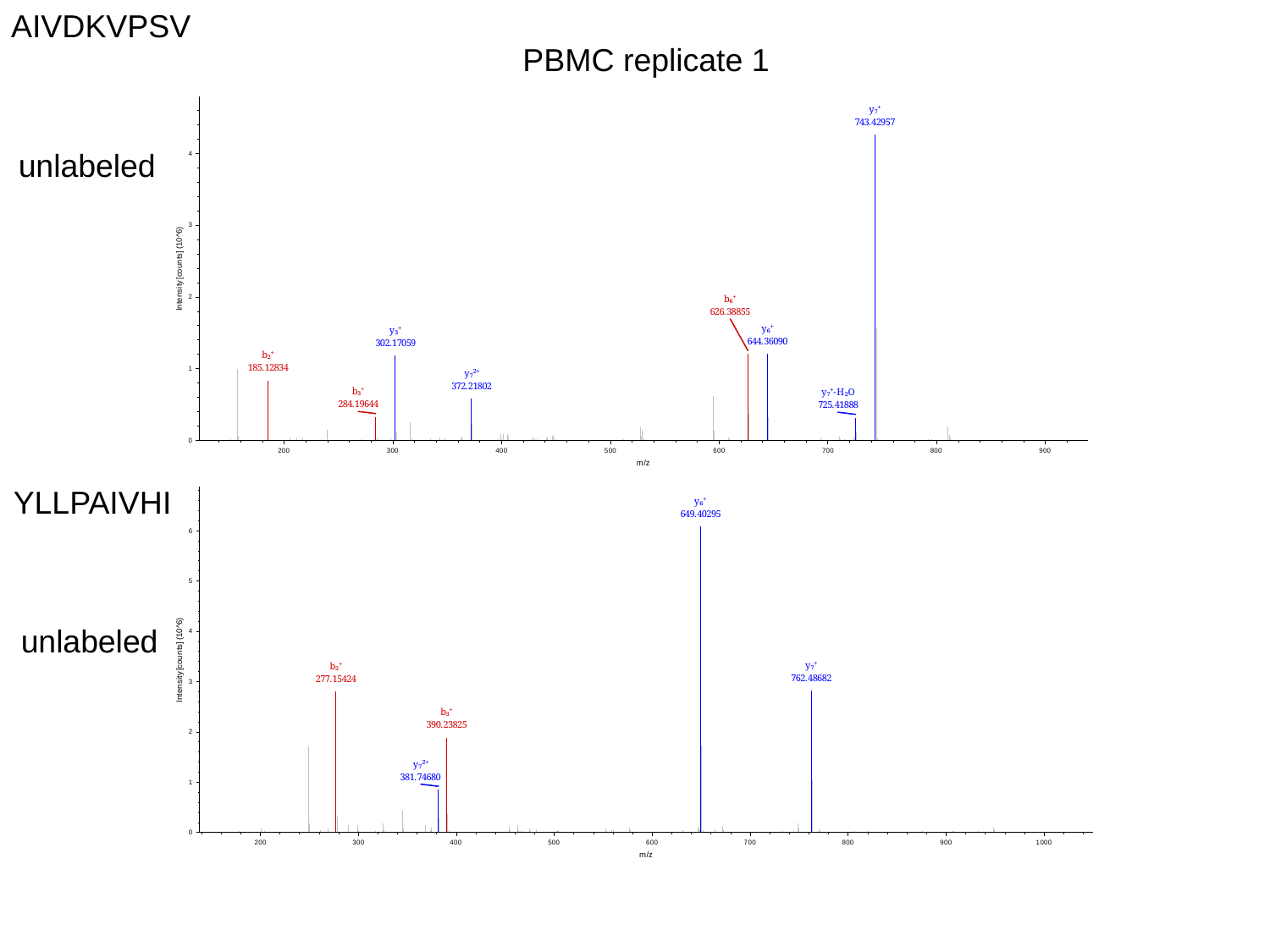

AIVDKVPSV
PBMC replicate 1
unlabeled
YLLPAIVHI
unlabeled

### Slide 5
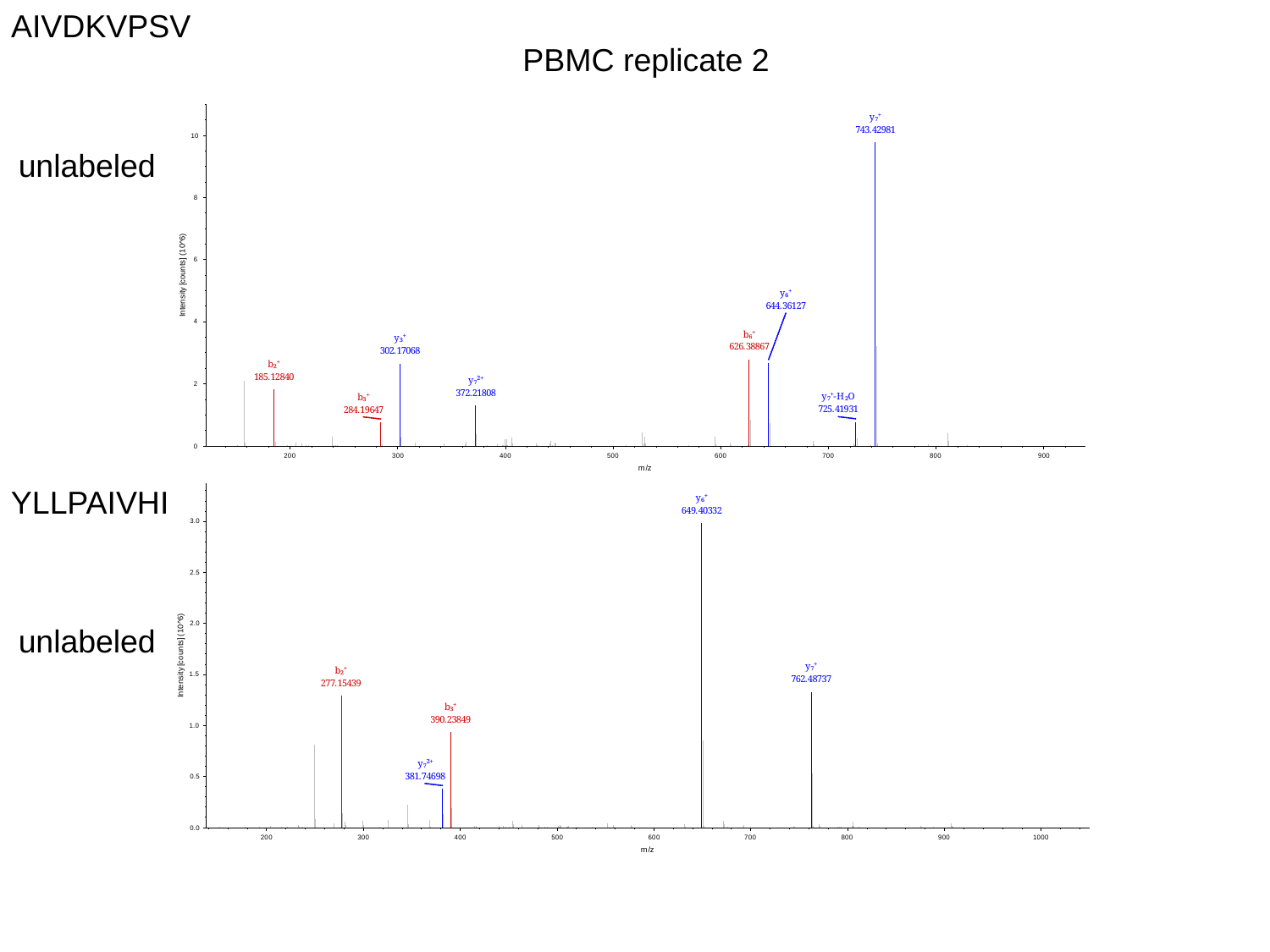

AIVDKVPSV
PBMC replicate 2
unlabeled
YLLPAIVHI
unlabeled

### Slide 6
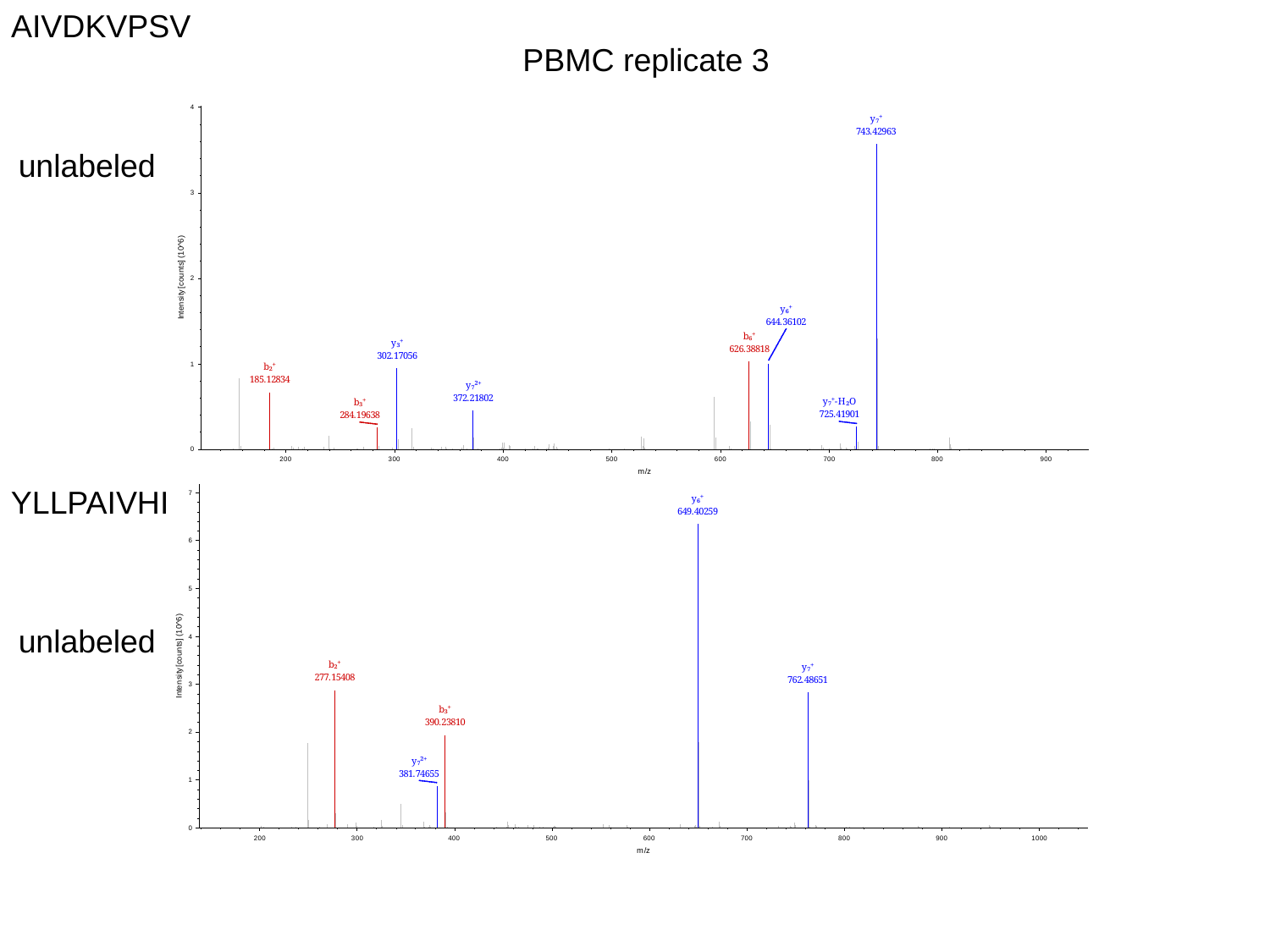

AIVDKVPSV
PBMC replicate 3
unlabeled
YLLPAIVHI
unlabeled

### Slide 7
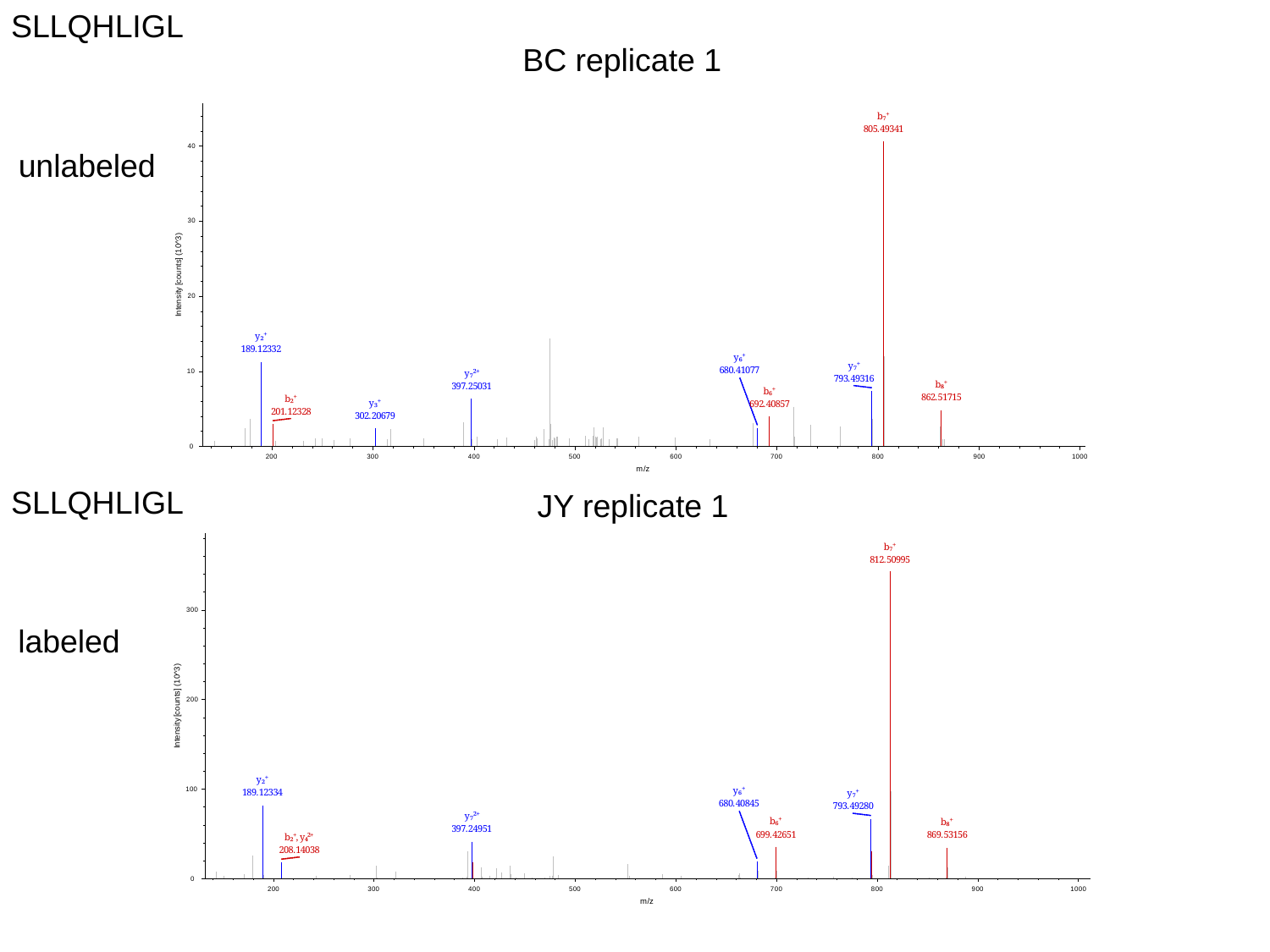

SLLQHLIGL
BC replicate 1
unlabeled
SLLQHLIGL
JY replicate 1
labeled

### Slide 8
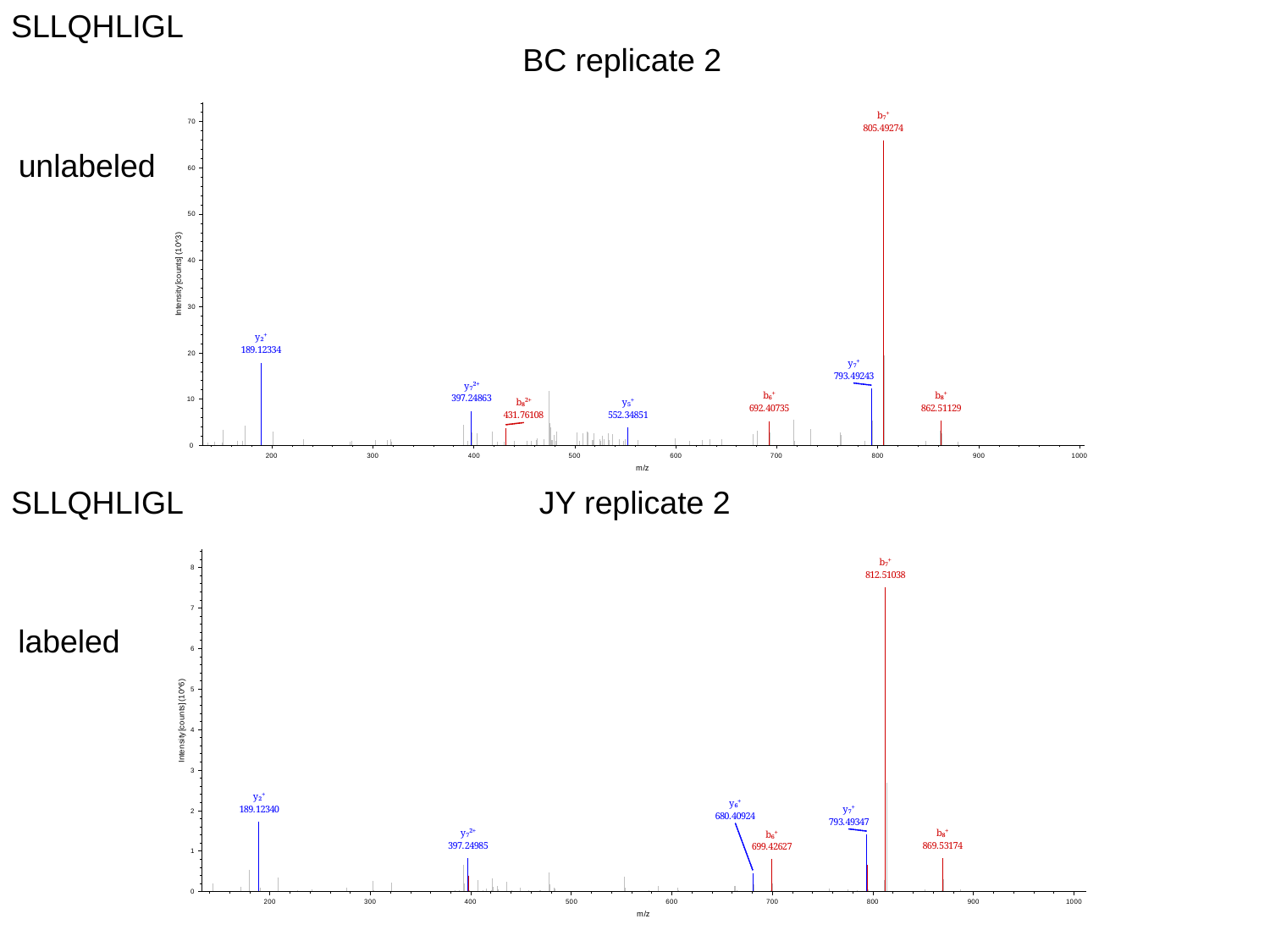

SLLQHLIGL
BC replicate 2
unlabeled
SLLQHLIGL
JY replicate 2
labeled

### Slide 9
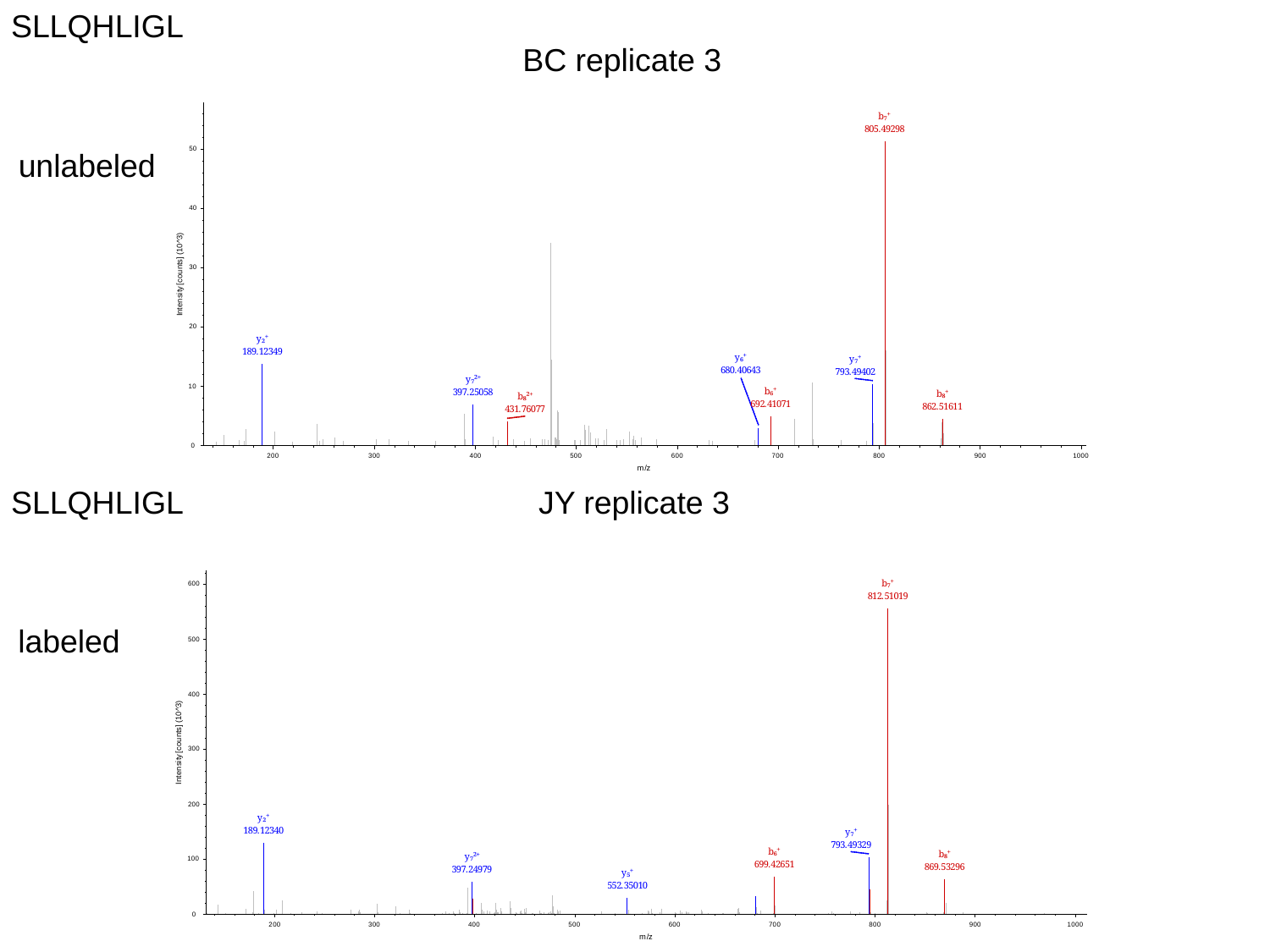

SLLQHLIGL
BC replicate 3
unlabeled
SLLQHLIGL
JY replicate 3
labeled
