## Supplemental Figure S3 for "Validation of a high-performance liquid chromatography-tandem mass spectrometry immunopeptidomics assay for the identification of HLA class I ligands suitable for pharmaceutical therapies"

### Slide 1
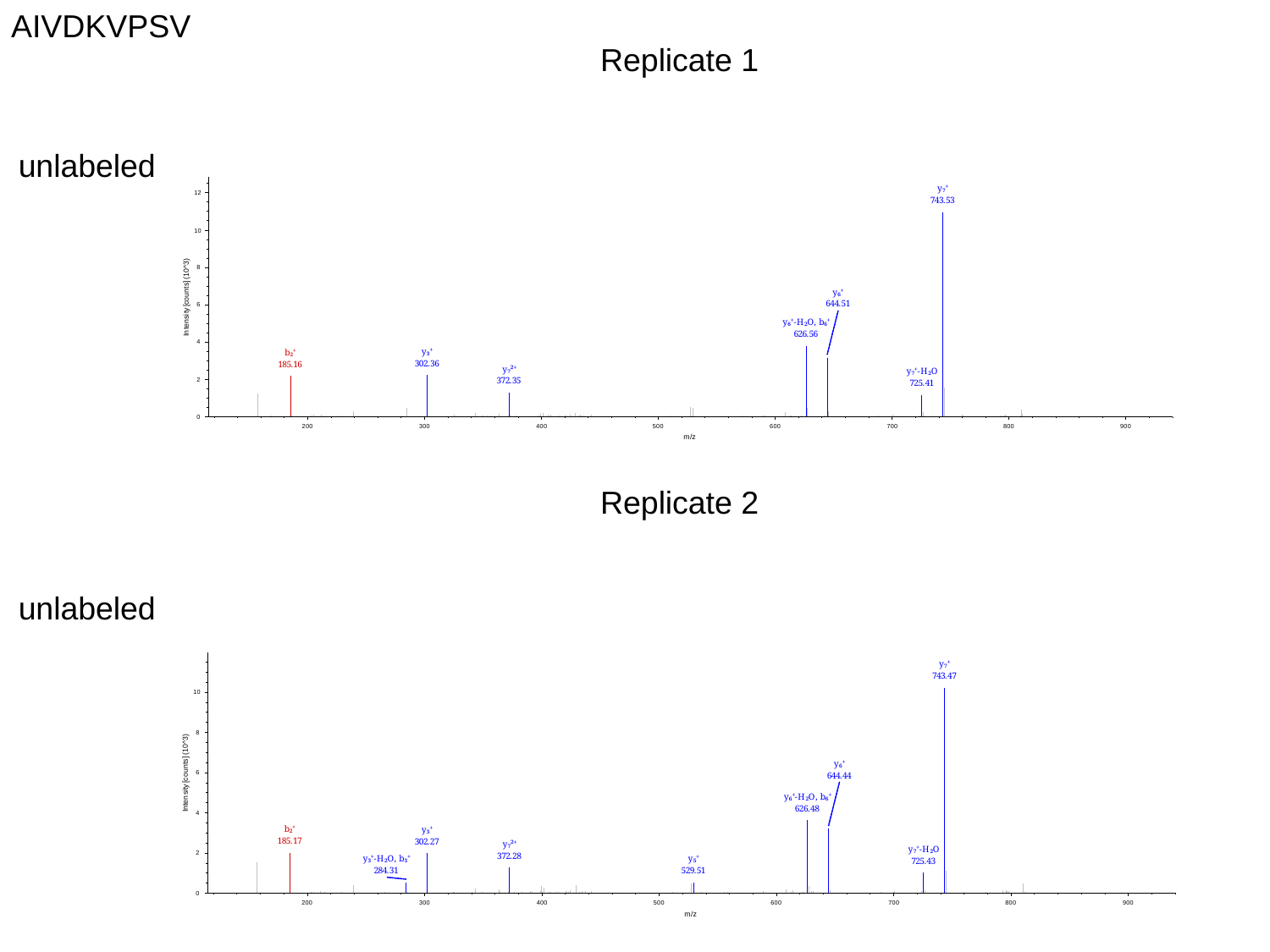

AIVDKVPSV
Replicate 1
unlabeled
Replicate 2
unlabeled

### Slide 2
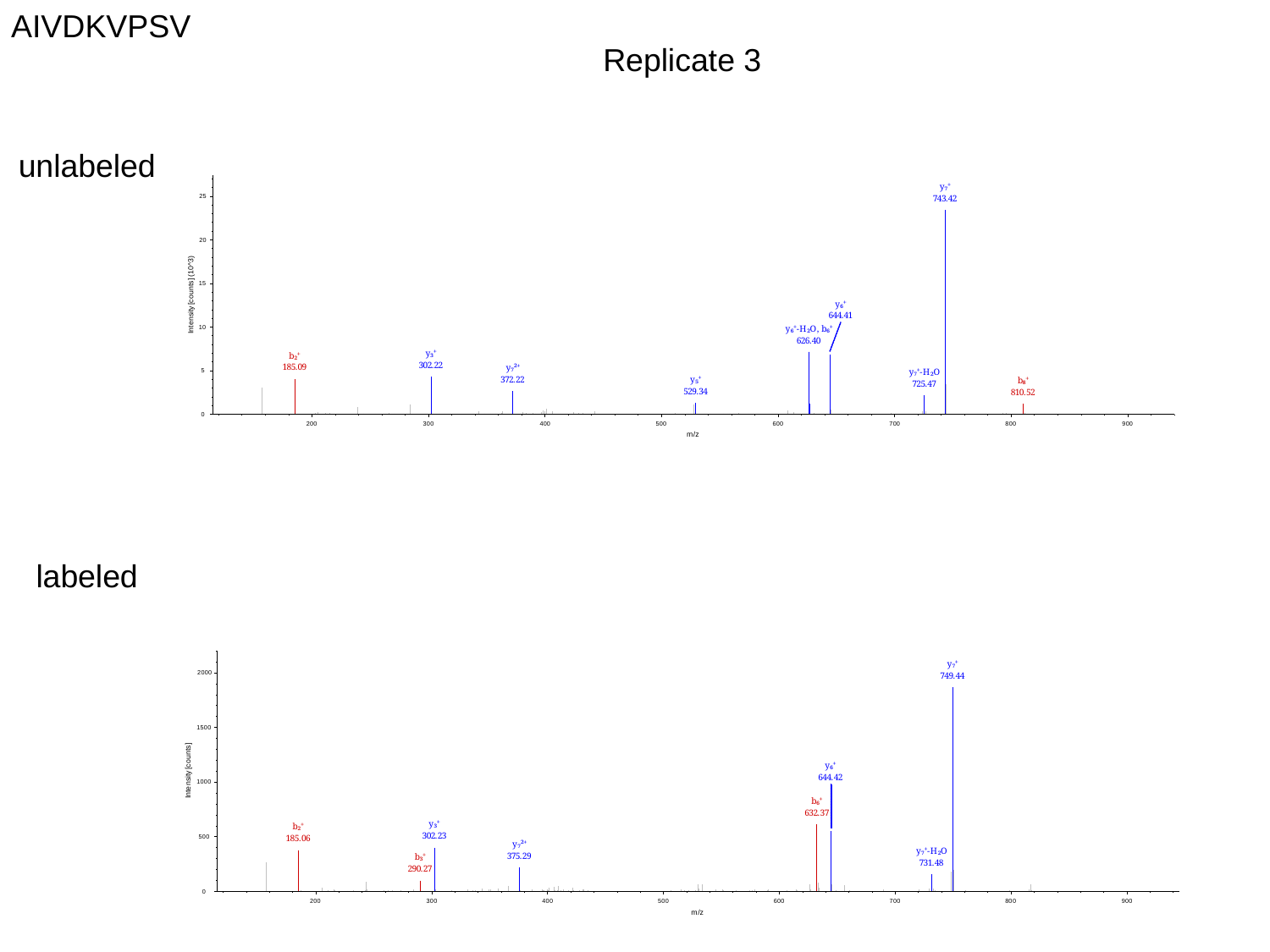

AIVDKVPSV
Replicate 3
unlabeled
labeled

### Slide 3
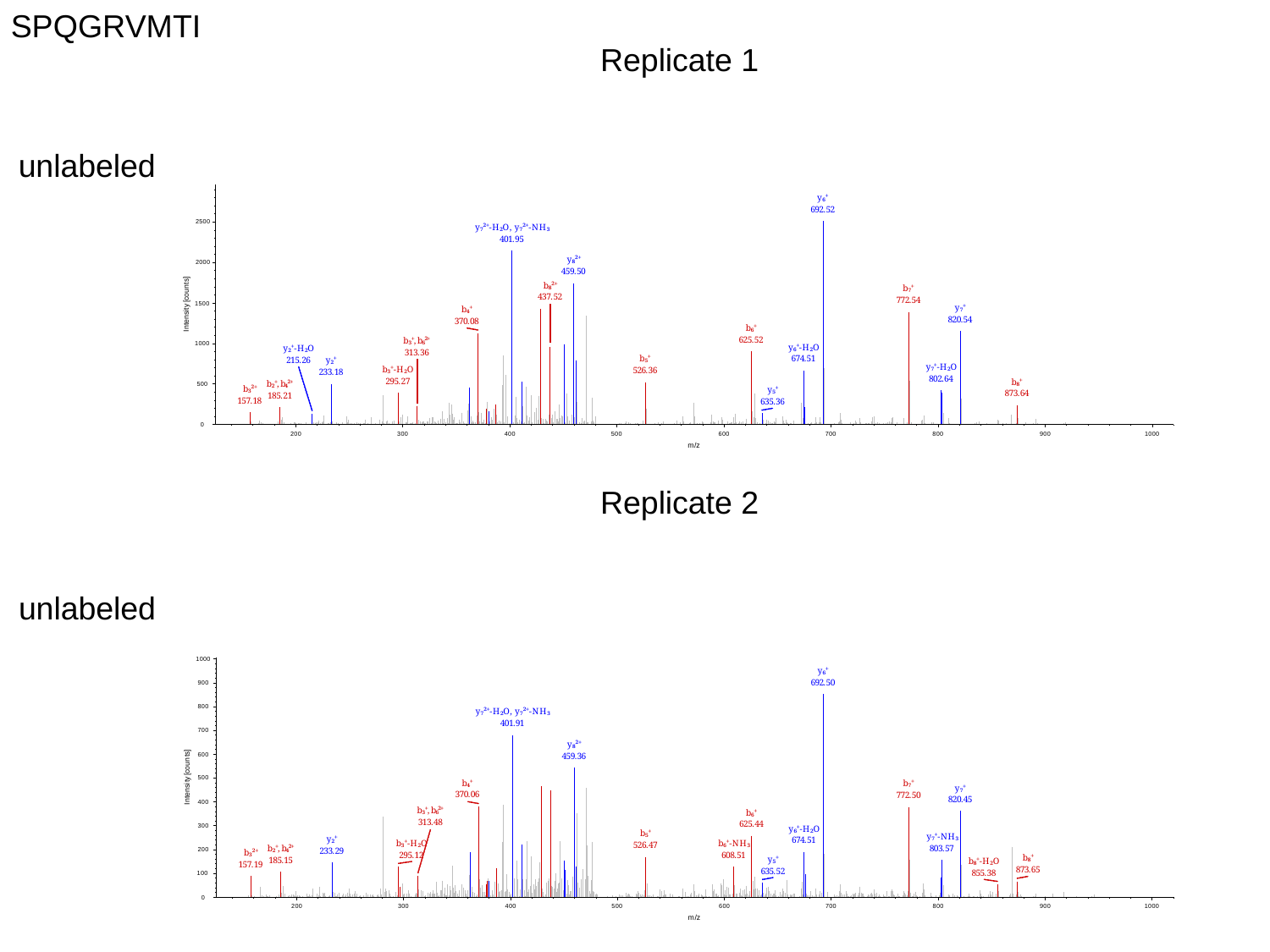

SPQGRVMTI
Replicate 1
unlabeled
Replicate 2
unlabeled

### Slide 4
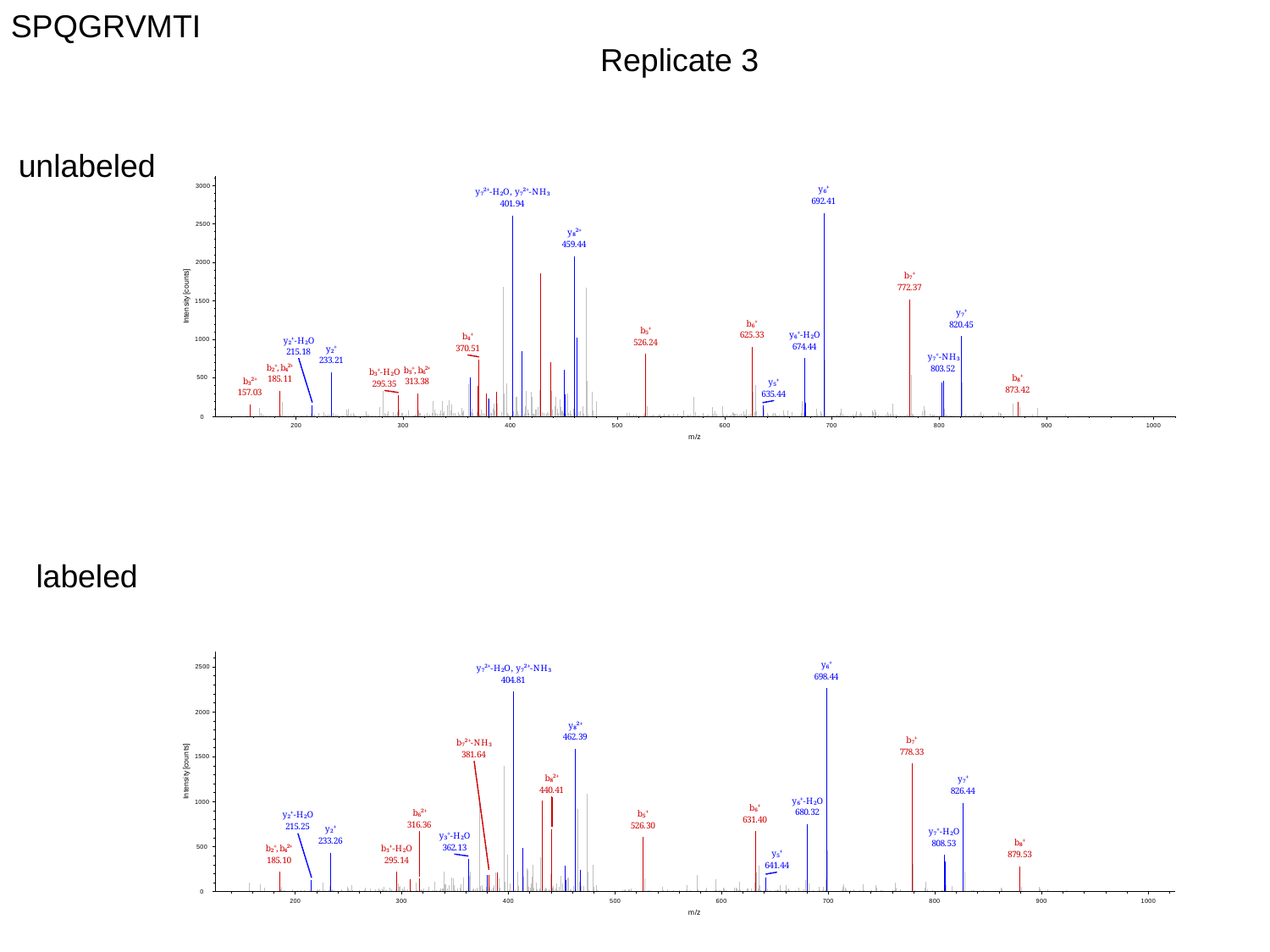

SPQGRVMTI
Replicate 3
unlabeled
labeled

### Slide 5
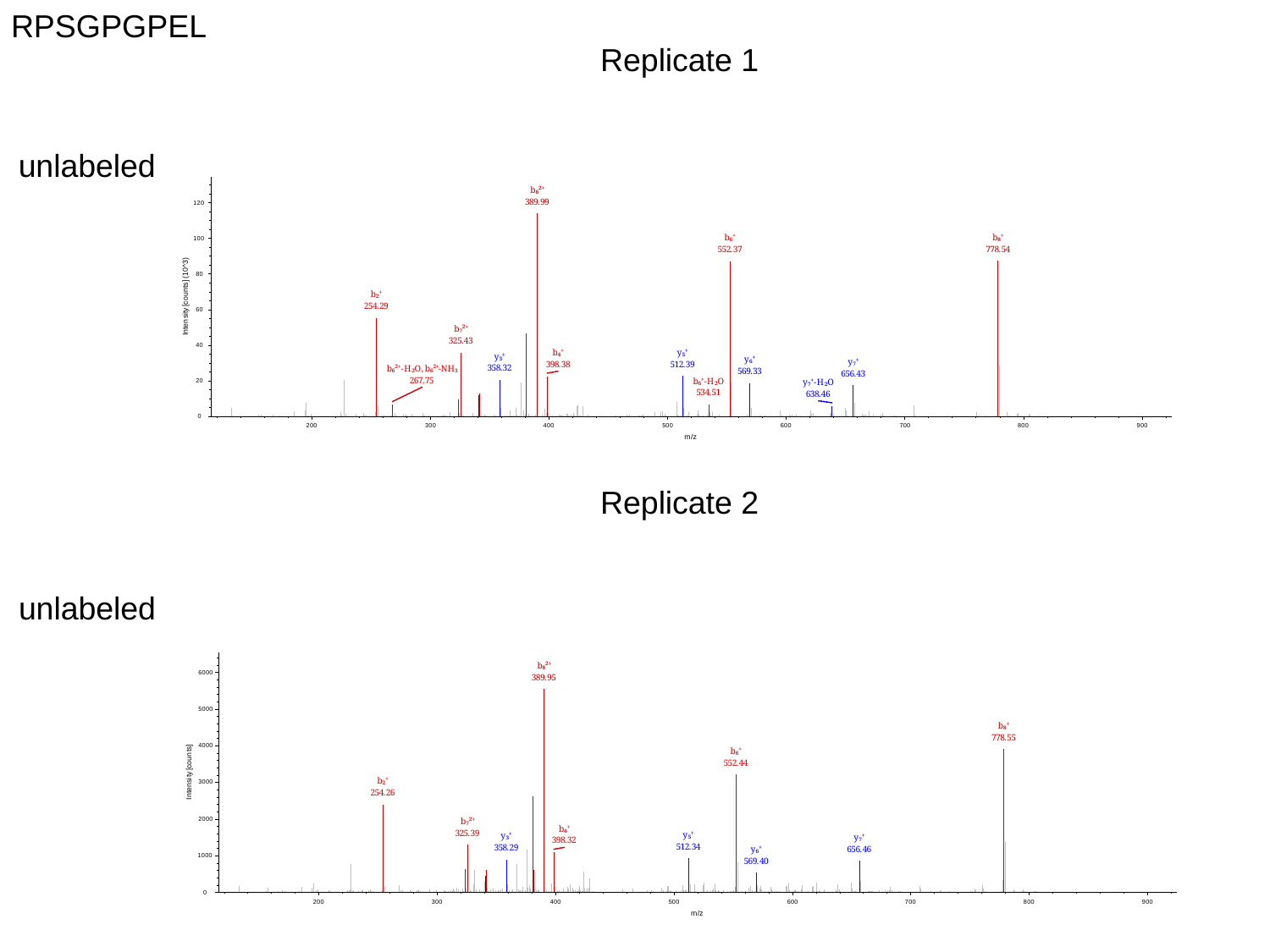

RPSGPGPEL
Replicate 1
unlabeled
Replicate 2
unlabeled

### Slide 6
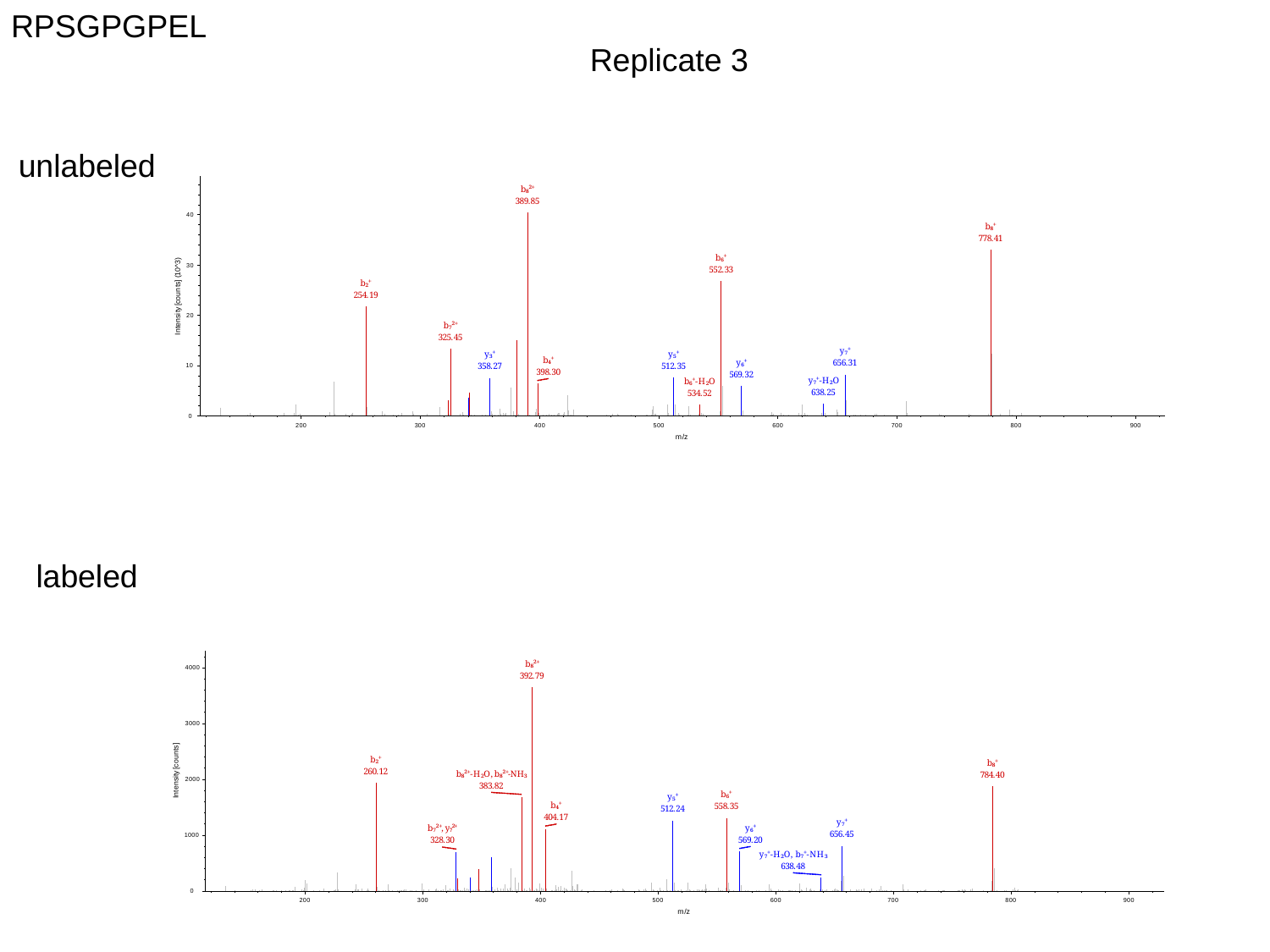

RPSGPGPEL
Replicate 3
unlabeled
labeled

### Slide 7
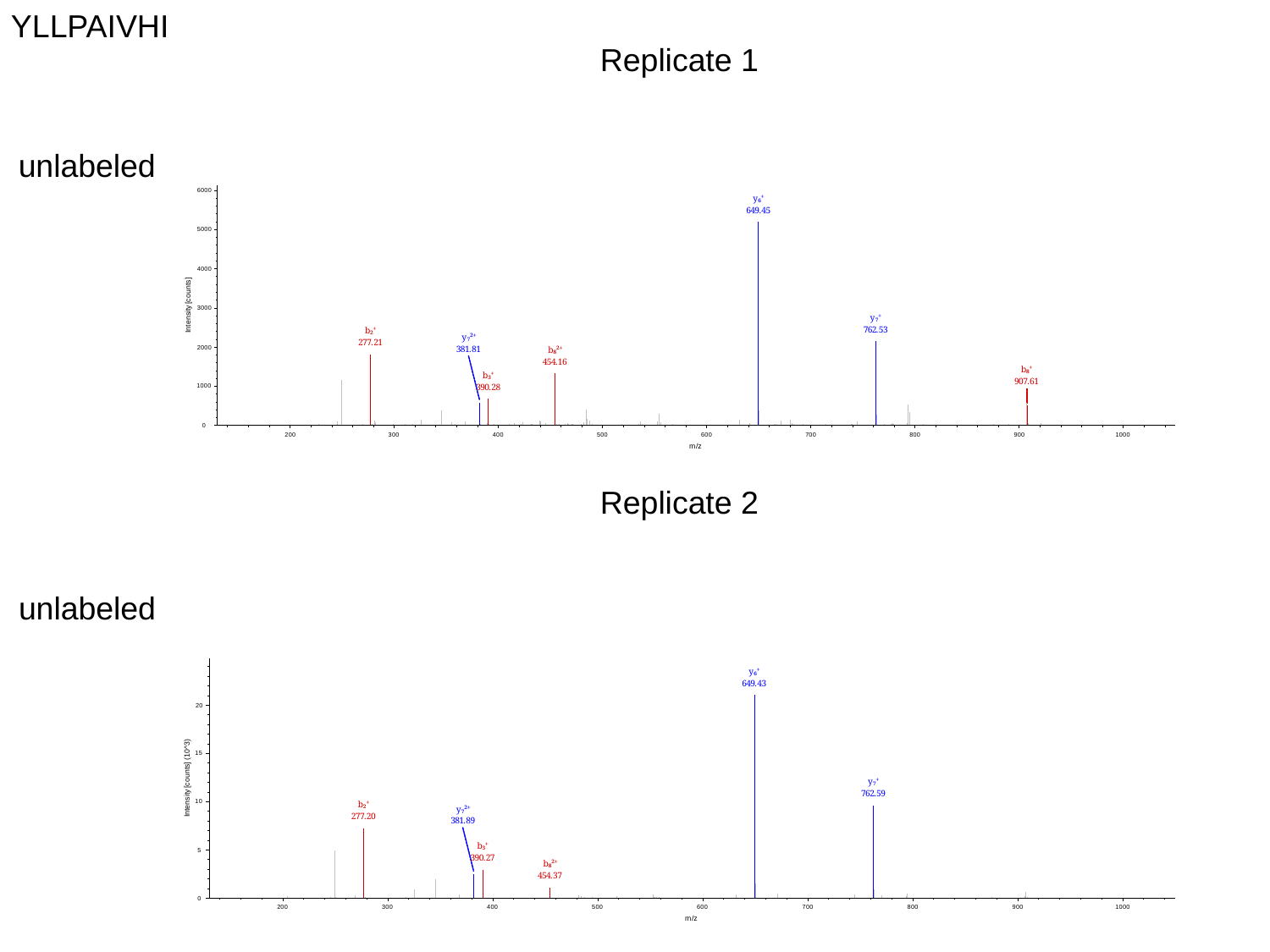

YLLPAIVHI
Replicate 1
unlabeled
Replicate 2
unlabeled

### Slide 8
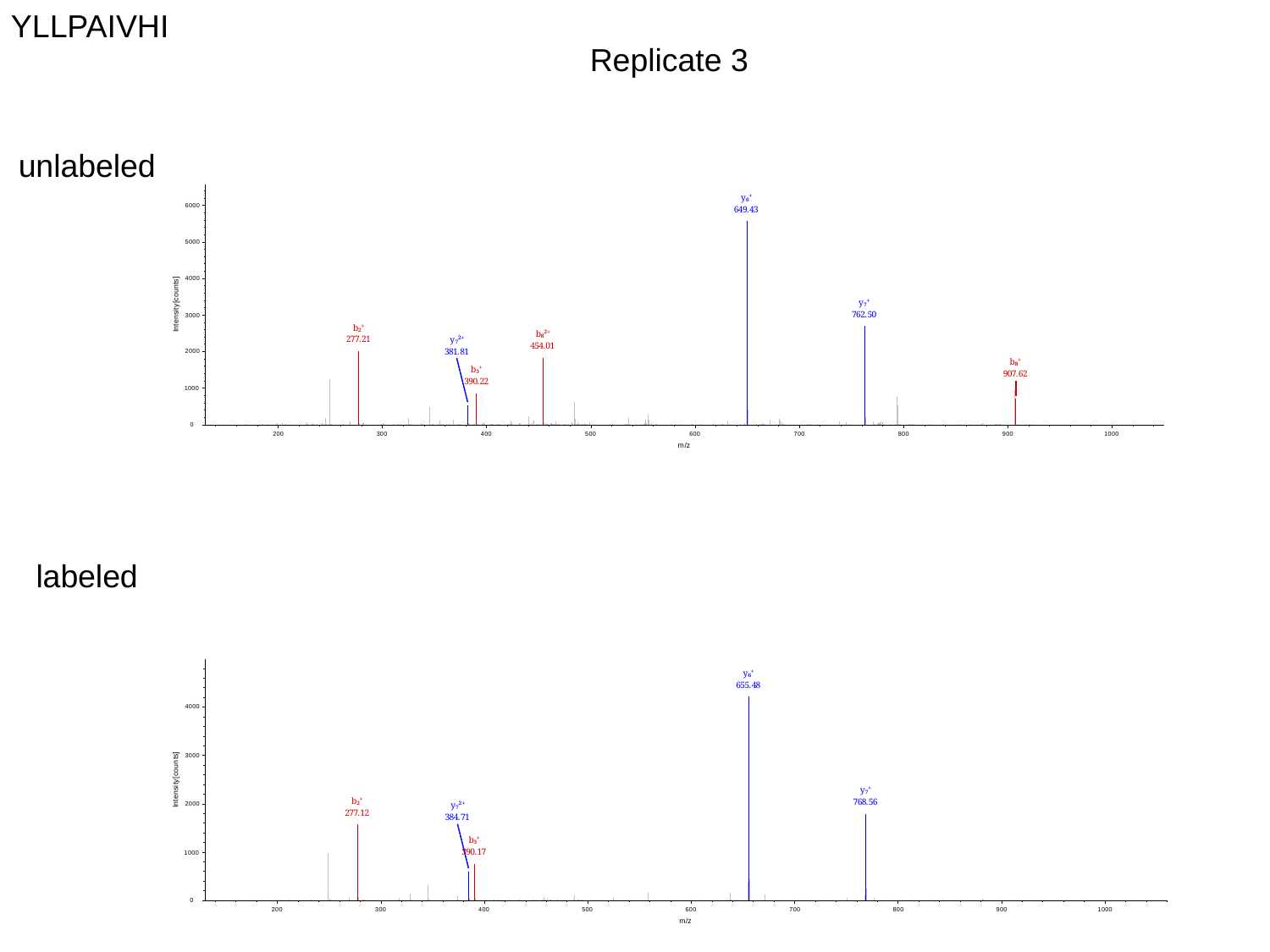

YLLPAIVHI
Replicate 3
unlabeled
labeled

### Slide 9
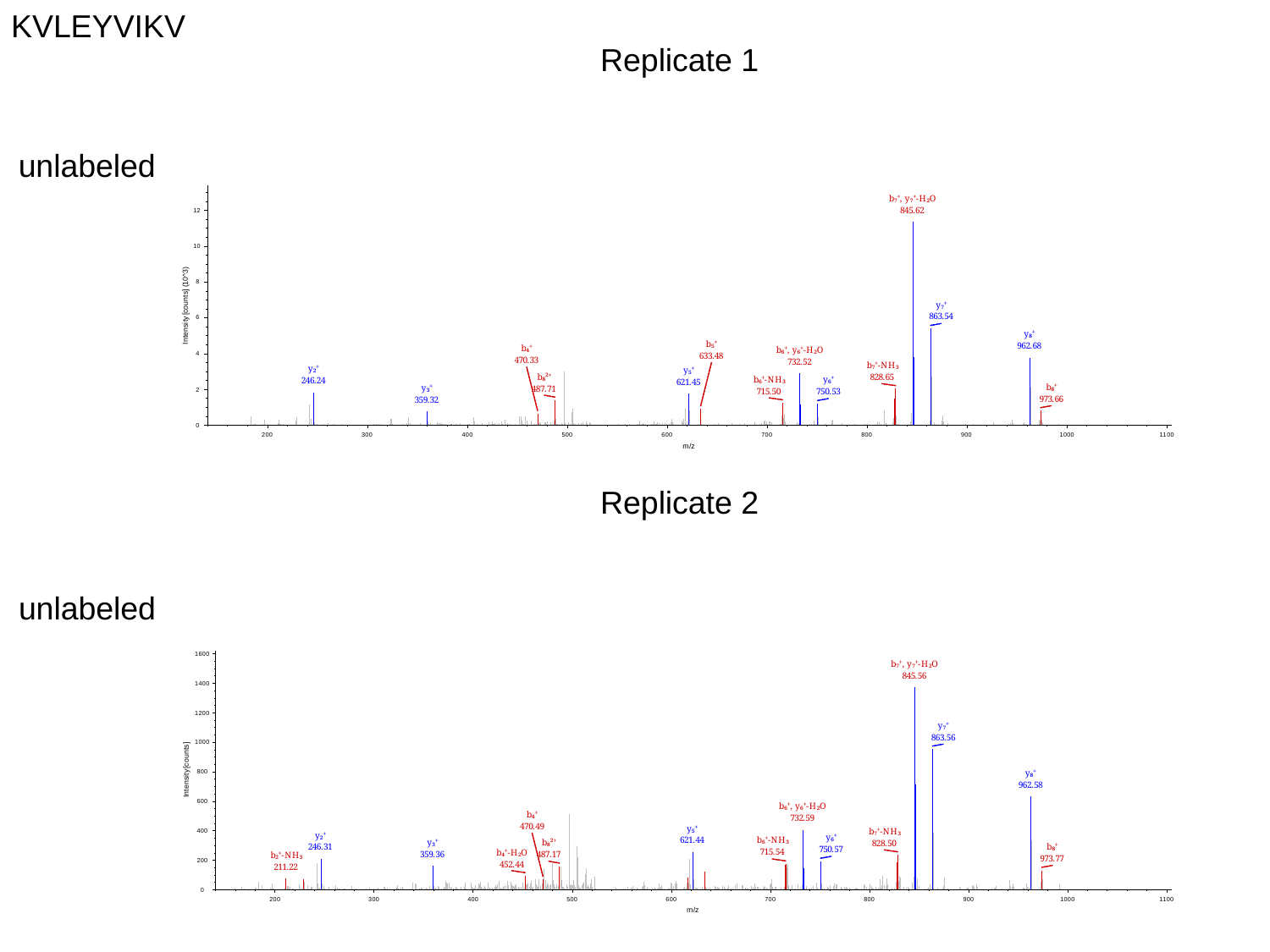

KVLEYVIKV
Replicate 1
unlabeled
Replicate 2
unlabeled

### Slide 10
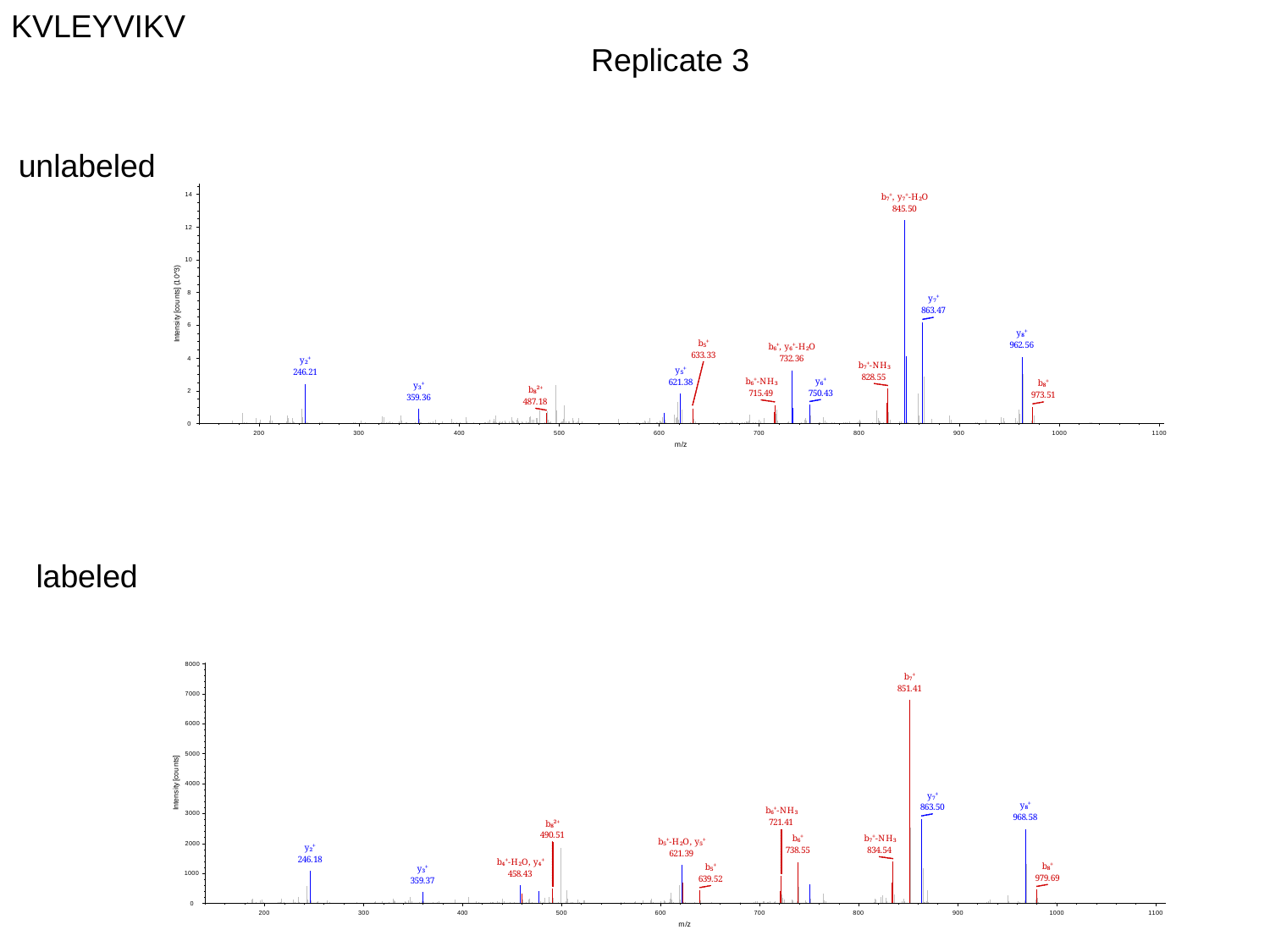

KVLEYVIKV
Replicate 3
unlabeled
labeled

### Slide 11
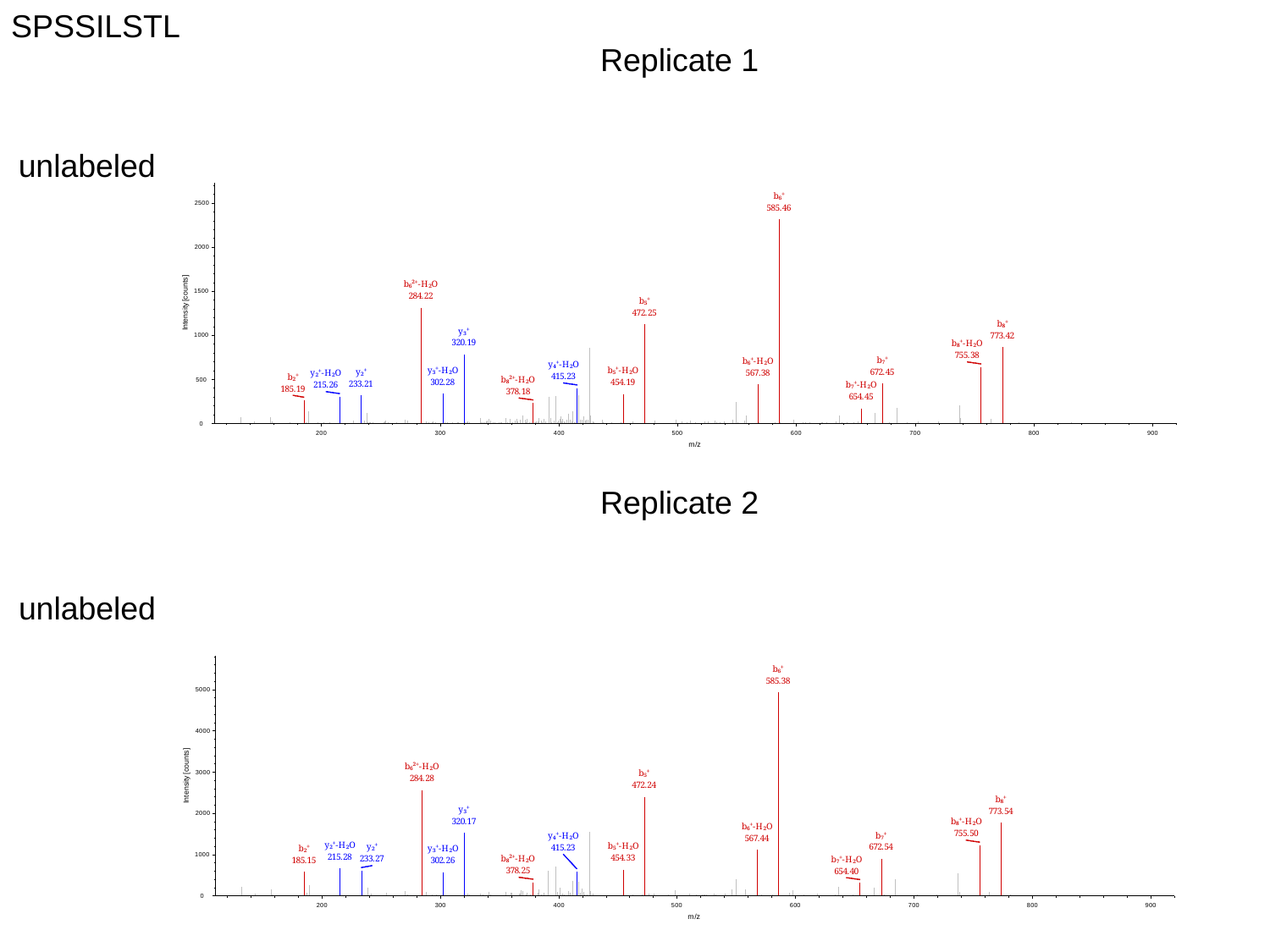

SPSSILSTL
Replicate 1
unlabeled
Replicate 2
unlabeled

### Slide 12
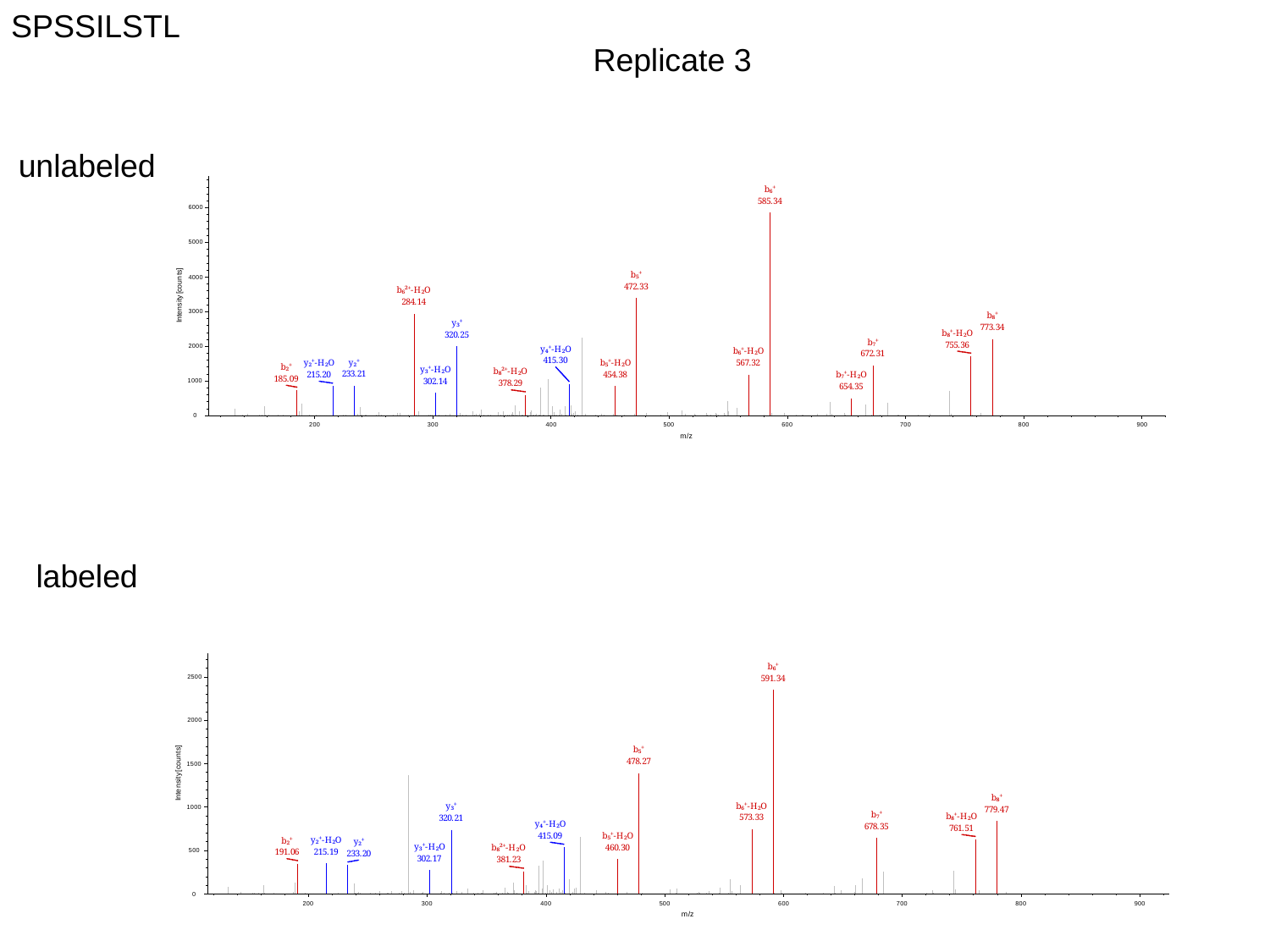

SPSSILSTL
Replicate 3
unlabeled
labeled
