## Supplemental Table S1 for "Validation of a high-performance liquid chromatography-tandem mass spectrometry immunopeptidomics assay for the identification of HLA class I ligands suitable for pharmaceutical therapies"

| **Batch** | **Sequence** | **Molecular weight** | **pI** | **GRAVY** |
| --- | --- | --- | --- | --- |
| 174121 | DG**P**SSAPATPTK | 1,128.20 | 5.84 | -1.00 |
| 174013 | SLNSN**V**YDV | 1,010.07 | 3.80 | -0.13 |
| 174014 | FPHLPGKTF**V**Y | 1,305.54 | 8.60 | 0.08 |
| 174015 | S**V**LTPLLLR | 1,011.27 | 9.47 | 1.31 |
| 174085 | RYQA**L**FHDF | 1,196.33 | 6.74 | -0.53 |
| 174122 | G**P**SSAPATPTK | 1,013.12 | 8.75 | -0.77 |
| 174123 | F**P**HLPGKTF | 1,043.23 | 8.76 | -0.22 |
| 174124 | FLL**P**AGWIL | 1,029.29 | 5.52 | 1.96 |
| 174125 | LL**P**AGWIL | 882.11 | 5.52 | 1.85 |
| 174126 | T**P**LLLRGL | 882.11 | 9.41 | 1.00 |
| 174017 | AI**V**DKVPSV | 927.11 | 5.88 | 1.01 |
| 174018 | YLLPAI**V**HI | 1,038.30 | 6.74 | 1.83 |
| 174019 | **V**YVVGTAHF | 992.14 | 6.71 | 1.29 |
| 174020 | GTY**V**SSVPR | 965.07 | 8.75 | -0.19 |
| 174021 | TYQE**V**AQKF | 1,113.24 | 5.66 | -0.84 |
| 174022 | SPQGR**V**MTI | 988.17 | 9.47 | -0.10 |
| 174086 | ITDSAGHI**L**Y | 1,089.21 | 5.08 | 0.47 |
| 174087 | RVYGG**L**TTK | 994.16 | 9.99 | -0.43 |
| 174088 | A**L**KTGIVAK | 900.13 | 10.00 | 0.80 |
| 174089 | SV**L**NLVIVK | 984.25 | 8.47 | 1.83 |
| 174090 | D**L**IIKGISV | 957.18 | 5.84 | 1.43 |
| 174127 | NTDS**P**LRY | 965.03 | 5.84 | -1.51 |
| 174128 | R**P**SGPGPEL | 909.01 | 6.00 | -1.18 |
| 174023 | DLKEKKE**V**V | 1,087.28 | 6.18 | -1.11 |
| 174024 | TLHDQ**V**HLL | 1,075.23 | 5.90 | 0.17 |
| 174061 | N**V**GGLIGTPK | 955.12 | 8.75 | 0.16 |
| 174062 | FYFPTPT**V**L | 1,084.28 | 5.52 | 0.86 |
| 174063 | RSYHLQI**V**TK | 1,244.46 | 9.99 | -0.54 |
| 174064 | NPKAFFS**V**L | 1,022.21 | 8.75 | 0.62 |
| 174065 | NPS**V**REFVL | 1,060.22 | 6.00 | 0.12 |
| 174066 | **V**LVDQSWVL | 1,058.24 | 3.80 | 1.28 |
| 174068 | APDAKSF**V**L | 947.10 | 5.88 | 0.51 |
| 174069 | K**V**LEYVIKV | 1,090.37 | 8.50 | 0.92 |
| 174070 | G**V**YDGREHTV | 1,132.20 | 5.32 | -0.91 |
| 174071 | K**V**LEHVVRV | 1,078.32 | 8.75 | 0.61 |
| 174072 | ALDEK**V**AEL | 987.12 | 4.14 | 0.11 |
| 174073 | G**V**YDGEEHSV | 1,091.10 | 4.13 | -0.82 |
| 174074 | F**V**YGEPREL | 1,109.25 | 4.53 | -0.44 |
| 174075 | NA**V**GVYAGR | 906.01 | 8.75 | 0.21 |
| 174076 | **V**WSDVTPLTF | 1,164.32 | 3.80 | 0.68 |
| 174077 | A**V**LPLTVAEVQK | 1,267.53 | 6.05 | 0.88 |
| 174078 | R**V**RELAVAL | 1,026.25 | 9.60 | 0.79 |
| 174079 | HLTE**V**YPEL | 1,100.24 | 4.51 | -0.22 |
| 174080 | S**V**LADLVTTK | 1,046.23 | 5.55 | 0.82 |
| 174081 | IPFSNPR**V**L | 1,042.25 | 9.75 | 0.37 |
| 174082 | **V**LYGPAGLGK | 974.17 | 8.56 | 0.56 |
| 174083 | SPS**V**SQLSVL | 1,016.16 | 5.24 | 0.77 |
| 174084 | **V**LYPVPLESY | 1,179.38 | 4.00 | 0.59 |
| 174092 | T**L**LKALLEI | 1,013.29 | 5.66 | 1.49 |
| 174093 | A**L**REEEEGV | 1,031.09 | 4.09 | -1.01 |
| 174094 | S**L**LKFLAKV | 1,018.31 | 10.00 | 1.29 |
| 174095 | **L**IHFLLLK | 996.30 | 8.76 | 1.93 |
| 174096 | G**L**YDGREHSV | 1,132.20 | 5.32 | -0.96 |
| 174129 | LAQ**P**PSGQR | 953.07 | 9.75 | -1.14 |
| 174130 | F**P**SLREAAL | 1,003.17 | 6.00 | 0.40 |
| 174132 | S**P**SKAFASL | 907.03 | 8.47 | 0.26 |
| 174205 | RLLDS**V**SRL | 1,058.25 | 9.60 | 0.17 |
| 174206 | TYSEKTT**L**F | 1,089.21 | 5.66 | -0.56 |
| 174207 | S**L**LQHLIGL | 993.21 | 6.46 | 1.31 |
| 174208 | S**P**SSILSTL | 904.03 | 5.24 | 0.73 |
| 174210 | VLLAGFK**P**PL | 1,054.34 | 8.72 | 1.27 |
| 174211 | LYL**P**KSWTI | 1,120.36 | 8.59 | 0.32 |
