## Supplemental Table S2 for "Validation of a high-performance liquid chromatography-tandem mass spectrometry immunopeptidomics assay for the identification of HLA class I ligands suitable for pharmaceutical therapies"

| **Sample** | **Purpose** | **Condition/ number** | **Technical replicate** | **Raw file name** |
| --- | --- | --- | --- | --- |
| JY cell line | Precision | 1 (day 0) | 1 | JY_repeatability_1_1 |
| JY cell line | Precision | 1 (day 0) | 2 | JY_repeatability_1_2 |
| JY cell line | Precision | 1 (day 0) | 3 | JY_repeatability_1_3 |
| JY cell line | Precision | 2 (day 0) | 1 | JY_repeatability_2_1 |
| JY cell line | Precision | 2 (day 0) | 2 | JY_repeatability_2_2 |
| JY cell line | Precision | 2 (day 0) | 3 | JY_repeatability_2_3 |
| JY cell line | Precision | 3 (day 0) | 1 | JY_repeatability_3_1 |
| JY cell line | Precision | 3 (day 0) | 2 | JY_repeatability_3_2 |
| JY cell line | Precision | 3 (day 0) | 3 | JY_repeatability_3_3 |
| JY cell line | Precision | 1 (day 7) | 1 | JY_intermediate_precision_1_1 |
| JY cell line | Precision | 1 (day 7) | 2 | JY_intermediate_precision_1_2 |
| JY cell line | Precision | 1 (day 7) | 3 | JY_intermediate_precision_1_3 |
| JY cell line | Precision | 2 (day 7) | 1 | JY_intermediate_precision_2_1 |
| JY cell line | Precision | 2 (day 7) | 2 | JY_intermediate_precision_2_2 |
| JY cell line | Precision | 2 (day 7) | 3 | JY_intermediate_precision_2_3 |
| JY cell line | Precision | 3 (day 7) | 1 | JY_intermediate_precision_3_1 |
| JY cell line | Precision | 3 (day 7) | 2 | JY_intermediate_precision_3_2 |
| JY cell line | Precision | 3 (day 7) | 3 | JY_intermediate_precision_3_3 |
| JY cell line | LOD, specificity, accuracy | spiked 0.1 fmol synthetic peptides | 1 | JY_100amol_1_1 |
| JY cell line | LOD, specificity, accuracy | spiked 0.1 fmol synthetic peptides | 2 | JY_100amol_1_2 |
| JY cell line | LOD, specificity, accuracy | spiked 0.1 fmol synthetic peptides | 3 | JY_100amol_1_3 |
| JY cell line | LOD, specificity, accuracy | spiked 1 fmol synthetic peptides | 1 | JY_1fmol_1_1 |
| JY cell line | LOD, specificity, accuracy | spiked 1 fmol synthetic peptides | 2 | JY_1fmol_1_2 |
| JY cell line | LOD, specificity, accuracy | spiked 1 fmol synthetic peptides | 3 | JY_1fmol_1_3 |
| JY cell line | LOD, specificity, accuracy | spiked 10 fmol synthetic peptides | 1 | JY_10fmol_1_1 |
| JY cell line | LOD, specificity, accuracy | spiked 10 fmol synthetic peptides | 2 | JY_10fmol_1_2 |
| JY cell line | LOD, specificity, accuracy | spiked 10 fmol synthetic peptides | 3 | JY_10fmol_1_3 |
| JY cell line | LOD, specificity, accuracy | spiked 100 fmol synthetic peptides | 1 | JY_100fmol_1_1 |
| JY cell line | LOD, specificity, accuracy | spiked 100 fmol synthetic peptides | 2 | JY_100fmol_1_2 |
| JY cell line | LOD, specificity, accuracy | spiked 100 fmol synthetic peptides | 3 | JY_100fmol_1_3 |
| PBMC | Robustness | 1 | 1 |  |
| PBMC | Robustness | 1 | 2 |  |
| PBMC | Robustness | 1 | 3 |  |
| CLL | Robustness | 1 | 1 |  |
| CLL | Robustness | 1 | 2 |  |
| CLL | Robustness | 1 | 3 |  |
| BC | Robustness | 1 | 1 |  |
| BC | Robustness | 1 | 2 |  |
| BC | Robustness | 1 | 3 |  |
