## Supplemental information for "Validation of a high-performance liquid chromatography-tandem mass spectrometry immunopeptidomics assay for the identification of HLA class I ligands suitable for pharmaceutical therapies"

### Supplemental Table Legends

**Supplemental Table S1:** Selected individual synthetic peptides for the validation of the LC-MS/MS method. The 62 synthetic isotope labeled peptides consist of different amino acid sequences and physicochemical properties and depict a cross-section of peptides identified in a typical immunopeptidomics experiment. For the assessment of the differences between the peptides, the grand average of hydropathicity (GRAVY), the theoretical isoelectric point (pI) under physiological conditions, and the molar mass of the peptides were calculated using ProtParam (https://web.expasy.org/protparam). Heavy-labeled amino acids are indicated in bold (leucine +7 Da, proline and valine +6 Da). Abbreviations: pI, isoelectric point, GRAVY, grand average of hydropathicity.

**Supplemental Table S2:** Experimental design. Information about the sample, performed analysis, number of replicates, and raw file name of each sample is provided. Abbreviations: LOD, Limit of detection; PBMC, Peripheral blood mononuclear cell; CLL, Chronic lymphocytic leukemia; BC, Bladder cancer.

**Supplemental Table S3**: Limit of detection. List of identified peptides in four aliquots of HLA eluted ligands from JY cells spiked with 0.1 fmol, 1 fmol, 10 fmol, and 100 fmol isotope labeled synthetic peptides. The lowercased amino acids indicate the isotope labeled amino acids. Abbreviations: ΔM, mass deviation; PPM, parts per million; RT, retention time.

**Supplemental Table S4**: Specificity. Information about the precursor mass, the mass deviation from the theoretical mass, the retention time, and the selected top five product ions is provided. Abbreviations: ΔM, mass deviation; PPM, parts per million; RT, retention time.

**Supplemental Table S5**: Precision. List of identified peptides in nine separate analytical runs and further nine runs after one week. Abbreviations: PSM, peptide-to-spectrum matches; PEP, posterior error probability; Xcorr, cross correlation score, [illegible]M, mass deviation; PPM, parts per million; RT, retention time.

**Supplemental Table S6**: Specificity in primary patient samples. Information about the precursor mass, the mass deviation from the theoretical mass, the retention time, and the selected top five product ions is provided. Abbreviations: ΔM, mass deviation; PPM, parts per million; RT, retention time.

**Supplemental Table S7**: Specificity of the LTQ Orbitrap XL containing LC-MS/MS system. Information about the precursor mass, the mass deviation from the theoretical mass, the retention time, and the selected top five product ions is provided. Abbreviations: ΔM, mass deviation; PPM, parts per million; RT, retention time.

### Supplemental Figure Legends

**Supplemental Figure S1**: Validation of the specificity using the MS/MS spectra of the natural presented peptides with the corresponding synthetic peptides of the six selected peptides AIVDKVPSV, SPQGRVMTI, RPSGPGPEL, YLLPAIVHI, KVLEYVIKV, and SPSSILSTL.

**Supplemental Figure S2**: Validation of the specificity using the MS/MS spectra of the natural presented peptides with the corresponding synthetic peptides of AIVDKVPSV and YLLPAIVHI in CLL and PBMC samples and SLLQHLIGL in the BC sample.

**Supplemental Figure S3**: Investigation of the specificity of the LTQ Orbitrap XL LC-MS/MS system using the MS/MS spectra of the natural presented peptides with the corresponding synthetic peptides of the six selected peptides AIVDKVPSV, SPQGRVMTI, RPSGPGPEL, YLLPAIVHI, KVLEYVIKV, and SPSSILSTL.
